# Spinal neuron diversity scales exponentially with swim-to-limb transformation during frog metamorphosis

**DOI:** 10.1101/2024.09.20.614050

**Authors:** David Vijatovic, Florina Alexandra Toma, Zoe P. M. Harrington, Christopher Sommer, Robert Hauschild, Alexandra J. Trevisan, Phillip Chapman, Mara J. Julseth, Susan Brenner-Morton, Mariano I. Gabitto, Jeremy S. Dasen, Jay B. Bikoff, Lora B. Sweeney

## Abstract

Vertebrates exhibit a wide range of motor behaviors, ranging from swimming to complex limb-based movements. Here we take advantage of frog metamorphosis, which captures a swim-to-limb-based movement transformation during the development of a single organism, to explore changes in the underlying spinal circuits. We find that the tadpole spinal cord contains small and largely homogeneous populations of motor neurons (MNs) and V1 interneurons (V1s) at early escape swimming stages. These neuronal populations only modestly increase in number and subtype heterogeneity with the emergence of free swimming. In contrast, during frog metamorphosis and the emergence of limb movement, there is a dramatic expansion of MN and V1 interneuron number and transcriptional heterogeneity, culminating in cohorts of neurons that exhibit striking molecular similarity to mammalian motor circuits. CRISPR/Cas9-mediated gene disruption of the limb MN and V1 determinants FoxP1 and Engrailed-1, respectively, results in severe but selective deficits in tail and limb function. Our work thus demonstrates that neural diversity scales exponentially with increasing behavioral complexity and illustrates striking evolutionary conservation in the molecular organization and function of motor circuits across species.

## INTRODUCTION

In vertebrates, spinal neurons are the final neural relay for implementing the variable and dynamic patterns of muscle contraction that underlie diverse locomotor behavior. Agnathans, most bony fish, and larval amphibians move via alternating left-right contraction of axial musculature that propagates in the rostral to caudal direction, thereby producing undulatory swimming.^1^ Tetrapods, in contrast, move via both axial musculature and coordinated contraction of limb muscles organized along flexor-extensor, left-right and proximodistal axes.^2^ These different movement patterns are governed by networks of motor neurons (MNs) and interneurons in the spinal cord.^3,4^ While some core aspects of spinal circuit architecture are conserved across vertebrate species,^5^ whether and how the molecular diversity of spinal neurons varies between mammals and other vertebrates, and how neuron diversity relates to their distinct movement patterns remains to be investigated.

The African clawed frog, *Xenopus laevis,* undergoes a dramatic reorganization of motor output during metamorphosis, providing a unique opportunity to evaluate the contribution of spinal neurons to the changing movement pattern of a single organism over developmental time. Movement rapidly changes during the two-month transition from hatchling tadpole to juvenile froglet^6^. Beginning at Nieuwkoop and Faber (NF) stage 37-38 (NF37-38), tadpoles move via undulatory escape swimming. They then transform to free swimming at NF44, and finally to four-limbed movements (walking, hopping, and scooping) during metamorphosis (NF50-65). While the *Xenopus* hatchling escape circuit and its different motor outputs have been extensively studied using electrophysiology^7^, metamorphosing *Xenopus* motor circuits have not. How this metamorphic transformation unfolds, the extent to which motor and spinal interneuron identities change, and the role of interneurons in driving this behavioral transition remain largely unknown.

Comparative analysis of vertebrate spinal circuits suggests that the heterogeneity of inhibitory and excitatory interneurons that modulate MN activity may vary according to their motor output^5^. In simple escape swimming circuits, exemplified by those of the hatchling tadpole, five anatomically defined interneuron (IN) types—ascending aIN, commissural cIN, and descending dIN, dIc and dIa—are sufficient to generate alternating left-right patterns of undulatory tail movement.^7^ More complex swimming circuits, such as those of zebrafish, consist of at least three different MN types and interneuron types.^8^ Spinal neuron diversity appears to reach its apogee in limbed vertebrates, such as the mouse, in which 12 cardinal classes of spinal neurons arise from distinct progenitor domains along the dorsoventral axis, each expressing a unique transcriptional code^9^ and further classified by their physiology, projection patterns, and neurotransmitter expression.^10^

In mice, individual cardinal classes exhibit extensive subtype heterogeneity,^11^ as highlighted by studies on MN,^12,13^ V1,^14–17^ V2a,^18,19^ and V3^20–22^ interneuron diversity. MNs are organized into four columnar classes: (*i*) the lateral motor column (LMC), innervating limb muscles; (*ii*) the hypaxial motor column (HMC), innervating hypaxial muscles; (*iii*) the preganglionic column (PGC), innervating sympathetic ganglia; and (*iv*) the medial motor column (MMC), innervating axial muscles.^12^ These motor columns are further subdivided along the rostrocaudal, dorsoventral and mediolateral axes into divisions and motor pools innervating individual muscles.^12,23^ Spinal V1 interneurons, essential for both swim and limb-based movement, provide over 50% of inhibitory inputs onto MNs^24^ and can be classified into at least four clades and over 50 predicted cell types based on the combinatorial expression of 19 transcription factors and their cell body positions in the spinal cord.^14,15^ V1 subtypes further vary in their transcriptional profiles within limb and non-limb spinal segments.^16^ To what extent this neuronal architecture is conserved in amphibians, such as frogs, and how it scales during the swim-to-limb transition of metamorphosis, remains to be explored.

Here, we perform a detailed analysis of locomotor behavior across *Xenopus* frog metamorphosis and relate swim versus limb movement patterns to the molecular and cellular architecture of the spinal cord. We observe a large increase in number and diversity of motor and V1 inhibitory neurons that parallels the diversification of movement. MNs in the early swim circuit have a uniform MMC subtype identity, whereas during metamorphosis, their number and diversity expand to form the same four molecular and anatomical columns observed in the mouse. V1 inhibitory interneurons also expand and diversify during metamorphosis. At the climax of limb circuit development, the same individual transcription factors that define V1 subpopulations in mice are conserved in both proportion and position in the frog. Differences between species are only observed at the level of V1 subtypes, defined by the co-expression of two or three transcription factors. Lastly, we demonstrate that disruption of motor and V1 interneuron development via CRISPR-based loss-of-function of FoxP1 and Engrailed-1 (En1) respectively, leads to loss of molecular and behavioral complexity in frogs, recapitulating similar loss-of-function experiments in mice. We thus show that MN and V1 interneuron heterogeneity scales with the metamorphic transition from swimming to limb movement, culminating in a remarkably similar degree of neuronal diversity between frogs and mice, species that last shared a common ancestor nearly 360 million years ago.

## RESULTS

### Swim-to-limb transition in movement during frog metamorphosis

To characterize how movement changes during tadpole-to-frog metamorphosis, we developed a quantitative behavioral assay to measure locomotion in freely moving *Xenopus laevis* from larval NF37-38, the peak of escape swimming,^25,26^ to juvenile frog stages, the end of metamorphosis^6^ (**Figure 1A-G**). Animals were video-tracked and videos subjected to pose estimation by the deep-learning framework SLEAP^27,28^ (**Figure S1A-D**). Metamorphic stages were split into seven bins according to anatomical features, such as the emergence of hindlimb locomotion at NF57-58 and forelimb at NF59-62^6^ (**Figure 1D-E**). For each bin, two SLEAP models were trained: a centroid model to detect the animal’s center point (**Figure S1C)** and a centered model to track the position of the limbs and tail (**Figure S1D)**. All models achieved high levels of tracking precision, as confirmed by quality control metrics (**STAR Methods**).

**Figure 1.**
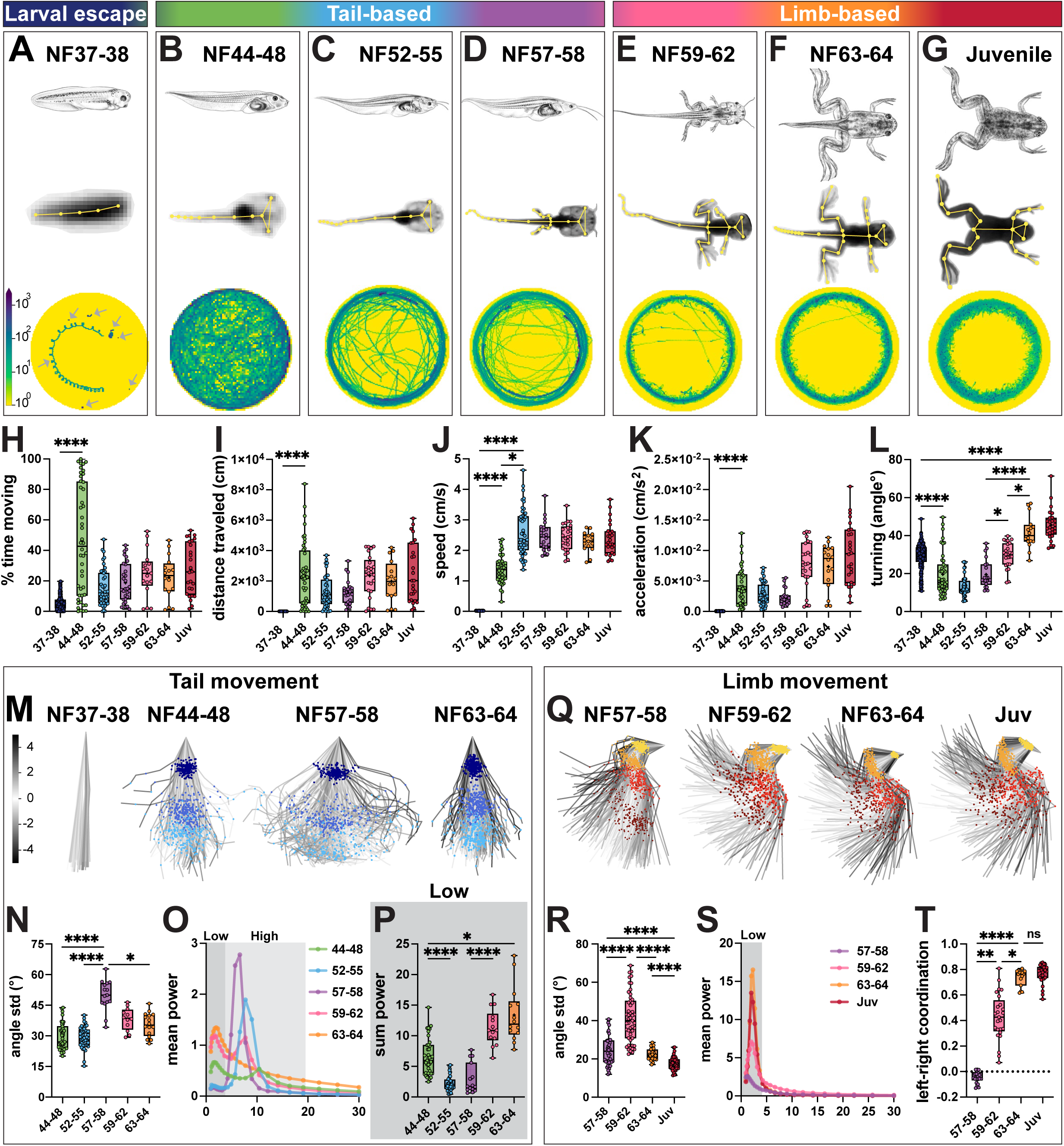
Loss of tail and emergence of limb movement during frog metamorphosis. A-G. Larval escape, tail-based and limb-based locomotion during *Xenopus laevis* metamorphosis. Stages across frog metamorphosis are divided into seven bins according to their anatomical features (Nieuwkopp and Faber; NF^116^): NF37-38 (**A**; dark blue), NF44-48 (**B**; green), NF52-55 (**C**; blue), NF57-58 (**D**; purple), NF59-62 (**E**; pink) NF63-64 (**F**; orange) and juvenile (**G**; red). We further grouped stages in larval escape (**A**), tail-based (**B-D**) and limb-based (**E-G**). For each stage, a schematic of tadpole anatomy (**top row**, adapted from Xenopus illustrations © Natalya Zahn, 2022^117^); a SLEAP skeleton (yellow) superimposed onto an image of a recorded animal with all tracked points indicated (**A-G**, **middle row**); and an example of the distance traveled by 7 animals for NF37-38 (**A**, **bottom row**, arrows) or by a single animal from NF44-48 to juvenile stage (**B-G**, **bottom row**) are shown. Trajectories of the distance traveled show distinct patterns at each stage: coiling escape swimming (**A**; animals are mostly stationary), free-feeding exploration of the whole dish (**B**) with a transition to edge tracking from NF44-48 (**B**) to juvenile (**G**) stage. **H-L. Quantification of tadpole and frog movement.** The percentage of time spent moving (**H**) and the length of the distance traveled (**I**) per one-hour imaging session, increase from NF37-38 to NF44-48 and then stay constant for successive stages (**H-I**; NF37-38 versus NF44-48, p = <0.0001). Mean speed increases stepwise from NF37-38 to NF52-55, and then remains constant (**J**; for NF37-38 versus NF44-48 and NF37-38 versus NF52-55, p = <0.0001; NF44-48 versus NF52-55, p = 0.019). Acceleration increases from NF37-38 to NF44-48 and then remains constant (**K**, NF37-38 versus NF44-48, p = <0.0001). Turning, calculated as the mean directional change of the body-part trajectory every 8th frame, decreases from NF37-38 to NF44-48, then increases from NF57-58 to juvenile stage (**L**; 0 degree angle is parallel indicating no turning; for NF37-38 versus NF44-48, NF37-38 versus juvenile and NF57-58 versus NF63-64, p = <0.0001; for NF57-58 versus NF59-62 and NF59-62 versus NF63-64, p = 0.02). **M-P. Range and frequency of tail movement change across tadpole metamorphosis.** PCA plots represent the position of the tail and its range of movement during 256 random frames (**M**; tail top, dark blue; tail mid, blue; tail tip, light blue). The first visible increase in tail movement is from NF37-38 to NF44-48 and the second from NF44-48 to NF57-58, then the range decreases from NF57-58 to NF63-64 (**M**). Quantification of the range of movement at the tail tip shows a peak at NF57-58 and a decrease from NF59-64 as the tail recedes (**N**; for NF44-48 versus NF57-58 and NF52-55 versus NF57-58, p = <0.0001; NF57-58 versus NF63-64, p = 0.048). Mean power spectrum of frequency of tail tip oscillations for each stage of metamorphosis, with low (0.9-4.5 Hz, dark gray) and high (4.5-20 Hz, light gray) frequency bins highlighted (**O**). From NF44-48 to NF59-62, the frequency spectrum is bimodal with a peak in the low and high frequency bins; at NF63-64, it is unimodal with only one low frequency peak (**O**). The amount of tail tip movement in the low frequency bin, represented by the sum power, decreases from NF44-48 to NF52-55, and then increases until NF63-64 (**P**; for NF44-48 versus NF52-55 and NF57-58 versus NF59-62, p = <0.0001; NF44-48 versus NF63-64, p = 0.02). **Q-T. Gain of hindlimb movement during frog metamorphosis.** PCA plots represent the position of the hindlimb and its range of movement during 256 random frames showing an increase in range from NF57-58 to NF59-62 (**Q**; hip, yellow; knee, orange; ankle, red; foot, brown). Quantification of the range of knee movement shows an initial increase from NF57-58 to NF59-62 and then a decrease until juvenile stage (**R**; for NF57-58 versus NF59-62, NF57-58 versus juvenile, NF59-62 versus NF63-64, and NF63-64 versus juvenile, p = <0.0001). Mean power spectrum of the knee oscillations for each stage of metamorphosis shows a single peak in the low frequency range (**S**; 0.9-4.5 Hz, dark gray). Coordination of the left and right knees changes from random at NF57-58 to bilaterally synchronous at NF63-64 (**T**; +1 = synchronous, 0 = random, −1 = alternating; NF57-58 versus NF59-62, p = 0.003; NF57-58 versus NF63-64, p = <0.0001; NF59-62 versus NF63-64, p = 0.02). n = 172 animals for NF37-38; n = 47 animals for NF44-48; n = 24 animals for NF52-55, n = 11 animals for NF57-58, n = 13 animals for NF59-62, n = 8 animals for NF63-64, n = 13 animals for juvenile stage. Scale bar in **A** indicates the number of times the animal was present in a specific area of the dish from no time (10^0^ frames, yellow) to many times (10^3^ frames, blue). Scale bar in **M** indicates the color-code of the first principal component of variation of the aligned tail and limb positions in **M** and **Q**.

We first compared general features of tadpole and frog movement across all developmental stages (**Figure 1H-L**). NF37-38 larval tadpoles were largely stationary, moving only short distances at a low speed and acceleration (**Figure 1A, H-K**). Their movement was characterized by stereotyped corkscrew spiraling with a high amount of turning (**Figure 1L; Movie S1**), consistent with previous observations.^26^ As tadpoles transitioned to free-swimming at NF44-48, they increased their moving time by 9-fold and covered 600x longer distances, with higher mean speed and acceleration but decreased turning (**Figure 1B, H-L**). This high level of movement persisted until later stages (**Figure 1C-I; Movie S2-S7**), with speed and acceleration remaining stable across development, whereas turning further increased beginning at NF59-62 (**Figure 1J-L**). At later limb stages, tadpoles changed from exploring the whole dish to preferentially tracking its edge (**Figure 1B v. 1E-G**), a behavior observed across species^29^ with a non-visual sensory basis in tadpoles.^30,31^ Based on these observations, metamorphosis can be subdivided into three general phases of motor behavior, characterized by increasing propulsion, precision and adaptability: larval escape swim (NF37-38, **Figure 1A**), tail-based swim (NF44-48 to NF57-58; **Figure 1B-D**), and limb-based locomotion (NF59-62 to juvenile stages, **Figure 1E-G**).

We next performed kinematic analysis of tail, hindlimb and forelimb joint movement across developmental stages. Principal component analysis (PCA) revealed an initial increase in the range of tail movement from larval to free-swimming stages, peaking during tail-based swim stages and subsequently decreasing at limb-based stages (**Figure 1M-N)**. The frequency of tail oscillation also correspondingly changed from tail-to limb-based stages (**Figure 1O**): we observed two dominant frequency peaks, one at a low frequency of ∼1.5-2 Hz and another at a high frequency of ∼6-10 Hz (**Figure S1E,G**), consistent with previous estimates.^32^ At metamorphosis, tail movement progressively transitioned from proportional low- and high-frequency oscillations, to majority high-frequency oscillations, and finally, to majority low-frequency oscillations (**Figure 1O-P; Figure S1E-G**). The first transition to high-frequency tail movement correlated with an increase in locomotor speed, while the second transition from high- to low-frequency oscillatory behavior correlated with the emergence of limb movement (**Figure 1J,O-T**). Accordingly, at the final stage of tail recession, the tail no longer exhibited a bimodal frequency distribution and instead showed only low-frequency movement at 2.2 Hz (**Figure 1O; Figure S1E,G**) that aligned with distribution and frequency of limb movement (**Figure S2D, I vs. O)**.

To explore the differential contribution of each tail region to movement, we analyzed the range and frequency at each point along the tail’s rostrocaudal axis. The range of tail movement was similar across the tail at the free-swimming stage, but as metamorphosis proceeded, we observed an increase in localized movement at the tail tip, followed by a reduction in the range and synchrony of movement between tail segments as it receded (**Figure S2A-B**). Frequency analysis along the rostrocaudal axis showed that while the tail tip and midpoint followed the same pattern of oscillations, the tail top–the most rostral point–always displayed higher power, suggesting it functions as the initial hub of tail oscillation (**Figure S2E-I**). The late-stage matching of tail and limb frequencies was also most pronounced at the tail top (**Figure S2D**), the region closest to the limbs.

In parallel, limb movement also developed, revealing three main kinematic features. First, the limbs began to move during metamorphosis, with the angle and range of knee and ankle increasing from NF57-58 to NF59-62 (**Figure 1Q-R; Figure S1H; Figure S2J-K**), precisely at the timing of their transition from passive to active movement.^33,34^ However, at the hip and foot, this initial increase in range was not observed (**Figure S2K**). Moreover, for all hindlimb joints, as well as for the elbow, the overall range of movement decreased after NF59-62 (**Figure 1RQ-R; Figure S1K-M; Figure S2K**). The end result of metamorphosis is thus a decrease in the range of movement and a change in its form, with more localized movements emerging at later stages. Second, synchrony of limb movement increased in a stepwise manner at all joints (**Figure 1T; Figure S2L**), consistent with emergence of bilateral kicking, the dominant form of movement in *Xenopus laevis*^35^ and many other frogs.^36^ Synchrony of the foot was the lowest in magnitude and the last to develop (**Figure S2L**), showing a delay in proximodistal limb coordination. Third, the frequency of hind- and forelimb joint oscillations displayed a single peak at low frequency which gradually increased in power over metamorphosis (**Figure 1S; Figure S1I,N-O; Figure S2M-Q**). The dominant frequency of limb movement initially matched the tail’s frequency of ∼1.5 Hz (**Figure S2C**), and then shifted to ∼2.2 Hz at NF63-64 (**Figure S2D**), mirroring that of mouse hindlimb stepping.^37^ Both the power of the frequency of limb movement and its dominant frequency reached their final state at NF63-64 (**Figure S1I-J; Figure S2M-Q**) demonstrating limb circuits are fully formed before the tail fully recedes.

In summary, our kinematic quantification makes several novel observations about how motor behavior changes during tadpole-to-frog metamorphosis. First, we define distinct larval escape, tail-based and limb-based locomotor phases, each characterized by unique locomotor profiles. Second, we observe not only a loss of tail coordination and gain of limb coordination over metamorphosis but also identify rhythmic coupling between the tail and limbs at intermediate stages. This analysis sets the stage for dissecting the cellular components within the spinal cord that generate each locomotor pattern.

### Motor neurons expand and diversify during frog metamorphosis

The observed transformation of motor behavior during frog metamorphosis necessitates the addition of new muscle groups and corresponding neural circuits to regulate their activity. Motor neurons (MNs), which segregate into well-described molecularly, anatomically and functionally distinct columns, divisions, and pools in mammals,^12^ provide a starting point to evaluate how movement is generated and controlled in the tadpole and frog.

To evaluate whether the mouse MN subtype architecture was conserved in frogs, we used immunohistochemistry to profile the number and type of MNs in larval escape swim (NF35-38), free-swim (NF44-47), and limb (NF54-55) spinal circuits. At all developmental stages, a ventral population of spinal neurons co-expressed the MN transcriptional determinants, Isl1 and Mnx1/Hb9^38–41^ **(Figure 2A-D; Figure S3A-B**). At early larval swim stages, there were ∼5 Isl1/2/Hb9-expressing MNs per 15 µm section (**Figure 2A, D**), consistent with previous estimates from retrograde labeling experiments.^42^ As larval tadpoles transitioned to free swimming at NF44, the number of MNs doubled at all spinal cord levels (**Figure 2B, D; Figure S3A-B**). This increase became even more pronounced during metamorphosis, with a 10-fold expansion at limb and 5-fold expansion at thoracic levels by NF54-55 (**Figure 2C-D**), a stage characterized by peak MN number (**Figure S4A-B, E-G**) and limb development.^6^

**Figure 2.**
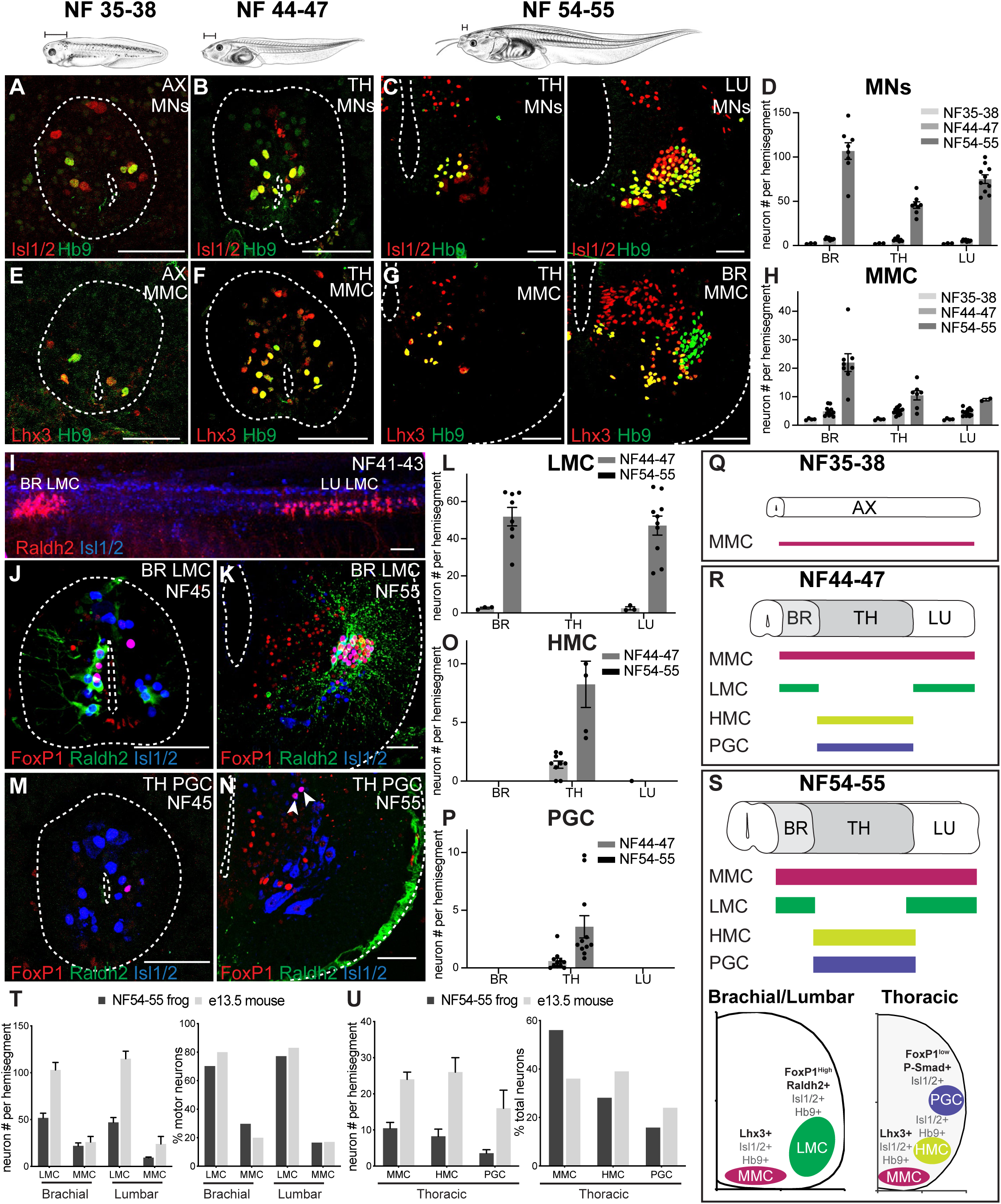
Spinal motor neurons expand and diversify during *Xenopus* frog metamorphosis. A-D. Motor neuron number increases during metamorphosis. Motor neurons in the axial (AX), thoracic (TH), and lumbar (LU) spinal cord express the pan-motor neuron markers Hb9 (green) and Isl1/2 (red) and increase in number between NF35-38 (**A**), NF44-47 (**B**) and NF54-55 (**C**). Bar graph (**D**) shows the total number of Hb9^+^ Isl1/2^+^ (NF37-38 and NF44-47) or ventral Isl1/2^+^ (NF54-55) motor neurons per 15 μm ventral horn (mean ± SEM for n = 3–10 animals) of brachial, thoracic, and lumbar spinal cord. **E-H. Medial motor column (MMC) emerges in larval *Xenopus* and expands during metamorphosis.** The antibody against the Lhx3 transcriptional determinant of MMC identity (red) labels a subset of Hb9^+^ motor neurons (green) in the spinal cord of NF35-38 (**E**), NF44-47 (**F**), and NF54-55 (**G**) tadpoles. Shown are axial (AX; NF35-38), thoracic (TH; NF44-47 and NF54-55) and brachial (BR; NF54-55) sections (see **Figure S3** for the images of additional immuno-labeled TH, BR and LU sections). The graph (**H**) shows the number of Lhx3^+^ Hb9^+^ MMC motor neurons per 15 μm ventral horn (mean ± SEM for n = 4–14 animals) at brachial, thoracic and lumbar levels at NF35-38, NF44-47, NF54-55. **I-P. Lateral (LMC), preganglionic (PGC) and hypaxial (HMC) motor columns emerge in free-swimming tadpoles and expand during metamorphosis.** Side view of an NF41-43 spinal cord (**I**) stained for the LMC determinant, Raldh2 (red), and Isl1/2 (blue) reveals the nascent brachial and lumbar populations of limb-innervating motor neurons. The LMC determinants, FoxP1 (red) and Raldh2 (green), jointly label a motor neuron subset (Isl1/2, blue) at brachial and lumbar levels at NF44-47 (**J**) and NF54-55 (**K**). The number of LMC motor neurons (**L**) per 15 μm ventral horn (mean ± SEM for n = 3–10 animals) of brachial, thoracic, and lumbar segments at NF44-47 and NF54-55. Frog PGC motor neurons labeled by the PGC transcriptional determinants, FoxP1 (red) and Isl1/2 (blue), at thoracic levels at NF44-47 (**M**) and 54-55 (**N**). Neither LMC nor PGC is present at NF35-38 (see **Figure S3G-J**). The number of HMC (**O**) and PGC (**P**) motor neurons per 15 μm ventral horn (mean ± SEM for n = 5–11 animals) of brachial, thoracic, and lumbar levels at NF44-47 and NF54-55. HMC motor neuron number at the thoracic level was calculated by subtracting the number of MMC and PGC motor neurons from the total number of motor neurons. **T-U. Conservation of MN proportions in developing limb circuits of frogs and mice.** Number of LMC and MMC motor neurons (left) and their percentage (right) in the total motor neuron population at brachial and lumbar levels (**T**) at frog NF54-55 and E13.5 mouse. Number of HMC, PGC, and MMC motor neurons (left) and their percentage (right) in the total motor neuron population at the thoracic level (**U**) at frog stage NF54-55 and E13.5 mouse. Shown is the mean ± SEM for n = 5–11 animals. The embryonic mouse counts were extracted from Agalliu *et al*, 2009 with the SEM estimated from the provided plots. **Q-S. Summary of motor neuron development in tadpoles.** Schematics showing the rostrocaudal distribution of MMC (Lhx3^+^, Isl1/2^+^ and Hb9^+^), HMC (Isl1/2^+^ and Hb9^+^) and PGC (FoxP1^low^, Raldh2^+^, Lhx3^+^, Isl1/2^+^ and Hb9^+^) subsets in NF35-38 (**Q**), 44-47 (**R**), and 54-55 (**S, top**) spinal cords. Schematized spinal cord hemi-section of a limb and thoracic segment depicting the relative position and molecular markers of each motor column along the dorsoventral, mediolateral axis (**S, bottom**). Shown is either a spinal cord cross section (NF35-38/44-47) or hemi-section (NF54-55) with the central canal and outer edge indicated (dotted line). Scale bar, 50 μm (except in **I**, 100μm). Tadpole drawings adapted from Xenopus illustrations © Natalya Zahn (2022).^117^

We next probed the identity of the MNs that are added during metamorphosis. In the mouse, MN columns, divisions and pools are distinguished by transcription factor expression and axonal projection pattern.^43^ Those in the medial motor column (MMC) express Lhx3 and Lhx4 and project to the axial musculature lining the vertebral column.^44,45^ Consistent with an MMC identity, early larval MNs uniformly co-expressed Hb9 and Lhx3 (**Figure 2E**). At free-swimming stages, this MMC population doubled in number at all spinal levels (**Figure 2F, H)**. As tadpoles metamorphosed, MMC MNs further expanded in number with a specific enrichment at brachial levels (**Figure 2G-H**), paralleling the increase and remodeling of axial muscles.^46^

MNs in the mouse further segregate into segmental populations, with the lateral motor column (LMC) innervating limb muscles, the preganglionic motor column (PGC) that innervating the sympathetic chain ganglion, and the hypaxial motor column (HMC) that innervating the hypaxial musculature of the ventral torso.^12^ As in mice, we found FoxP1-positive and Raldh2/Aldh1a2-positive MNs located in the ventrolateral spinal cord at brachial and lumbar but not thoracic levels in frogs (**Figure 2K-L**), consistent with an LMC identity^47^ and confirming previous observations.^48^ As in amniotes,^43,49^ several conserved molecular features were observed in the frog: the expression of Isl1 and Hb9 subdivided the LMC into a medial and lateral division (**Figure S3C-D)**; a subpopulation of MNs expressed FoxP1 but not Raldh2 resembling the digit MNs^50^; and the transcription factor cScip/Pou3f1 labeled a small group of limb level MNs as has been shown for the flexor carpi ulnaris motor pool in mice^51^ (**Figure S3E**).

Extending our analysis to earlier tadpole development, we found that there were no neurons of the LMC subtype, defined by FoxP1, Raldh2, and Isl1 co-expression, at the larval swim stage NF35-38 (**Figure S3G-H**). Surprisingly, at free-swimming stages, we identified two small populations of LMC-like neurons—one rostral and one caudal—that express FoxP1 and Raldh2, and project to the limb bud, which we termed the pioneering LMC (**Figure 2I-J, L**). Using FoxP1, pSmad, and Isl1 co-expression markers,^47^ we also found PGC-like neurons in the pioneering thoracic region of free-swimming tadpoles that further expanded during metamorphosis forming a mediolateral cluster in the thoracic spinal cord (**Figure 2M-P; Figure S3B,F**). This time course of emergence of PGC parallels the formation of organs.^52^ Finally, we scored the development of the hypaxial motor column (HMC), defined by Isl1, Hb9 and the absence of all other columnar identifiers.^47^ The HMC followed a similar temporal and spatial pattern to that of the PGC—with the HMC also restricted to thoracic levels and doubling in size at metamorphosis. These data support high molecular conservation of MNs at both the population and subtype level in the frog (**Figure 2Q-S**) and reveal that expansion of MN number and subtype diversity is a hallmark feature of the frog tail-to-limb transformation.

### Comparing motor neuron diversity in frog and mouse

We next compared the molecular development of motor neurons in mouse and frog, identifying analogous stages of limb-circuit development based on the time course of spinal neuron and limb development (**Figure S4A-K**). We found that tadpoles had approximately half the number of limb and thoracic MNs per column as mice (**Figure 2T-U**). However, across spinal cord levels, the proportion of each subtype was largely conserved. Limb-level MNs comprised ∼70-80% LMC and ∼20% MMC in both species, while thoracic-level MNs subdivided into ∼35-55%

MMC, ∼30-40% HMC, and ∼15-25% PGC (**Figure 2T-U**, bottom). At larval escape swim stages, however, MNs were uniformly of the MMC type (**Figure S3H**) and, at free-swimming stages, 60% of MNs were MMC and 40% were either LMC or a combination of HMC/PGC by level (**Figure S3I-J**). These data support a transition from a distinct swimming to a conserved mouse-like MN cell type architecture during frog metamorphosis.

### Linking the molecular identity of motor neurons to their anatomical projection pattern

To assay whether molecularly conserved *Xenopus* MN subtypes exhibited similar anatomical projections as in mice, we generated a MN-specific transgenic line in which the murine microRNA-218 enhancer element^53^ drives fluorophore expression, termed *Xla.Tg(218-2:GFP)^Swee^*; **Figure 3A-I**). Molecular characterization of *218-2:GFP* showed that GFP expression at the larval tadpole stage was restricted to MNs, co-labeled by Isl1/2 (**Figure 3G**). Later, at NF45, *218-2:GFP* marked both the newly formed pioneering LMC, co-labeled by Raldh2, and the MMC, distinguished by its distinct ventromedial position and expression of Isl1/2 but not Raldh2 (**Figure 3H**). At NF50-52, after LMC MN addition (**Figure S4B, G**), *218-2:GFP* also was expressed in both LMC and MMC MNs, identified by Isl1/2 and the presence or absence of FoxP1, respectively (**Figure 3I**).

**Figure 3.**
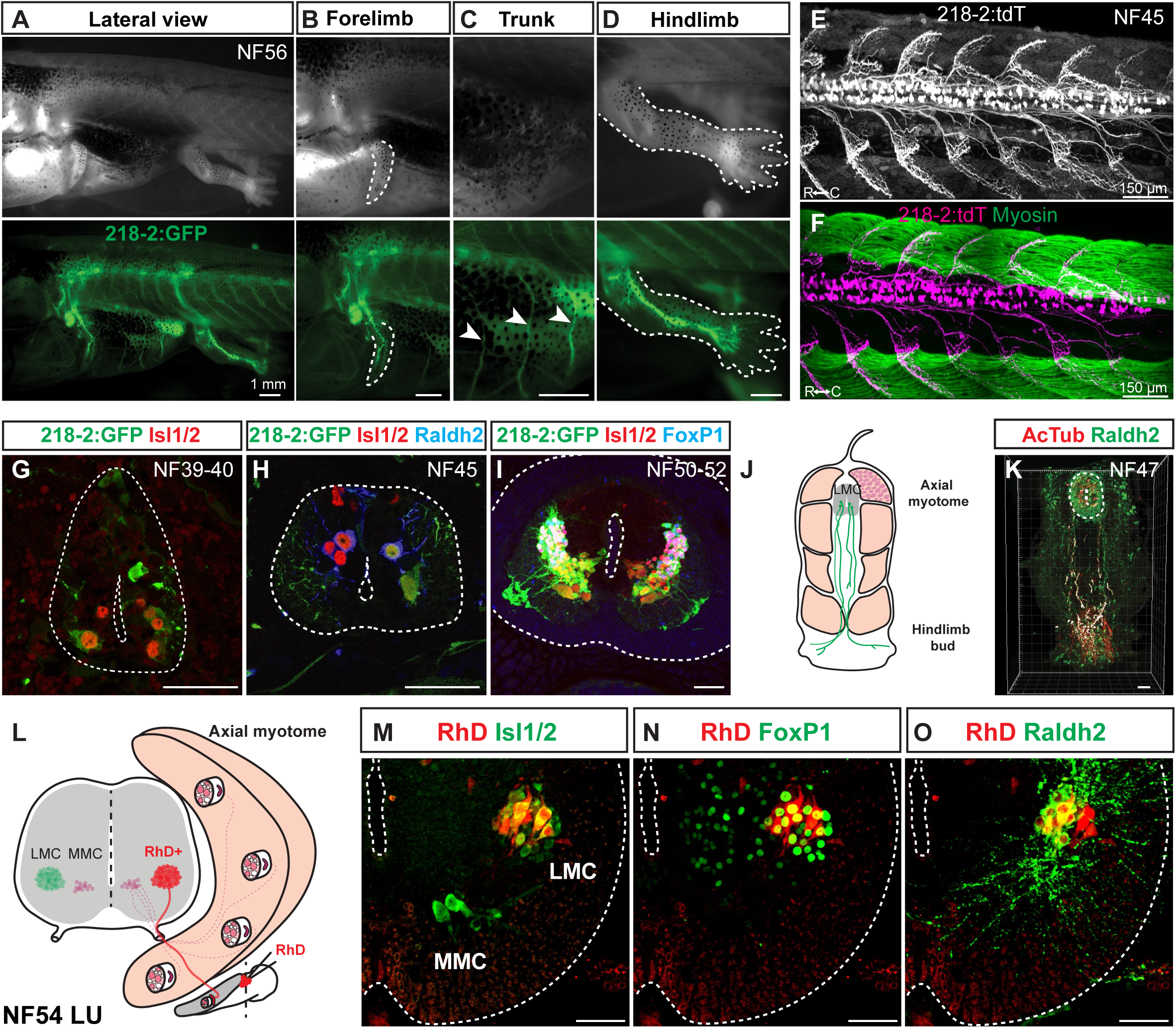
Linking motor neuron molecular profiles to anatomical projection pattern during frog metamorphosis. **A-D** GFP-labeled motor neurons in a limbed NF56 *218-2:GFP* tadpole (**A**) innervate the forelimb (**B**), trunk (**C**), or hindlimb (**D**), in line with the expected innervation patterns of the molecular populations detected at this stage. Scale bar, 1 mm. **E-F.** Motor neurons in free-swimming NF45 *218-2:GFP:tdT* tadpoles are distributed throughout the spinal cord and extend axonal arbors into the myotomal cleft, as shown by myosin heavy chain immunohistochemistry (green; **F**). Scale bar, 150 µm. **G-I.** GFP-labeled motor neurons at the limb levels in *218-2:GFP* tadpoles at NF39-40 (**G**), NF45 (**H**) and NF50-54 (**I**) express the pan-motor neuron marker Isl1/2 (red). At NF45 and at NF50-52, a Raldh2/FoxP1-positive population (blue) of motor neurons, consistent with their LMC identity, also expresses GFP. Scale bar, 50 µm. **J-K.** A pioneering population of LMC motor neurons at free-swimming stage NF47 expresses the LMC marker Raldh2 (green) and their axons, marked with the acetylated-α-Tubulin antibody (red), extend to the developing limb area prior to limb bud emergence. Scale bar, 50 µm. **L.** Schematic of medial and lateral motor column innervation patterns and the location of the rhodamine dextran application. Scale bar, 50 µm. **M-O.** Retrograde labeling with rhodamine dextran (RhD, red) marks a population of laterally positioned motor neurons that co-express MN (Isl1/2, green; **M**) and LMC (FoxP1/Raldh2, red; **N**/**O**) markers. Scale bar, 50 µm.

Given the selectivity of *218-2:GFP* for MNs across tadpole development, we utilized it to characterize the axonal projection pattern of LMC, HMC, and MMC to muscles. At NF56, GFP-positive axons exited the brachial/lumbar enlargements and innervated the forelimb and hindlimb (**Figure 3B, D**). Clear innervation of trunk muscles by *218-2:GFP*-positive axons was also observed at thoracic levels (**Figure 3C**). In addition, at the free-swimming stage, *218-2:GFP*-positive MNs projected to the myotomal cleft of the tail musculature (**Figure 3E-F)**. Taken together, this confirms *218-2:GFP* labels all MN types and muscle projections.

To examine if muscle projections were already established for the newly identified pioneering LMC at swim stages, we evaluated co-expression of Raldh2 and acetylated tubulin in MN axons at NF47. Co-labeled pioneering LMC axonal projections extended from the spinal cord to the hindlimb bud (**Figure 3J-K**), pioneering the innervation of the limb before metamorphosis. Finally, as an independent confirmation of LMC-to-limb connectivity, we administered fluorescently labeled dextran into the proximal limb at NF54 and observed colocalization of dextran, the MN marker Isl1, and the LMC markers FoxP1 and Raldh2 in ventral LMC MNs within the spinal cord (**Figure 3L-O**). The molecularly characterized MN subtypes in the frog spinal cord thus exhibited similar projection patterns as equivalent MN subtypes in the mouse.^47^

### V1 inhibitory neurons increase in number during frog metamorphosis

This high level of MN conservation raised the question of whether interneurons in the spinal motor system are also conserved between frogs and mice. Larval spinal circuits contain two inhibitory and four excitatory interneuron types based on their electrophysiological and anatomical properties.^54^ Amongst these, only aINs have been molecularly characterized and analogized to the V1 interneuron cardinal class based on the expression of the homeobox gene Engrailed-1.^55^ However, at later developmental stages, our knowledge of the molecular, anatomical or functional properties of frog interneurons is limited.

We thus aimed to characterize spinal interneurons in the frog by focusing on V1 interneurons (V1s). This diverse population of ventral inhibitory neurons directly modulates MN firing and influences locomotor speed^56–58^ and flexor-extensor alternation,^59,60,24,61^ both of which vary over the metamorphic transition from tadpole to frog (**Figure 1**). We identified a ventral population of neurons that expressed En1 (**Figure 4A-D**), a well-established V1 marker across species,^62,63^ and assessed how the number of V1s scaled with MN number from swim to limb-circuit stages. At escape swimming stages, we observed about ∼1 V1 interneuron for every ∼2.5 MNs per 15 μm spinal tissue hemisection (**Figure 4A, E**). As tadpoles transitioned to free swimming and began to regulate their speed and turn frequency (NF44-47; **Figure 1**), V1 number increased to ∼2 and MN number to ∼5.5 cells (**Figure 4B, E**). Finally, with the emergence of MN subtype diversity and coordinated limb movement during metamorphosis, V1 number peaked at ∼40 in thoracic and ∼72 in the lumbar hemi-sections (**Figure 4C-D**), a ∼70-fold expansion in cell number.

**Figure 4.**
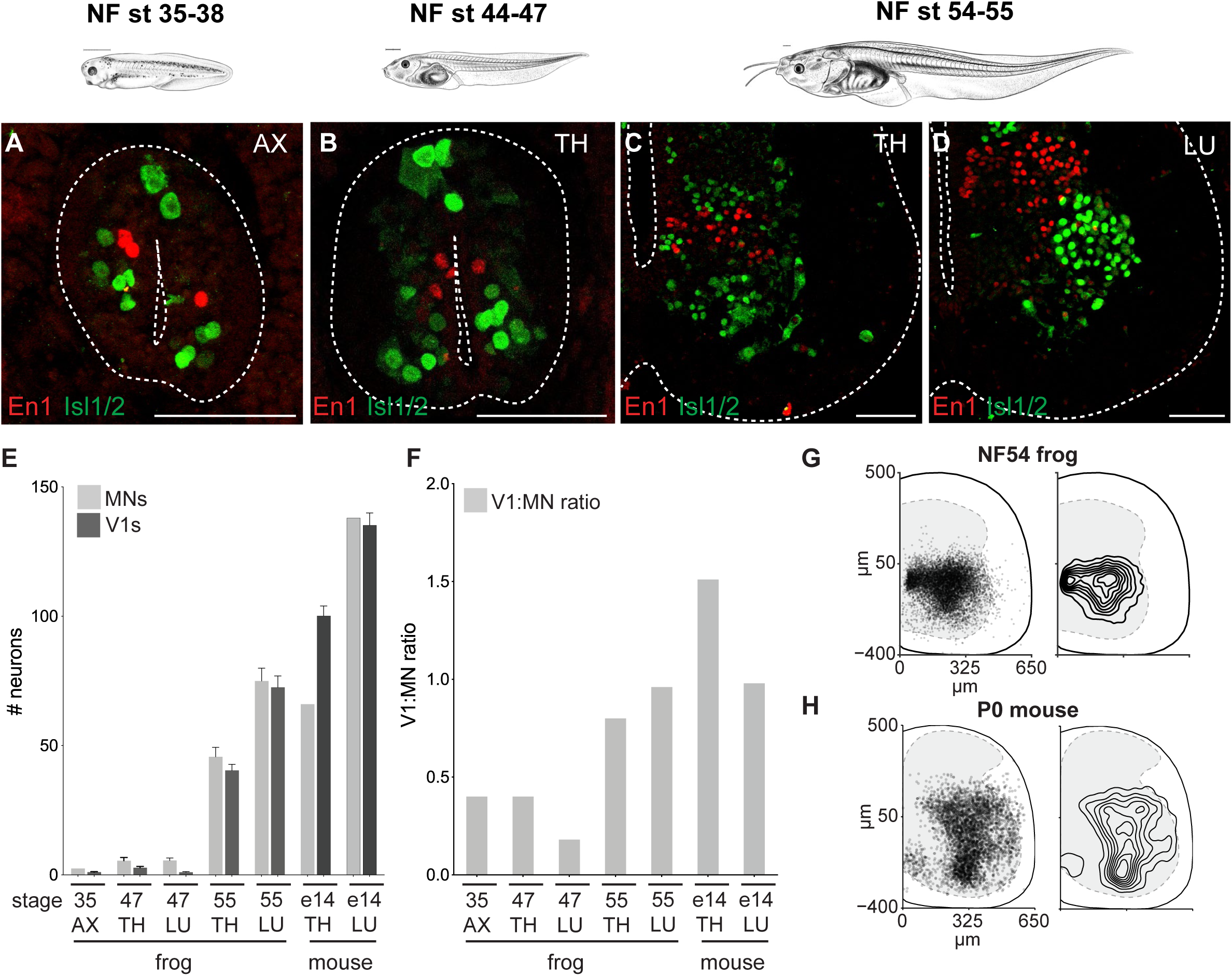
V1 inhibitory interneurons increase in number with metamorphic expansion of motor neurons. **A-D** Immunoreactivity against the Engrailed1 (red), a V1 inhibitory interneuron (V1) marker, and Isl1/2 (green), a motor neuron (MN) marker, labels ∼1 V1 and ∼2.5 MNs at NF35-38 (**A**), ∼2 V1s and ∼5.5 MNs at the thoracic levels at NF44-47 (**B**), and around ∼40 and ∼45 V1s and MNs at the thoracic levels (**C**)and ∼72 and ∼75 V1 and MNs at the lumbar levels at NF54-55 (**D**), respectively. Tadpole drawings adapted from Xenopus illustrations © Natalya Zahn (2022).^117^ **E-F.** Number (**E**) and the ratio (**F**) of V1s and MNs at axial (NF35), thoracic and lumbar (NF47 and 55 tadpole and E14 mouse) levels. At NF35 and NF47, the V1:MN ratio is under 0.5, and then approaches 1 at NF54-55 for both thoracic and lumbar segments in metamorphosing frogs, similar to in the embryonic mouse. Shown in **E** is the mean ± SEM for n = 4–17 animals per 15 μm ventral horn. **G-H.** Position of V1 interneurons at lumbar levels of NF54 frog (**G**) and P0 mouse (**H**). Plotted on the left are individual cells with 50% transparent black to highlight overlap.

This expansion led us to evaluate how the ratio between V1s and MNs, a numerical proxy for interneuron regulation, varied across metamorphosis. We found that the V1-MN ratio increased from ∼0.5:1 to ∼1:1 as tadpoles transitioned from swimming to limb movement (**Figure 4F**), indicating a preferential expansion of interneurons relative to MNs. The resulting 1:1 V1-to-MN ratio mirrored that of mice at comparable developmental stages^14,16,64^ and thus emerged as a conserved feature of developing tetrapod limb circuits.

We additionally compared the settling position of V1s between frogs and mice, as previous reports correlate settling position with functional properties.^14,17,60^ As in mice,^14,24,65^ the V1 population in frogs spanned nearly the entire ventral horn of the spinal cord and in both species, with a peak distribution at the middle of the mediolateral axis of the hemi-section (**Figure 4G-H; Figure S5B-C**), again demonstrating potent cross-species conservation.

### V1 subtype heterogeneity emerges during frog metamorphosis

In mammals, V1 interneurons are subdivided by transcription factor expression into four distinct clades and various subtypes that vary in their molecular profile, settling position, projection patterns and electrophysiological properties.^11,14,16,17,66–69^ They include the physiologically defined Renshaw cells and Ia-inhibitory neuron subpopulations for recurrent and reciprocal inhibition, respectively.^17,60^ Whether this high level of V1 heterogeneity represents a specialization for complex limb movement in mammals or a conserved trait across vertebrates remains unknown.

We profiled expression of nine transcription factors at larval swim, free swim and developing limb stages, focusing on the clade markers—MafA, Pou6f2, FoxP2, and Sp8 (**Figure 5A-D**)— and other transcription factors (TF) that subdivide V1 interneurons in mice (**Figure S6A-D**; Bikoff et al., 2016). At larval stages, V1s were largely homogeneous and lacked three out of four clade markers (NF35-38; **Figure 5A, D**). Approximately half of larval V1 interneurons expressed MafA and MafB (**Figure 5A, D; S6A,D**), a Renshaw cell-like expression profile.^70^ With the emergence of free-swimming and the development of hypaxial musculature, organs, and dorsal root ganglia,^6^ we observed a sharp rise in V1 transcriptional heterogeneity (NF44-47; **Figure 5B, D; Figure S6D-H**): the Pou6f2 and FoxP2 clades emerged (**Figure 5D**) and the proportion of FoxP1-, Otp-, and Nr3b3-positive V1 interneurons increased (**Figure S6D-H**). Then, with limb circuit addition at NF54-55, we observed another dramatic increase in V1 transcriptional heterogeneity with the expression of all TFs including the four V1 clade markers (**Figure 5C-D; Figure S6C-D, G-H**). Expression of the FoxP TFs exemplified this increase: at larval stages, only FoxP4 was expressed, subdividing V1 interneurons into on/off populations; at free-swimming stages, FoxP1, P2, and P4 were all expressed but only in a small percentage of V1 interneurons; after metamorphosis, all FoxP-expressing V1 subsets further increased at both thoracic and limb levels (**Figure S6G-H**). In addition, as in mice,^17^ the first clade to emerge was MafA-positive, followed by the Pou6f2-positive and the FoxP2-positive, and finally the Sp8-positive clades (**Figure 5E**), indicating a remarkable conservation between species in the timing of V1 interneuron development.

**Figure 5.**
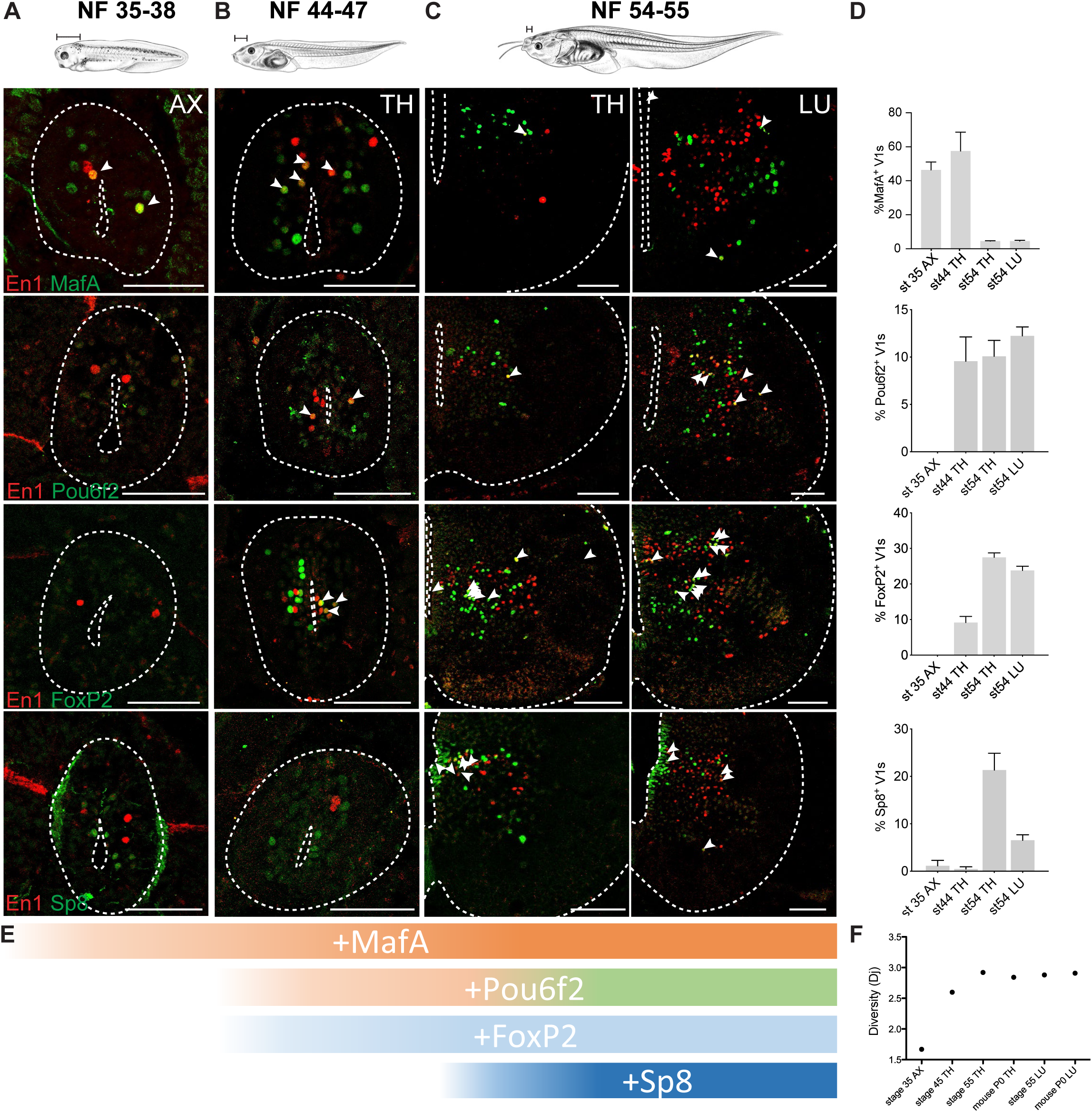
Temporal pattern of V1 clades emergence during the *Xenopus* swim-to-walk transition. **A** At escape swimming stages, NF35-38, V1s show little diversity. The Pou6f2, FoxP2, and Sp8 clades are absent, and around 50-60% of V1s are marked by MafA and MafB. **B.** At free-swimming stages, NF44-47, V1s start to diversify. The Pou6f2 and FoxP2 clades emerge. **C.** During metamorphosis, with limb emergence, the Sp8 clade emerges and V1s acquire the four clade organization observed in the mouse. **D.** Percentage of V1 interneurons expressing a single clade marker (FoxP2, Pou6f2, Sp8, MafA) in axial (NF35-38), thoracic (NF44-47 and NF54-55) or lumbar spinal cord (NF54) (mean ± SEM, n = 4–10 animals). **E.** The sequence of V1 clade emergence. MafA present in escape swimming. MafA, Pou6f2, FoxP2 present in free-swimming. All four clades present at limb-circuit stages. **F.** Entropy analysis of “diversity” index based on transcription factor expression shows a significant increase in overall transcriptional diversity between NF35 and NF45, and a peak of diversity reached at NF54-55. The diversity at the peak matches that of the neonate mouse. Tadpole drawings adapted from Xenopus illustrations © Natalya Zahn (2022).^117^

To quantify the level of transcriptional diversity across stages, we utilized entropy analysis, which is a measure of diversity by information theory. Based on fractional TF expression at each stage, the transcriptional entropy was calculated.^71^ We found an initial increase in entropy between larval and free-swimming stages, and a second jump at metamorphosis with the development of limb circuits (**Figure 5F**). At peak limb circuit development, frogs and mice were comparable in their level of entropy (**Figure 5F**).

Our analysis thus demonstrates that as frogs transition from larval escape to free-swimming and eventually to limb movement, they not only add more V1 interneurons but those V1 interneurons diversify into subpopulations that share the same transcription factor expression profiles as mice.

### Conservation of limb- and thoracic-level V1 transcriptional diversity between frog and mouse

Next, we directly compared V1 transcriptional subtypes in frogs and mice, focusing our analysis on the NF54-55 frog and P0 mouse stages, which represent the respective peaks of V1 interneuron differentiation in both species (Bikoff et al., 2016; **Figure S4C-D, H-K)**. In mice, several rules define V1 transcriptional heterogeneity.^14–16^ (*i*) V1 clades are non-overlapping, differ in their relative size, and occupy distinct settling positions; (*ii*) a panel of additional TFs mark different proportions of V1 interneurons with distinct mediolateral-dorsoventral positions; (*iii*) functionally defined populations such as Renshaw cells and Ia-inhibitory neurons are distinct in their transcriptional profiles^17,60,72^; and (*iv*) additional V1 subtypes are generated by the combinatorial expression of two and three TFs, with some co-expressed and others never overlapping.

Remarkably, in the lumbar spinal cord, we found nearly all hallmark features of mouse V1 interneurons were recapitulated in the frog. First, the four V1 clade markers—MafA, Pou6f2, FoxP2, Sp8—were mutually exclusive in their expression (**Figure 6A-C**) and marked the same proportion of V1s across species, with FoxP2 the largest, Sp8 and Pou6f2 intermediate, and MafA the smallest in size (**Figure 6D)**. Their relative settling positions were also similar: V1^Sp^^8^ interneurons were medial, V1^Pou6f2^ lateral, V1^FoxP2^ central, and V1^MafA^ the most ventral (**Figure 6B**). Second, 10 of 19 TFs associated with V1 diversity in the mouse labeled a similar percentage of V1 interneurons in the frog (**Figure 6D**). These one TF (V1^1TF^) populations each settled in distinct dorsoventral and mediolateral positions (**Figure S7K-Q**). Third, we detected a subpopulation of ∼10% of V1s that co-expressed MafB and Calbindin (**Figure S7G-G’’**), corresponding to the molecular signatures of Renshaw cells and suggesting their conservation between frogs and mice. Fourth, two (V1^2TF^) and three (V1^3TF^) TF combinations in frogs carve out numerous molecularly distinct subpopulations as in mice.^14,15^ Analysis of 24 V1^2TF^ combinations found that the rules of mouse TF co-expression in V1s were largely upheld in the frog (**Figure 6E**, top). FoxP1 and FoxP4 transcription factors never were co-expressed with Pou6f2 or Sp8, and Otp was only co-expressed with FoxP2 and Sp8 but not Pou6f2. At the level of V1^2TF^, species differences emerged in V1 subset proportion: out of the 24 V1^2TF^ combinations tested, four labeled mouse-enriched V1 subsets and two labeled frog-enriched V1 subsets. Evaluation of V1^3TF^ revealed even more populations that were species-enriched (**Figure 6E**, bottom). Interspecies differences can thus be revealed by the expression, or lack thereof, of specific combinations of TFs.

**Figure 6.**
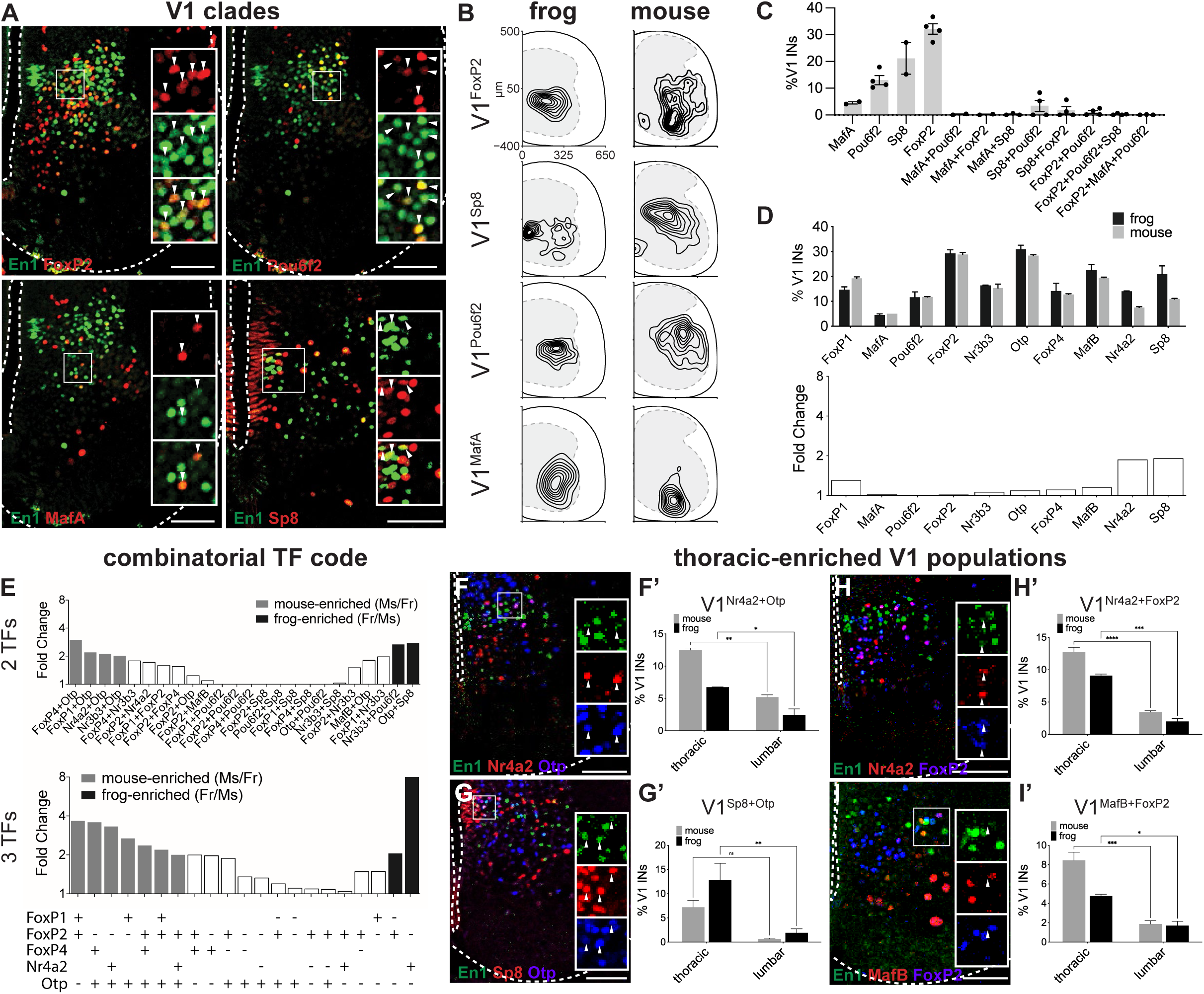
Conservation of V1 clade organization and transcriptional diversity between frog and mouse. **A. Mouse V1 clades are conserved in the frog**. Antibodies against FoxP2, Pou6f2, MafA, and Sp8 transcription factors (green) label subsets of En1+ V1 interneurons (red) in lumbar spinal cord of NF54-55 tadpoles. Shown is a ventral hemi-section of spinal cord with the central canal and outer edge indicated (dotted line). **B. Spatial distribution plots of V1^FoxP2^, V1^Sp8^, V1^Pou6f2^, and V1^MafA^ at the lumbar level in NF54-55 tadpole (left) and P0 mouse (right).** Frog and mouse V1 clades have similar settling positions. Frog spinal cords were resized to mouse-like proportions (see **STAR Methods**). Plotted are interneurons from at least 20 lumbar spinal cord hemi-sections from at least 2 animals. **C. Frog V1 clades are mutually exclusive in their expression**. The bar plot shows the percentage of V1s expressing a singular or combination of clade markers, FoxP2, Sp8, Pou6f2 and MafA. Shown is mean ± SEM, n = 2–4 animals **D. V1 molecular subsets are present in similar proportions in the frog and mouse**. Upper bar plot shows the percentages of V1s expressing a given transcription factor in lumbar NF54-55 tadpole (black) and P0 mouse (gray) spinal cords determined by IHC. Lower bar plot shows the fold change in percentage of V1 subsets between the frog and mouse; no change larger than 2-fold observed. Shown is mean ± SEM (2TF: n = 2–6 animals; 3TF: n = 2 animals). **E. V1 interneurons marked by two and three transcription factors reveal species-enriched subsets.** Shown is fold enrichment of V1^2TF^ and V1^3TF^ interneurons with > 2-fold enrichment in NF54-55 frog (black) or P0 mouse (gray) spinal cord. **F-I**. **Mouse thoracic-enriched populations of V1 are present and enriched at the thoracic levels in frog.** V1^Nr4a2+Otp^, V1^Sp8+Otp^, V1^Nr4a2+FoxP2^, V1^MafB+FoxP2^ populations are present in the frog (gray) thoracic spinal cord (left) and significantly enriched compared to the lumbar level (right). The same rosto-caudal enrichment was reported in the mouse (black). Shown is mean ± SEM for n = 2–6 animals with significant differences (p < 0.05) plotted.

Following our limb-level V1 analysis, we evaluated V1 interneurons at thoracic levels, the location of autonomic- and torso -associated motor circuits.^12^ Our analysis showed a downscaling of thoracic circuits with half as many MNs and V1 interneurons at thoracic compared to both brachial and lumbar levels (**Figure S7A-D),** as in mice.^16,47,64,67^ As at limb levels, the proportions of 11 out of 27 V1^1TF^ subpopulations at thoracic levels were the same between species (**Figure S7E-F**). The settling positions of the thoracic V1 clades and most V1^1TF^ subpopulations were distinct within the ventral horn (**Figure S7H-Q**). V1^2TF^ combinations carved out species-specific thoracic V1 subpopulations (**Figure S7F**) and in both frogs and mice, thoracic-enriched V1^2TF^ subpopulations were marked by Nr4a2+Otp (**Figure 6F**), Otp+Sp8 (**Figure 6G**), FoxP2+Nr4a2 (**Figure 6H**), and FoxP2+MafB (**Figure 6I**). Our analysis of V1 diversity within thoracic spinal segments thus demonstrated a high degree of subset conservation between frogs and mice.

### FoxP1 loss-of-function impairs limb movement

Our cell-type analysis of frog spinal circuits revealed a high degree of conserved molecular and anatomical features between frog and mouse spinal neurons. Though molecularly aligned, it was still unclear whether these transcriptionally similar neural subsets perform similar functions within these distantly related locomotor circuits. To test their functional conservation, we thus used CRISPR gene editing to perturb MNs and V1 interneurons in frogs, evaluated the effect of this loss-of-function on cell types and behavior, and compared the resulting phenotypes with those from similar perturbations in mice.

In mice, the transcription factor FoxP1 determines the lateral motor column (LMC) of limb MNs.^47,73^ In the absence of FoxP1, LMC MNs lose their cell identity and mutant mice lack coordination of limb movement.^74^ Given its conserved expression, we determined whether this transcription factor similarly specified LMC identity in frogs. To perturb FoxP1 gene expression, we took advantage of recent advances in CRISPR/Cas9-mediated loss-of-function in F0 *Xenopus* frogs.^75^ By injecting a single guide RNA (sgRNA) together with Cas9 protein into one cell at the two-cell stage (**Figure 7A**), or into one cell-stage (**Figure S8A**), embryos, we generated unilateral or bilateral FoxP1 CRISPR mutant animals respectively, as evaluated by immunohistochemistry (**Figure 7B; Figure S8B**) and TIDE analysis (**Figure S8E-F**). Both unilateral and bilateral FoxP1 CRISPR animals developed normally up to limb-based stages, after which bilateral mutant animals exhibited defective limb posture and survived only until NF64, similar to mutant mice which die at birth.^47^ Unilateral mutants, however, were largely unaffected in their overall morphology, with only the fore and hindlimbs on the mutant side exhibiting an abnormal posture (**Figure 7F**). Unilateral FoxP1 CRISPR animals were thus used to examine the function of FoxP1 in limb-driven locomotion, while bilateral mutant tadpoles were used to study whether FoxP1 affects tail-based swimming.

**Figure 7.**
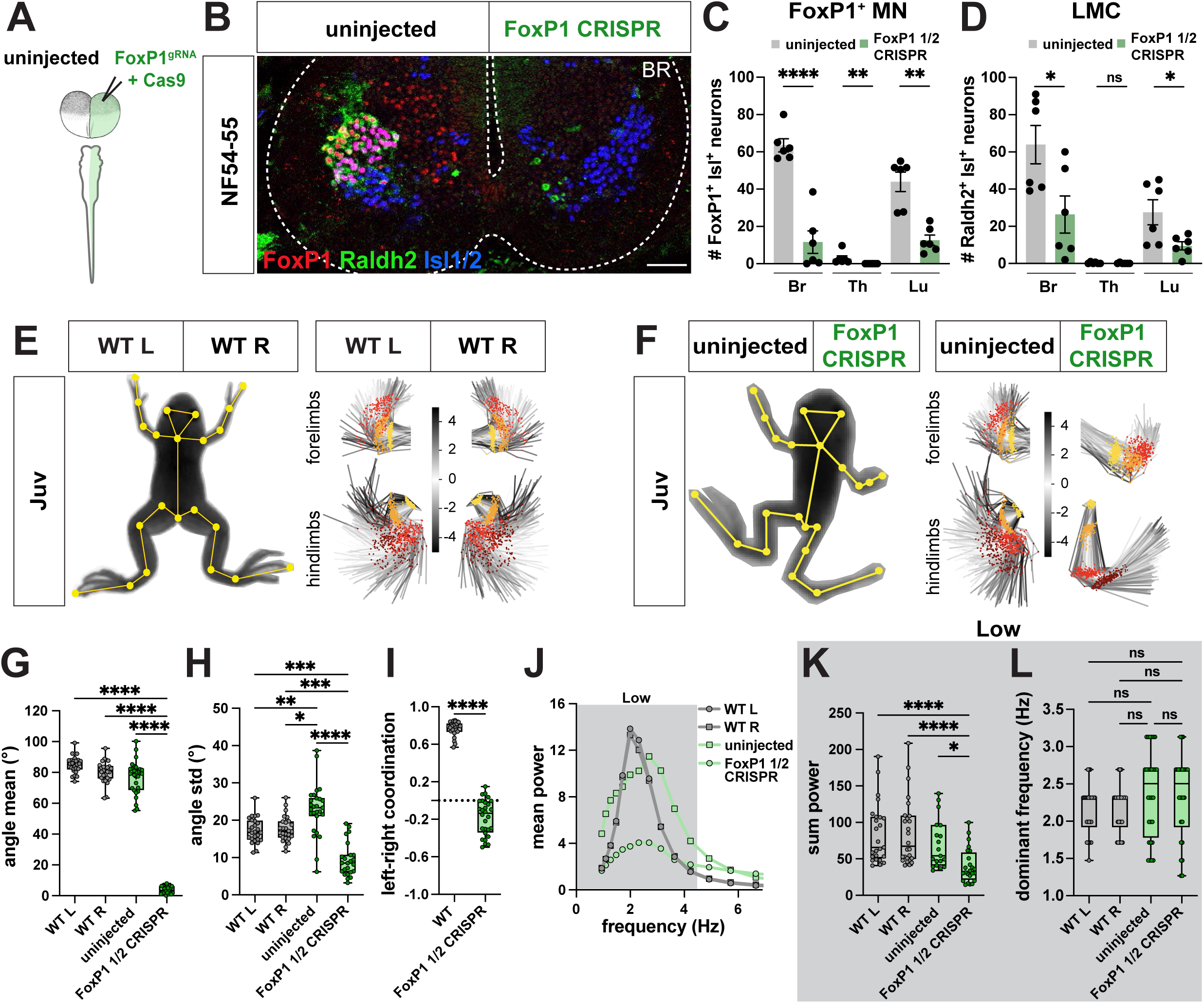
FoxP1 CRISPR loss-of-function causes loss of range and coordination of limb movement in *Xenopus* frogs. **A-B.** Generation of unilateral FoxP1 CRISPR mutant frogs by injection of FoxP1 sgRNA and Cas9 protein in a single cell at two-cell stage (**A**) results in NF54-55 tadpoles in which FoxP1 (red) and Raldh2 (green) immunoreactivity is selectively absent from the mutant side of the spinal cord (**B**). Isl1/2-positive (blue, marker for motor neurons in ventral spinal cord) neurons are present on both wildtype and mutant side of spinal cord (**B**). Scale bar, 50 μm. **C-D. Cell-type characterization in unilateral FoxP1 CRISPR mutant animals.** Quantification of spinal cord cell numbers at brachial (Br), thoracic (Th) and lumbar (Lu) reveals loss of FoxP1+ Isl1+ neurons at all levels (**C**; uninjected vs. FoxP1 ½: Br, p = <0.0001; for Th and Lu, p = 0.002) and loss of Raldh2+ Isl1+ neurons at brachial and lumbar levels (**D**; uninjected vs. FoxP1 ½: Br, p = 0.025; Lu, p = 0.029). n = 6 for WT, n = 6 unilateral FoxP1 CRISPR. **E-I. Loss of range and coordination of movement of the FoxP1 mutant hindlimb.** WT (**E**) and unilateral FoxP1 CRISPR (**F**) juvenile frogs with SLEAP skeleton (left, yellow) superimposed on animal image. PCA plots represent the position of the fore and hind limb and their range of movement during 256 random frames and show a different position and range of the FoxP1 CRISPR limbs compared to WT or the uninjected side (**E-F**, right; hip and shoulder, yellow; knee and elbow, orange; ankle and wrist, red; foot, brown). Scale bar in **E** and **F** indicates the color-code of the first principal component of variation of the aligned fore and hind limb positions. The FoxP1 CRISPR mutant knee also differs in its mean angle (**G**; for WT L versus FoxP1 ½, WT R versus FoxP1 ½ and uninjected versus FoxP1 ½, p = <0.0001), and its movement range is reduced (**I**; WT L versus FoxP1 ½, p = 0.0006; WT R versus FoxP1 half, p = 0.0002; uninjected versus FoxP1 1/2, p = <0.0001). In contrast, the uninjected side displays a higher range of movement (**H**; WT L versus uninjected, p = 0.009; WT R versus uninjected, p = 0.025). Left-right coordination between knee joints is lost in FoxP1 CRISPR animals (**I**; +1 = bilateral synchronous, 0 = random, −1 = alternate synchronous; WT versus FoxP1 ½ CRISPR, p = <0.0001). n = 13 for WT, n = 14 for unilateral FoxP1 CRISPR. **J-L. FoxP1 CRISPR mutant hindlimbs maintain dominant frequency but lose power**. Mean power spectrum of knee oscillations shows only one peak in the low frequency range for WT, uninjected and FoxP1 CRISPR hindlimbs (**J**; 0.9-4.5 Hz, dark gray). At the knee joint, the amount of movement in the low frequency bin (0.9-4.5 Hz), represented by the sum power, is lower on the mutant side compared to both the uninjected side and WT (**K**; for WT L versus FoxP1 ½ CRISPR and WT R versus FoxP1 ½ CRISPR, p = <0.0001; uninjected versus FoxP1 ½ CRISPR, p = 0.021). Dominant frequency is unaffected on both uninjected and FoxP1 CRISPR sides (**L**). n = 13 for WT, n = 14 for unilateral FoxP1 CRISPR.

We first evaluated FoxP1 loss-of-function at a cellular level. Using the pan-MN marker Isl1/2 alongside a FoxP1 antibody, we determined that the loss of FoxP1 was highly efficient, with an ∼80% loss of FoxP1 immunoreactivity in MNs on the mutant side (**Figure 7B-C**). An independent marker of LMC identity, the retinoic acid dehydrogenase gene Raldh2/Ald1a2, was similarly ∼70% reduced on the mutant side at limb levels (**Figure 7B, D**), consistent with previous findings in mice^47^ and supporting a conserved role for FoxP1 in establishing LMC cellular identity. In contrast, the number of MMC MNs (**Figure S8G)**, V1 inhibitory neurons (**Figure S8H)** and twelve V1 subtypes (**Figure S8I-J)**, with the exception of V1^FoxP^^1^, V1^FoxP2^ and V1^FoxP2+Otp^, were not reduced in number. Further analysis of MN axon projections in FoxP1 CRISPR mutants revealed that innervation of the limb was maintained, despite disruption of its molecular identity (**Figure S8K-L**), also consistent with observations in mice.^47^ Together, these results support a remarkable conservation in FoxP1-dependent specification of limb MNs across tetrapods.

To test the contribution of FoxP1 to limb-driven locomotion, we evaluated the motor behavior of unilateral FoxP1 CRISPR animals at juvenile stages. On a coarse level, we observed a loss of fore- and hindlimb movement on the mutant side (**Movie S8**). FoxP1 mutant frogs moved less, traveled shorter distances, and had reduced acceleration (**Figure S9A-E)**. However, this unilateral loss of movement did not influence the amount or direction of turning (**Figure S9F**). Further evaluation of the effects of FoxP1 mutation on limb kinematics via principal component analysis revealed differential positions of mutant hind and forelimb compared to WT (**Figure 7E-F)** and showed a clear loss in the range of movement on the mutant side, as measured by the mean and standard deviation of its angle (**Figure 7G-H; Figure S9G)**. The foot joint, however, was less affected (**Figure S9G**), as in the mouse.^74^ In contrast, on the uninjected side, an increased range of movement was observed in unilateral FoxP1 mutants (**Figure 7H**), potentially compensating for the impaired limb movement on the other side. This one-sided phenotype was also accompanied by a decrease in the synchrony of left-right hindlimb coordination (**Figure 7I; Figure S9H**). The residual movement of the mutant hindlimb was at the same dominant frequency as the uninjected side; however, its power was reduced (**Figure 7J-L; Figure S9I**). Finally, this selective loss of limb movement allowed evaluation of the interdependence of tail and limb circuit function. At a behavioral level, during free-swimming, overall movement (**Figure S9J-O**) and tail range were unaffected during free-swimming (**Figure S9J, P**), despite the loss of the pioneering LMC neurons. Taken together, our results demonstrate that FoxP1 is a conserved determinant of limb motor neuron identity and movement in frogs like mice.

### En1 loss-of-function impairs frequency of limb movement

We next extended our CRISPR mutant analysis to target V1 interneurons. In zebrafish, ablation of V1 interneurons lowers swimming frequency.^58^ Similarly, V1 ablation in mice reduces fictive locomotor frequency^76^ and in vivo, causes severe limb hyperflexion.^61,77^

We targeted V1 interneurons in *Xenopus* by designing a sgRNA to knock out the transcription factor En1 and used it to generate bilateral and unilateral mutant animals. To test the efficiency of En1 CRISPR loss-of-function, we evaluated the mutation rate and number of En1-positive cells in wild type versus CRISPR mutant animals (**Figure 8A-C**). Based on TIDE analysis,^78^ bilateral mutant animals were on average ∼80% mutant, whereas unilateral mutants were ∼25% mutant (**Figure 8C**). This high efficiency of gene disruption was recapitulated at the level of En1 protein expression, with a near complete loss of En1 immunoreactivity on both sides in bilateral (**Figure 8A**) and on one side in unilateral CRISPR mutants (**Figure 8B**).

**Figure 8.**
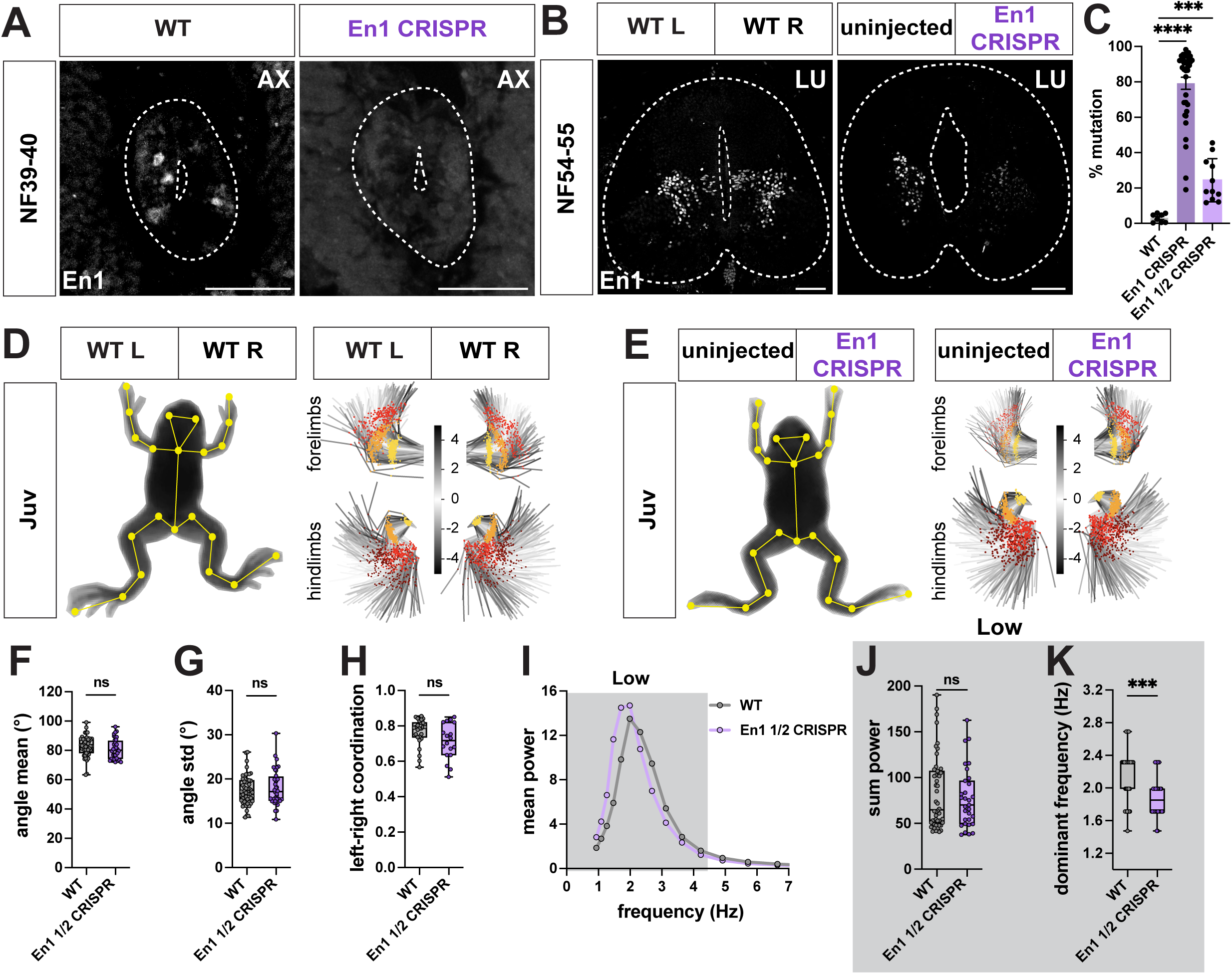
En1 CRISPR loss-of-function causes a loss of limb frequency in juvenile *Xenopus laevis*. **A-C.** Characterization of En1 CRISPR mutant animals En1 sgRNA and Cas9 protein were injected at one cell stage to generate bilateral mutant animals, resulting in loss of En1 immunoreactivity from both sides of the spinal cord at NF39-40 (**A** right, white), or in one cell at two-cell stage to generate unilateral mutant animals with loss of En1 immunoreactivity only from the En1 CRISPR side of the spinal cord at NF54-55 (**B** right, white). TIDE analysis reveals high efficiency of En1 sgRNA in generating NF44-48 bilateral, and ∼25% mutation rate for juvenile unilateral CRISPR animals (**C**; n = 8 for WT, n = 36 for En1 bilateral mutants, n = 8 for En1 unilateral mutants). Scale bar, 50 μm. **D-H. Range and coordination of movement are unaffected in juvenile unilateral En1 CRISPR mutant animals.** WT (**D**) and unilateral En1 CRISPR (**E**) juvenile frogs with SLEAP skeleton (yellow) superimposed on animal image. PCA plots represent the position of the fore and hind limb and their range of movement during 256 random frames and show no visible difference in range between WT, uninjected and En1 CRISPR sides (**D-E,** right; hip and shoulder, yellow; knee and elbow, orange; ankle and wrist, red; foot, brown). Scale bar in **D** and **E** indicates the color-code of the first principal component of variation of the aligned fore and hind limb positions. Unilateral En1 CRISPR mutant knees show similar mean angle (**F**) and angle range (**G**) as WT. Left-right coordination between knee joints is also unaffected in unilateral En1 CRISPR animals, as their pattern of movement resembles the bilateral synchronicity of WT (**H**; +1 = bilateral synchronous, 0 = random, −1 = alternate synchronous). N = 13 for WT, n = 8 for unilateral En1 CRISPR. **I-K. Lower dominant frequency in juvenile En1 CRISPR mutant hindlimbs.** Mean power spectrum of the knee oscillation shows only one peak in the low frequency bin (0.9-4.5 Hz, dark gray) for WT and unilateral En1 CRISPR animals (**I**). At the knee joints, the amount of movement, represented by the sum of the power, is similar between WT and unilateral En1 CRISPR animals (**J**). However, the dominant frequency of the knees is lower in unilateral En1 CRISPR animals compared to WT (**K**; WT vs En1 ½ CRISPR, p = 0.0002). n = 13 for WT, n = 8 for unilateral En1 CRISPR.

As V1 interneurons play a critical role in generating tail-based locomotion in zebrafish^58^, we evaluated the effect of bilateral En1 loss-of-function in the free-swimming tadpole stage using our pose-estimation pipeline (**Figure S10A-L**). Analysis of the overall movement of En1 mutant tadpoles revealed a reduction in the time spent moving, distance traveled, speed, and acceleration, compared to WT tadpoles (**Figure S10A-E**). En1 mutant tadpoles turned twice as much as WT animals and exhibited a larger range of tail movement (**Figure S10A,F-G**), an observation not previously made in larval zebrafish. However, bilateral En1 loss-of-function in tadpoles, similar to V1 disruption in zebrafish,^58^ caused a loss of high frequency and a gain of low frequency tail oscillation, with the dominant low frequency also lowered in mutant animals (**Figure S10H-L**). Our experiments prove a conserved role for En1 in regulating the frequency of swimming in tadpoles and zebrafish.

We next examined whether limb movement was affected in En1 CRISPR mutant frogs, as in mice.^79^ As bilateral En1 CRISPR mutants die by NF48 due to edema,^80^ a phenotype we also observed in our mutants, we only analyzed unilateral mutant juvenile frogs, which showed no edema, physical deformations, or any major deficits in overall movement (**Figure 8D-E, Figure S10M-R; Movie S9**). As no limb movement defects were visible by eye, and the precision of TIDE analysis did not allow us to prospectively determine which side was mutant, we pooled the data from the uninjected and injected sides of the unilateral En1 CRISPR mutants. At all hindlimb joints, quantification of limb movement in unilateral En1 mutant animals showed a selective decrease in the dominant frequency but not in its power or range of movement (**Figure 8F-G, I-K; Figure S10S-T,V-W)**. The coordination between left and right hindlimbs was also unaffected, with the mutant animals still capable of performing synchronized bilateral movements at this lower frequency (**Figure 8H; Figure S10U**).

Loss-of-function of the transcription factor En1 in tadpoles thus recapitulated the frequency defects observed after V1 disruption in zebrafish, and in frogs, disrupted the frequency of limb movement without decreasing its range or left-right coordination.

## DISCUSSION

Understanding how neural circuits are organized to implement movements of varying levels of complexity is a fundamental goal of neuroscience. Our cellular and behavioral analysis of spinal circuits during frog metamorphosis demonstrates that the number and diversity of spinal neurons increases dramatically as tadpoles transition from tail-based undulatory swimming to multi-limb behavior. This developmental transition culminates in limb circuits that exhibit a high degree of similarity to mammalian motor circuits, as shown by their conserved molecular cell type heterogeneity and function. Taken together, our study thus provides a template by which motor circuits scale in a manner that accommodates behavioral repertoires of increasing sophistication across development and evolution.

### Implications for increasing spinal neuron number and transcriptional heterogeneity

One of the most striking observations from our study is the dramatic expansion in spinal neuron number and diversity during frog metamorphosis. This increasing neuronal heterogeneity is largely governed by transcription factor expression during spinal circuit maturation. In the case of MNs, there is a well established precedent that differential transcription factor expression controls anatomical and functional distinctions in MN identity and circuitry in multiple species.^12^ Consistent with this hypothesis, limited neuronal diversity at early tadpole stages aligns with the anatomical or physiological homogeneity of motor or V1 interneurons at these early stages.^7^ A similar principle has been observed for other classes of spinal interneurons in the mouse,^3^ in which a temporal transcription factor code has been shown to subdivide brain and spinal neurons into long and short-range projecting interneuron populations.^68,81^ In the case of V1 interneurons, differences in clade TF expression in the mouse correlate with distinct motor and sensory neuron connectivity, intrinsic physiology, and target innervation patterns.^14,17^ These anatomical and functional properties are also mirrored by significant differences in ion channel expression across V1 clades. Analogous changes in anatomy, channel, and receptor expression are likely to characterize the emergence of limb motor circuits during frog metamorphosis, as suggested by the upregulation of dopamine D1 receptor,^82,83^ NOS^84,85^ and serotonin^86^ signaling.

### Developmental remodeling of spinal circuits during frog metamorphosis

Our finding that neuron number progressively increases from larval to free-swimming and then to limb stages demonstrates that developmental transitions in spinal circuits do not rely solely on a simple rewiring of existing neurons, but also on the generation and incorporation of new neurons. This raises several questions about neural circuit remodeling: (*i*) what drives changes in number and diversity, (*ii*) how are new neurons integrated into an existing circuit, and how do the new and existing circuits interact, and (*iii*) how are these circuit-level interactions reflected in locomotor behavior. The late-stage proliferation of progenitors during metamorphosis,^87,88,88^ long after embryogenesis and primary neurulation, requires a second phase of patterning to differentiate these newly added neuron types.^89^ We have found that the ectopic application of thyroid hormone induces the premature generation of motor and interneuron diversity (data not shown), consistent with the critical role of this signaling pathway in metamorphosis.^48,90,91^ In addition, as demonstrated in larval tadpoles, neuronal activity may contribute to specifying the subtype and neurotransmitter identity of such late-born neurons, either via homeostatic mechanisms^92,93^ or by interaction with other cues specifying cell identity.^94^ Newly generated limb-circuit motor and interneuron types must also be integrated into the existing swim circuit. Our discovery of pioneering LMC neurons suggests a role for larval circuits in facilitating the incorporation of new neurons into the spinal motor architecture, as observed during chick hindlimb development^95,95^ and during mouse MN development.^96,97^

It is less clear whether neurons in the larval swim circuit persist until after metamorphosis, and if so, whether they become functionally integrated into later-born limb circuitry alongside newly generated neurons. Initial studies indicate that primary axial MNs, identified by their large size, remain during metamorphosis.^88,98^ Persisting circuits could instruct newly emerging ones, supported by the rhythmic coupling of the swim and limb circuits that we observe at transitional metamorphic stages and that was previously described using *ex vivo* electrophysiological recordings.^34^ Such subjugation of new-to-existing circuits, or of receding-to-remaining circuits, newly described here, provides a generalizable mechanism of neural circuit remodeling via circuit-level interactions.

### Conservation and divergence of motor and V1 interneuron cell types

As the first vertebrates to emerge with limbs, amphibians occupy an optimal position on the vertebrate phylogenetic tree.^99^ Many also switch from a larval stage adapted for aquatic life to a limb-stage adapted for terrestrial locomotion during metamorphosis. This transition effectively recapitulates nearly 360 million years of evolutionary changes in behavior in just a few months of development. From this evolutionary-developmental perspective,^100^ our study offers a unique opportunity to examine the mechanisms underlying the vertebrate transition from water to land, the locomotor strategies employed during this switch, and the origin and coordination of the neural types underlying aquatic versus terrestrial locomotion.

The locomotor switch from swimming to limb propulsion requires the differential recruitment of axial versus appendicular muscles. In *Xenopus*, we find that larval MNs have a uniform, axial MMC-like molecular phenotype, consistent with what has been observed anatomically in lamprey^101^ and in larval zebrafish,^102^ though zebrafish exhibit additional molecular, anatomical and physiological distinction between MMC MNs that are involved in slow, intermediate, and fast locomotion.^103,104^ Moreover, along the rostrocaudal axis, *Xenopus* MNs segregate into a LMC, divisions and pools, just as they do in mice, with FoxP1 serving as an evolutionarily conserved specifier of LMC MN fate. Thus, shared strategies are employed to establish the building blocks of motor circuits.^105^ Recent findings in zebrafish and skate support that this common logic of MN specification may even extend beyond amphibians.^104,105^

While MNs have previously been shown to exhibit common molecular features across different vertebrate species, the extent to which spinal interneuron diversity is conserved is less well understood. The metamorphic switch from tail to limb movement necessitates not only recruitment of new axial and limb muscles but also regulation of their frequency and coordination across joints and limbs. In mice, such regulation is conferred in part by V1 inhibitory neurons, which influence the speed and range of limb movement.^24,56,61^ Here, we demonstrate that the transcriptional logic that subdivides V1 interneurons in the mouse similarly applies to the frog, with all four V1 clades present, including the V1^Foxp2^ clade, which is thought to contain group Ia interneurons involved in flexor-extensor alternation.^17,24,60^ This expansion of V1 interneuron diversity into molecularly distinct and spatially enriched subpopulations in parallel with the emergence of complex, limbed behavior in *Xenopus,* indicates that a high degree of spinal interneuron heterogeneity is not an exclusive feature of murine, or mammalian, motor circuits. Intriguingly, our finding that the transcription factor En1 regulates the frequency but not the range of movement in frogs coincides with similar observations in mice,^79^ demonstrating potent conservation of its function across species.

In larval tadpoles, in contrast, axial swimming can be generated without the high diversity of V1 interneurons observed later in development, a finding that is further corroborated by the homogeneity of larval V1 electrophysiological properties.^55^ Instead, at this stage, one dominant V1 clade is sufficient to control rostrocaudal and left-right coordination of axial muscles necessary for undulation. By extension, this predicts that V1 interneurons in lamprey and other organisms with axial-based escape swimming responses, may similarly lack V1 clade diversity. Molecular and functional heterogeneity, however, could exist for other interneuron subtypes involved in swimming, as has been described for excitatory interneurons in larval zebrafish.^106^ As such, a largely molecularly and physiologically homogeneous set of V1 interneurons may have originated in an aquatic ancestor of vertebrates, with additional clades comprising terrestrial V1 subtypes acquired later during evolution as a limb locomotor adaptation during the water-to-land transition. Our study in *Xenopus* thereby paves the way for future transcriptomic studies of V1 circuits in other understudied vertebrates, such as Agnathans, cartilaginous, and bony fish, that will shed more light on spinal interneuron adaptations during the evolutionary transition from water to land.

### Linking changes in cell types to motor strategies across vertebrates

Across frog metamorphosis, the developmental expansion of neuronal diversity reflects changes not only in anatomy but also in behavioral strategy. Larval tadpoles perform spiraling escape swimming; free-swimming tadpoles constantly swim in a directed manner so that they can freely feed; and frogs kick, hop, scoop and walk to support feeding, mating, and survival both in water and on land.^36^ Comparing tail-based swim behavior across species, zebrafish have a tail oscillation frequency of 15-100 Hz with a ∼42-degree average displacement,^107^ whereas, as we report here, *Xenopus* tadpoles exhibit a ∼1.6 Hz and ∼10 Hz peak frequency and only a maximal ∼10-degree displacement. In zebrafish, this high frequency movement has co-evolved with, and is supported by, the subdivision of many classes of motor and spinal interneurons into slow, intermediate and fast subpopulations.^8^ *Xenopus* tadpoles do not appear to exhibit such sub-specialization,^7^ likely reflecting their more limited behavioral repertoire.

With respect to limb-based movement in frogs and mice, while some common features exist in locomotor kinematics, such as a shared frequency of limb movement at ∼2.5 Hz,^37^ there are also substantial differences. Frog locomotion is largely dominated by left-right bilateral synchronous hindlimb kicking, and is not characterized by walking, trotting and galloping gaits with increasing speed.^36^ *Xenopus* specifically, a member of the “belly flopping” frogs,^108^ are largely not weight-bearing, unlike mice that move almost entirely with their torso off the ground.^109^ Moreover, in contrast to mice, frogs are not capable of performing fine-motor skills such as grasping, but instead execute coarse movements such as wiping.^110^

We now show that common behavioral features can be linked to highly conserved MN and V1 cell types. These relationships are specific to individual gene programs, as En1 loss-of-function disrupts both swim and limb movement whereas FoxP1 loss selectively affects limb movement, and they extend across species. The defects in limb movement in frogs perfectly recapitulate those in mice after similar FoxP1 or En1 mutations^47,79^; the disruption of swimming in tadpoles after En1 loss-of-function matches the effect of V1 ablation in zebrafish.^58^ Despite this conservation, the open question is what underlies the divergent features between species. One attractive source of divergence that emerges from our study is the fine-scale differences in the number, position and molecular identity of interneuron subtypes between mice and frogs. In mice, such differences relate to variant descending, sensory, and motor inputs.^14,17^ In frogs, these properties clearly differ: they reportedly lack a corticospinal tract^111^ and gamma MNs,^112,113^ crucial regulators of motor precision and muscle spindles for weight bearing, respectively. Future analyses in frogs, which are amenable to transcriptomic profiling, anatomical tracing,^114^ CRISPR-based loss-of-function studies, and behavioral tracking, will help parse the mechanisms of species-specific behaviors.

Taken together, our data reveal how vertebrate movement is generated using highly conserved cellular building blocks that comprise a shared substrate for limbed-based locomotion and scale with the swim-to-limb transformation during frog metamorphosis, revealing a fundamental principle of cell type scaling for complex behavior.

## LIMITATIONS OF OUR STUDY

Aligning developmental stages between two species from different vertebrate classes, especially those developing in distinct environments (*in utero* for mice versus *ex utero* for frogs) presents a challenge. For our cross-species comparison of cell types, we focused on the stages characterized by the peak of motor and interneuron generation: *Xenopus* NF54-55 (see the time course in **Figure S4**) and mouse embryonic stages E13.5–P0.^14,47,64^ These stages are also comparable in terms of limb development and muscle innervation.^115^

Due to the long generation time of *Xenopus* (>1 year to sexual maturity), our loss-of-function analysis was conducted using only the first generation of CRISPR-edited animals (F0). Though we took steps to ensure consistent mutant conditions by performing genotyping and immunohistochemistry, as outlined in **STAR Methods**, there is potential variability in mutation efficiency among the F0 animals.

On a molecular level, our study examines the metamorphic transformation of spinal circuits through the analyses of V1 interneurons and MNs because of their district roles in controlling swim and limb-based movement. Concurrently, other spinal and supra-spinal neuronal types emerge and begin to shape motor output in the frog, with their role in movement not examined here.

Finally, our study establishes a link between frog behavior and cellular changes in the spinal circuitry. While we characterized a diverse array of motor behavior patterns during frog development (e.g., escape swimming, free swimming, turning, four-limb movements, etc.), we recognize that frog locomotor repertoire comprises other movements not described here (e.g., scratching, scooping, asynchronous escape on land). These context or stimulus-dependent behaviors could be further explored using a specific stimulation paradigm or in a 3D arena, for example.

## ACKNOWLEDGMENTS

We would like to thank the members of the Sweeney Lab (especially Stavros Papadopoulos and Sophie Gobeil) for their contributions to this project and, in addition to the lab, Graziana Gatto and Mario de Bono, for discussion, and support. We are also grateful to Tom Jessell and Chris Kintner for their scientific insight and mentorship during the conception of this project. This project would also not have been possible with the technical support of the Matthias Nowak, Verena Mayer and the Aquatics as well as the Imaging and Optics Facility support teams (ISTA). In addition, we thank our funding sources for providing the resources to do these experiments: FTI Strategy Lower Austria Dissertation Grant Number FT121-D-046 (D.V.); Horizon Europe ERC Starting Grant Number 101041551 (L.B.S., F.A.T. and D.V); Special Research Program (SFB) of the Austrian Science Fund (FWF) Project number F7814-B (L.B.S); NINDS 5R35NS116858 (J.S.D); CZI grant DAF2020-225401 (DOI): 10.37921/120055ratwvi (R.H.); NIH grant number R01NS123116 (J.B.B); American Lebanese Syrian Associated Charities (ALSAC) (J.B.B.); German Academic Exchange Service (DAAD) IFI Grant Number 57515251-91853472 (Z.H.); and Project A.L.S. (S.B-M.).

## AUTHOR CONTRIBUTIONS

L.B.S. led and coordinated the project. D.V., F.A.T., Z.H., A.T., P.C, M.J.J., M.I.G. and L.B.S. implemented experiments. D.V. and L.B.S. led and collected all *Xenopus* cell type data. F.A.T, Z.H., and M.J.J. collected all behavioral data, in close collaboration with R.H and C.S. who designed the behavioral imaging setup and analysis pipeline, respectively. A.T, P.C., and J.B.B. collected all mouse data. S. B-M. generated antibodies. D.V., F.A.T., C.S. and L.B.S wrote the manuscript. J.D., J.B.B, S. B-M. and M.I.G. edited the manuscript and provided critical feedback on the project.

## TABLES

**Table S1. Primer sequences used for PCR genotyping and TIDE analysis.**

**Table S2. Single guide RNA sequences used to generate CRISPR mutants.**

**Table S3. PCR conditions used for genotyping FoxP1 and En1 genes.**

**Table S4. Metrics of the behavioral tracking models.**

## MOVIES

**Supplemental Movies S1-S7. Examples of tadpole and frog motor behavior at each stage.**

**Supplemental Movie S8. Representative unilateral FoxP1 CRISPR mutant frog.**

**Supplemental Movie S9. Representative unilateral En1 CRISPR mutant frog.**

## EXPERIMENTAL METHODS

### RESOURCE AVAILABILITY

#### Lead contact

Further information and requests for resources and reagents should be directed to and will be fulfilled by the lead contact, Lora B. Sweeney.

#### Materials availability

This study generated several new *Xenopus* antibodies and transgenics lines, listed in the table below. The materials will be made available upon request to the lead contact.

#### Data and code availability

The code used for analysis of the behavioral data is available at Github (https://github.com/sweeneylab/MN_V1_analysis).

### KEY RESOURCES TABLE

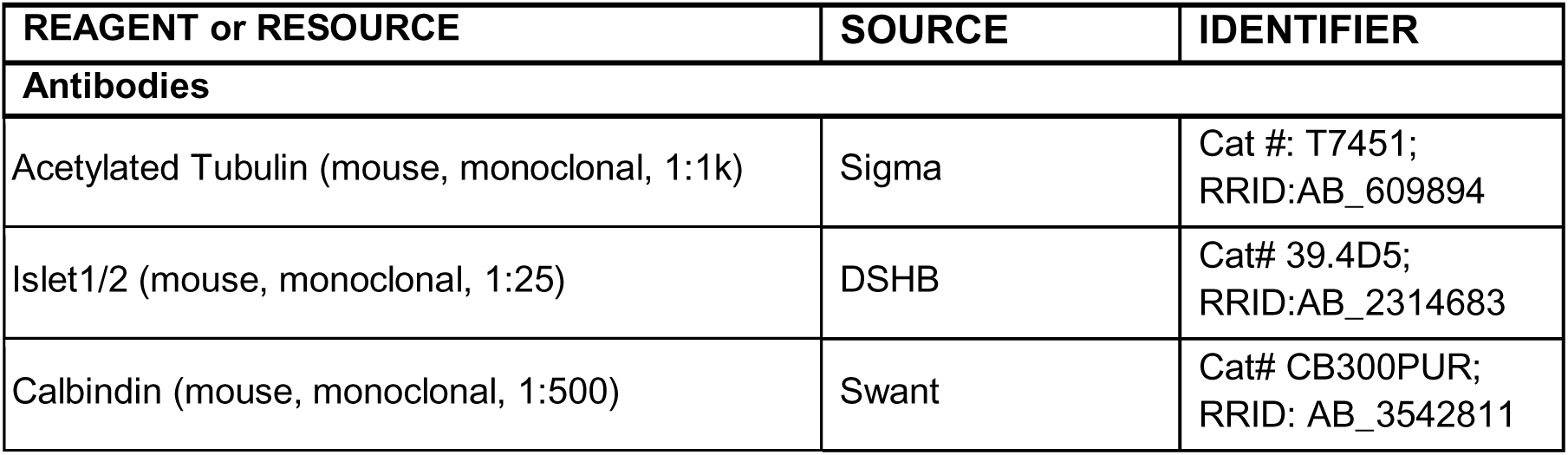
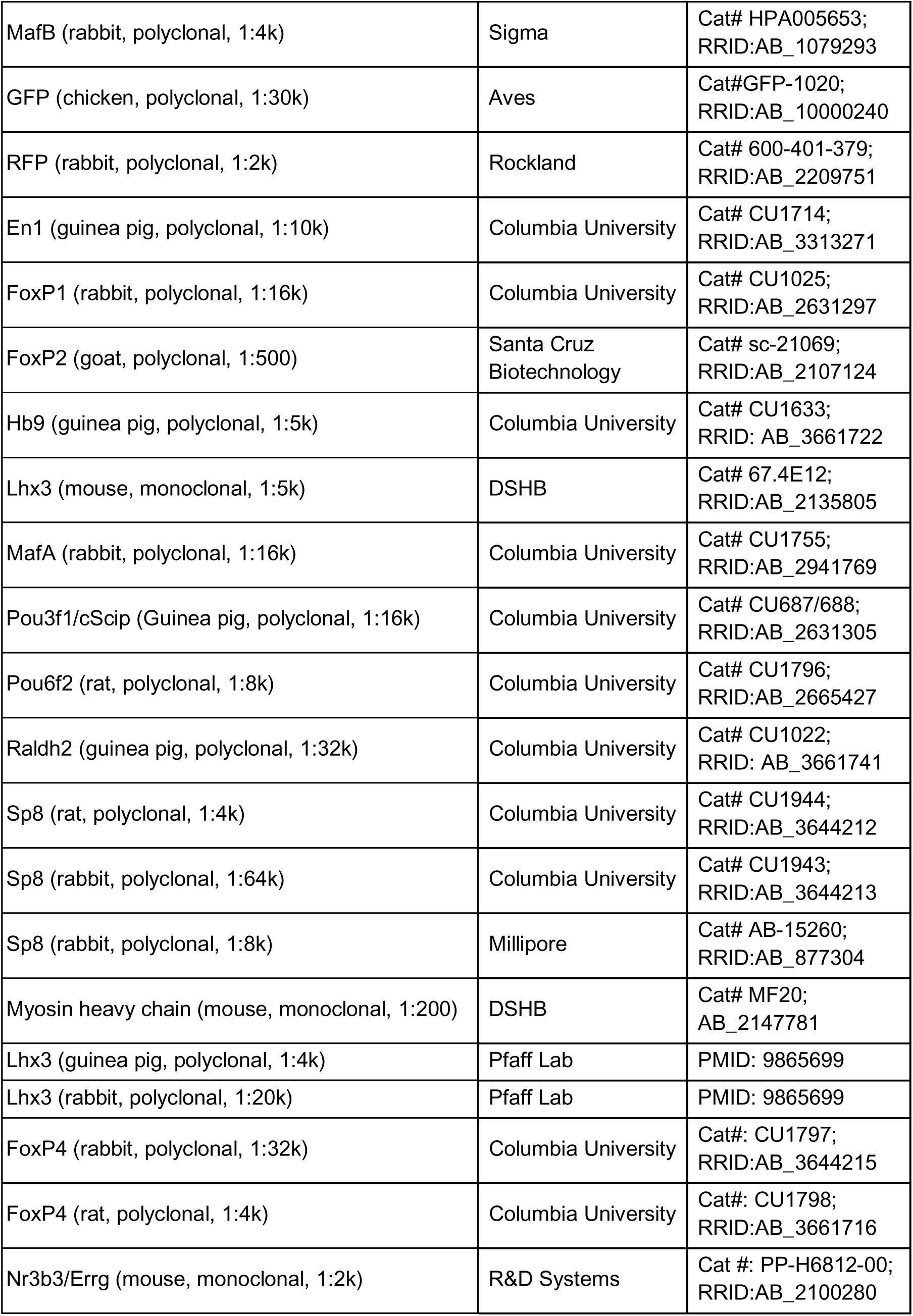
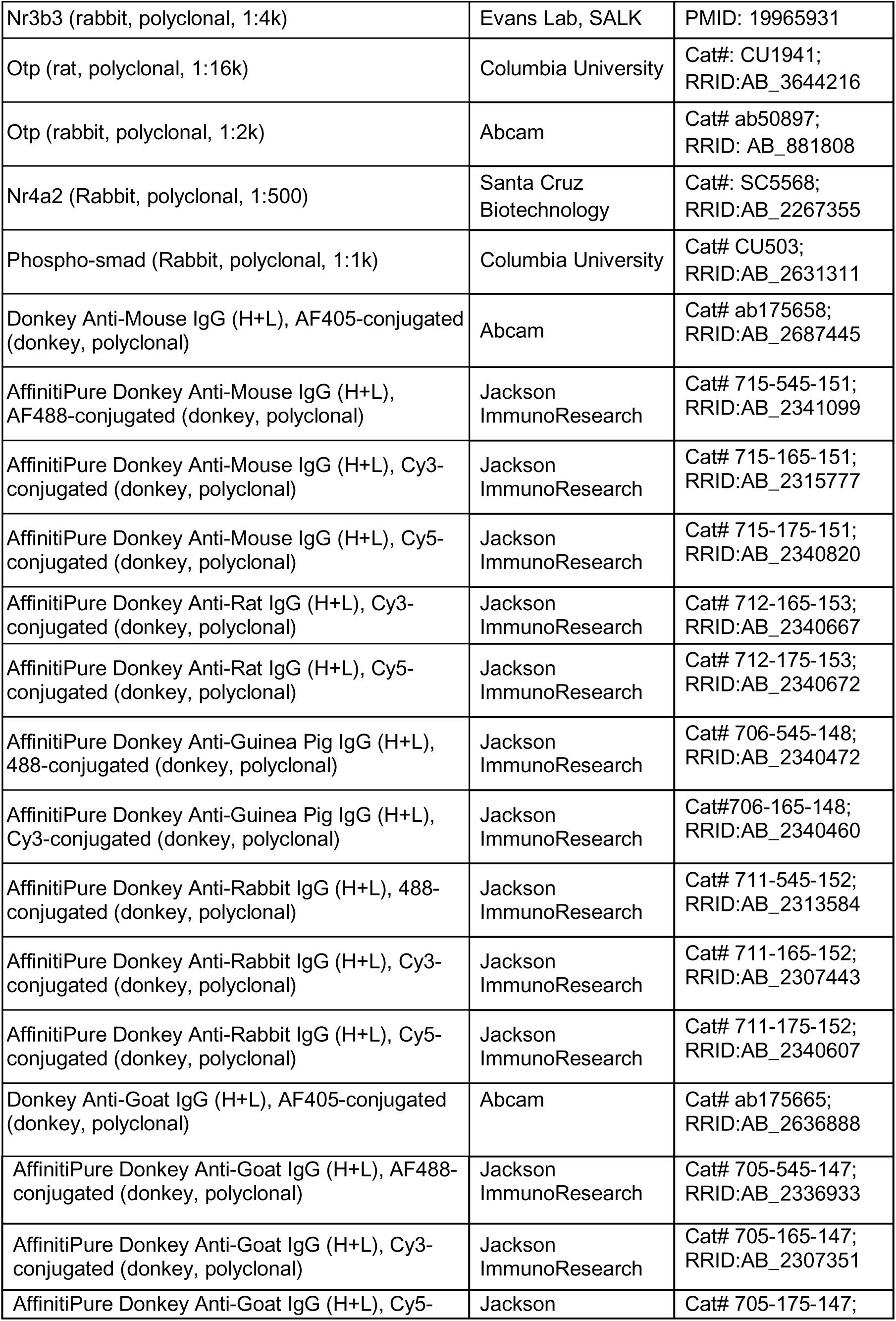
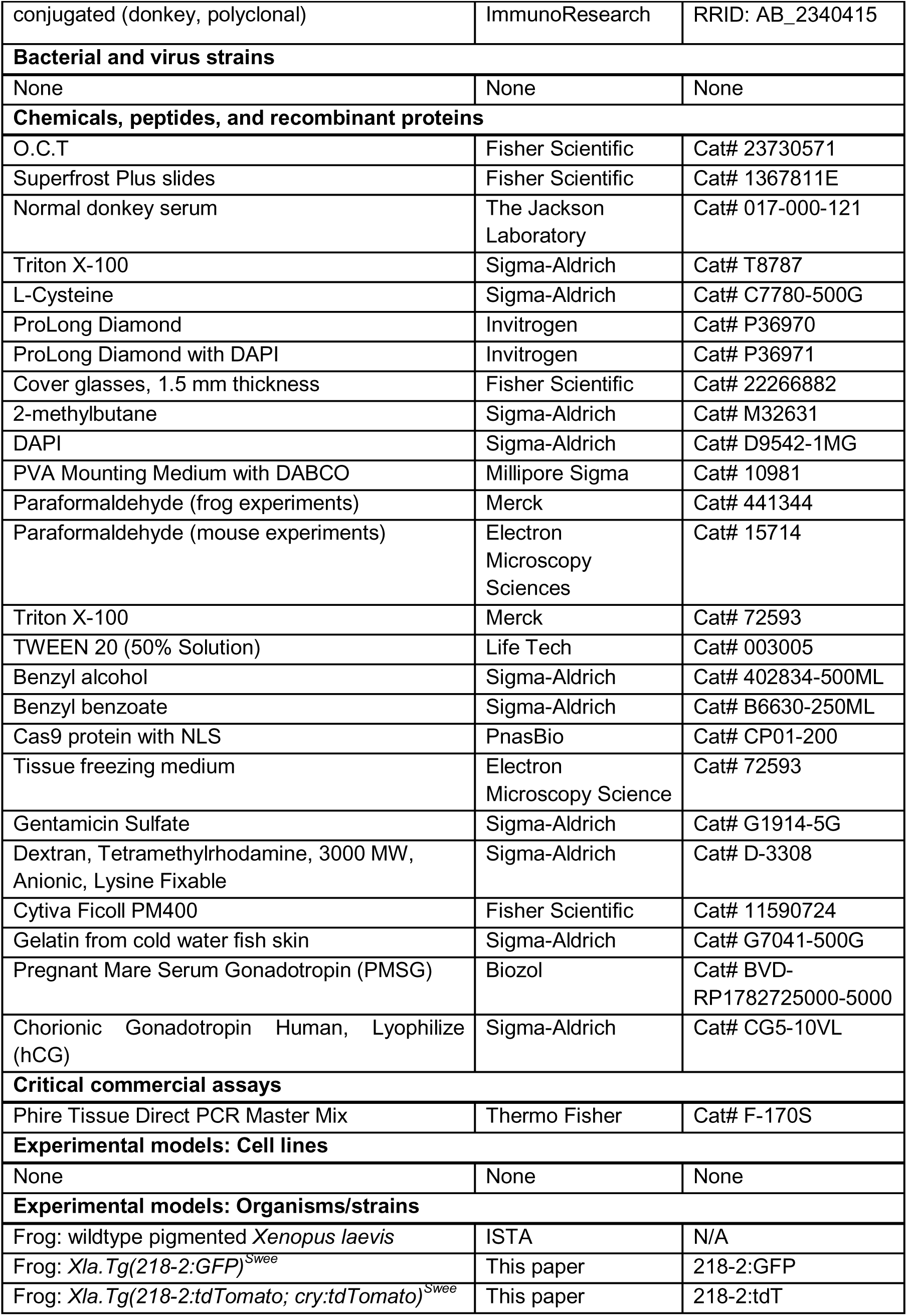
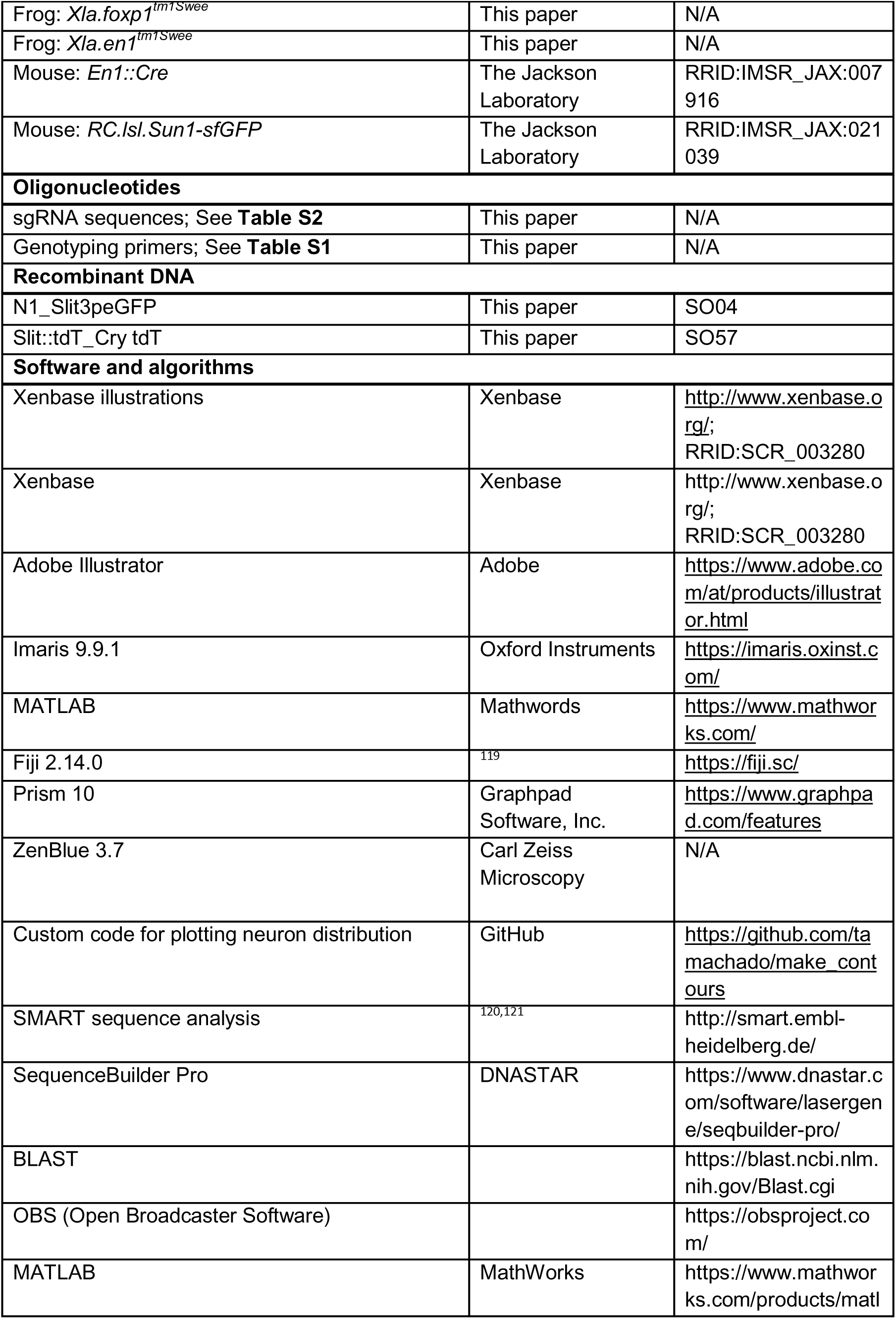
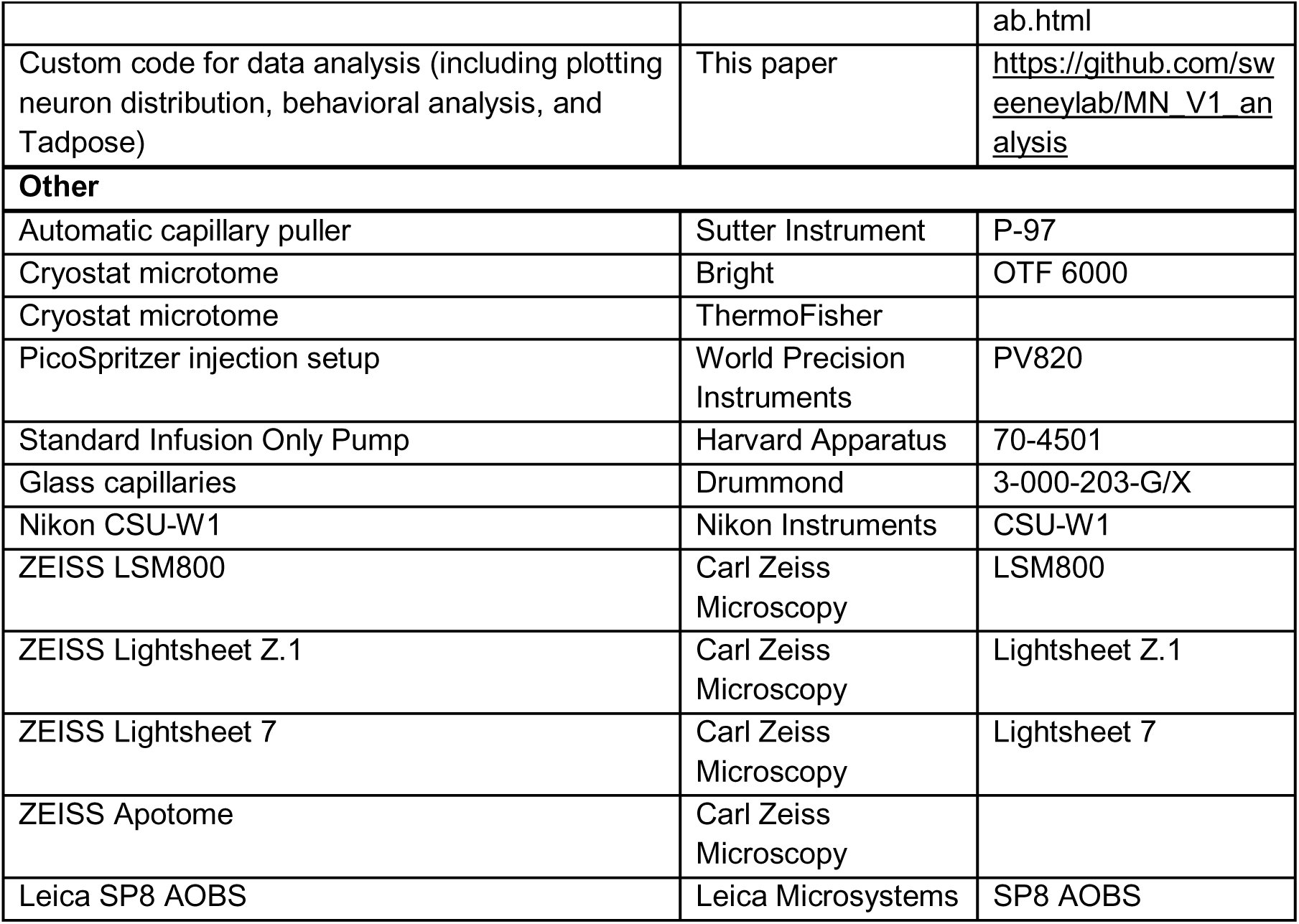

### EXPERIMENTAL MODEL DETAILS

#### Animals

Pigmented *Xenopus laevis* tadpoles were bred and raised at the Institute of Science and Technology Austria. Animals were maintained in specialized frog facilities, with the water temperature kept between 18–22 °C and a 12-hour light:12-hour dark cycle. All experimental procedures and animal husbandry followed protocols approved by the local authorities (permit number 2020-0.550.806, 2020-0.762.370, 2022-0.137.228, 2023-0.591.050 and 2024-0.019.606). Tadpoles and frogs of both sexes were used for experiments indiscriminately, as sex cannot be determined at these stages.

All mouse experiments were performed by the Jay Bikoff and group in accordance with National Institutes of Health (NIH) guidelines and approved by the Institutional Animal Care and Use Committee (IACUC) of St. Jude Children’s Research Hospital. Animals were housed on ventilated racks with controlled temperature and humidity, a 12-hour light/dark cycle, and *ad libitum* food and water. Mice of both sexes were used for experiments.

### METHOD DETAILS

#### Antibodies

The commercially available antibodies were obtained from the sources listed in the reagent table above. Additional antibodies against *Xenopus* proteins were generated in rat and rabbit at Covance using the following peptide sequences: *Xenopus laevis* OTP, CSSPDSSDVWRGSSIASLRRK (C-Term uniprot A0A1L8I213 (L) and A0A1L8HRF6 (S); Entrez gene 108717966 and 108707273); *Xenopus laevis* SP8 CHSPDLLHPPDRNGLE (C-Term uniprot Q5XGT8; Entrez Gene 495143); *Xenopus laevis* FoxP4 VGREGSGSGETNGELNPC (N-Term uniprot Q4VYR7; Entrez Gene 733240).

#### Immunohistochemistry and Microscopy of Frog Tissue

Spinal cord tissue was dissected and fixed in 4% paraformaldehyde (PFA) in phosphate buffer (w/v) for 90 minutes on ice, followed by cryoprotection overnight at 4°C in 15% sucrose-PBS solution containing 8% cold fish skin gelatin (w/v).

For antibody staining of sections, the tissue was embedded in tissue freezing medium, frozen on dry ice, and cryosectioned at 15 µm. Spinal cross sections were dried for 30 minutes at room temperature and then for another 4 hours at 4 °C. Once dry, sections were rehydrated with 1x PBS containing 0.2% Triton (PBST) for 2-5 minutes, washed another time with PBST, and left to incubate in primary antibody/PBST solution overnight at 4°C. Tissue was then washed three times with PBST. Secondary antibody incubation was performed in a dark chamber at room temperature for 30 minutes followed by three PBST washes. Sections were then mounted with 80-100 μL PVA/DABCO, coverslipped, and left to cure at room temperature for at least 2 hours before imaging. Images of spinal sections were acquired at a confocal (ZEISS LSM800), spinning disk (Nikon CSU-W1) or widefield (ZEISS Apotome) microscope using a 20x objective.

For whole-mount immunohistochemistry, tadpoles were fixed in PFA, washed three times with PBS and once with PBS containing 0.1% Tween or 0.2% Triton (PBST). For BABB clearing, tissue was then incubated in the primary antibody mixture at 4 °C for 4–6 days, washed three times in PBST, and incubated in the secondary antibody solution overnight. Tissue was washed in PBST three times and transferred into two parts benzyl benzoate and one part benzyl alcohol until clear. For CUBIC clearing, we adapted the protocol from Ueda Lab ^122^. Briefly, directly after PFA fixation, tissue was washed three times in PBS and incubated in 50% CUBIC-L/R1a solution for three hours at room temperature, followed by incubation in 100% CUBIC-L/R1a for 2-4 days at 37 °C. Tissue was washed 6 times with PBS, and immunostained as described above. Prior to imaging, tissue was transferred to CUBIC-R+(N) for refractive index matching. Cleared tadpoles were imaged using a lightsheet microscope (Figure 2I using ZEISS Lightsheet Z.1; Figure 3E-F using ZEISS Lightsheet 7).

Representative images of the spinal cord were obtained by creating maximum intensity projections from Z stacks using Zen (ZEISS) or Fiji softwares.

#### Immunohistochemistry and Microscopy of Mouse Tissue

Adult pregnant female mice were deeply anesthetized via intraperitoneal injection of Avertin (Tribromoethanol, 240 mg/kg body weight) and then transcardially perfused sequentially with ice-cold PBS followed by 4% PFA (diluted in PBS). Individual embryos were not separately perfused. The spinal column was dissected and post-fixed overnight at 4 °C, followed by cryoprotection in 30% sucrose (w/v) in 0.1M PB for at least 24 hours at 4 °C. Tissue was embedded in O.C.T., frozen, cryosectioned in the transverse plane at 10 microns, and mounted on Superfrost Plus slides. Immunohistochemistry was performed by blocking in 10% normal donkey serum and 0.3% Triton X-100, exposure to primary antibodies (guinea pig anti-En1 diluted at 1:16000 in PBST, and chicken anti-GFP diluted at 1:30000 in PBST) overnight at 4°C, and fluorophore-conjugated (Alexa Fluor Plus 488 or Alexa Fluor Plus 555) secondary antibodies for 1 hour at room temperature. Sections were mounted using ProLong Diamond and coverslipped for imaging (Fisher Cover Glass, Cat# 22266882). Confocal images were acquired on a Leica SP8 AOBS (Leica Microsystems) confocal microscope using a 20x/0.8 NA objective. The positions of En1+ V1 neurons were identified using FIJI cell counter and plotted using custom Matlab scripts (https://github.com/tamachado/make_contours). To account for variation in spinal cord size along the rostrocaudal axis, a representative spinal cord image was first reoriented in FIJI so that the dorsal edge of the spinal cord was parallel with the top of the image. All other spinal cord images were aligned and scaled to the reoriented image by using the FIJI registration plugin (Align Images Using Line ROI). First, a line was drawn between the dorsal and ventral edges of the spinal cord through the central canal. The plugin then uses the location, length, and angle of the obtained line to translate, scale, and rotate all other images.

#### Retrograde Labeling

Prior to retrograde labeling, preoperative analgesia and anesthesia was administered to metamorphic tadpoles. Animals were placed onto a Tricaine-soaked gauze inside a sterile dish. Hindlimbs were severed at the limb base and crystals of Rhodamine Dextran were applied to the exposed area. After crystal application, tadpoles were placed into sterile 0.1x Marc’s Modified Ringers (MMR) (recipe for 10x recipe: 1 M NaCl; 20 mM KCl; 10 mM MgSO_4_·7H_2_O; 20 mM CaCl_2_·2H_2_O; 1 mM EDTA; 50 mM HEPES, pH adjusted to 7.5) supplemented with 50 μg/ml gentamicin. Two days post-administration, tadpoles were fixed for further analysis.

#### *Xenopus* transgenesis

To induce ovulation, *Xenopus* females were pre-primed with 100 units of pregnant mare serum gonadotropin (PMSG; dorsal lymph sac injection) 3-12 days before egg laying, followed by priming with 1000 units of human chorionic gonadotropin (hCG; dorsal lymph sac injection) the day prior to egg collection. On the egg laying day, frogs were transferred into a tank filled with 1x egg laying solution diluted in MilliQ water (recipe for 10x stock: 1.2 M NaCl; 10 mM KCl; 24 mM NaHCO_3_; 8 mM MgSO_4_·7H_2_O; 300 mM; Tris; 4 mM CaCl_2_·2H_2_O; 3 mM Ca(NO_3_)_2_·4H_2_O, pH adjusted to 7.4-7.5 with concentrated HCl). Several hours after, eggs were collected, dejellied using 1x MMR solution containing 3% cysteine (pH 8.2), and subsequently washed twice in 1x MMR. Eggs were then transferred to agarose-coated dishes containing cold injection buffer (6% Ficoll, 0.5x MMR without Ca or EDTA). In parallel, sperm nuclei were prepared following the transplantation protocol outlined in ^123^. Specifically, 4 μl of sperm nuclei preparation was incubated with 2.5 μl of linearized DNA plasmid (100 ng/μl) for 5 minutes at room temperature. This mixture was then combined with 16 μl of sperm dilution buffer and incubated for another 10 minutes at room temperature. The final mixture was diluted by resuspending 6 μl of the prepared sperm nuclei solution in 200 μl of sperm dilution buffer. For nuclear injections, we adapted the method from ^123^. Briefly, glass capillaries were used to introduce the sperm nuclei mixture into unfertilized eggs. Following injection, the eggs were activated for 5 minutes in a calcium-ionophore-containing activation buffer (6% Ficoll, 0.5x MMR). Post-activation, embryos were thoroughly rinsed in post-injection buffer (1% Ficoll, 0.1x MMR). Cleaving embryos were subsequently transferred to 0.1x MMR supplemented with 50 μg/ml gentamicin and incubated at 15 °C until the onset of gastrulation. Around NF40, GFP or tdTomato fluorescence was detectable in the hindbrain and spinal cord, enabling the selection of transgenic embryos. Genotype confirmation was performed via PCR genotyping (See **Table S1** for primer sequences).

#### CRISPR-Cas9 knockout generation

Ovulation was induced as described above. Eggs were collected, rinsed with deionized water and *in vitro* fertilized using freshly dissected and smashed testis in 0.1x MMR. The embryos were then dejellied using 0.1x MMR solution containing 3% cysteine (pH 8.2), and subsequently washed twice in deionized water and twice in 0.1x MMR. Embryos were then transferred to petri dishes containing cold RNA injection buffer (1% Ficoll, 0.5x MMR) on a 16°C cold plate. Embryos were injected with either 3 or 16 ng of purified Cas9-NLS protein diluted in DNA-free water (Synthego), and 5 ng sgRNA (Synthego) either (i) twice into diagonally opposite points of the animal pole between 20 minutes and before cleavage to generate bilateral mutant animals, or (ii) once into only one cell at two-cell stage to generate unilateral mutant animals. Mutant and wild-type embryos were then moved to petri dishes containing 0.1x MMR supplemented with 50 μg/ml gentamicin and kept on a 16°C cold plate at least until the onset of gastrulation.

#### sgRNA design

To design sgRNAs, the mRNA and DNA sequences of both long and short isoforms of the gene of interest were taken from Xenbase. Single guide RNAs (sgRNAs) were designed using *Xenopus laevis* genome versions 9.2 or 10.1 with ChopChop (https://chopchop.cbu.uib.no/). sgRNAs were designed to target a 100% conserved region of the gene of interest in both short and long chromosomes. Additionally, sgRNAs were further selected such that the PAM site aligned with a highly conserved amino acid within this domain. sgRNA were synthesized *in vitro* at Synthego (https://www.synthego.com/). A full list of sgRNA sequences is presented in **Table S2**.

#### *Xenopus laevis* genotyping

To confirm the efficiency of the sgRNAs in generating bilateral and unilateral mutant animals, embryos were genotyped 2 days post fertilization. Each embryo was transferred in a separate Eppendorf tube with 100μl of 50 mM NaOH and boiled at 95-100°C for 15-30 minutes. After briefly vortexing the dissolved embryo, 15μl of Tris pH 8.8 was added and samples were centrifuged at 15,000 rpm at room temperature for 5 min and stored at −20°C. To obtain DNA material from mutant tadpoles and frogs, the animals were first anesthetized by transferring them to a 0.1x MMR with 0.01% tricaine solution and then the tail or the toe was clipped, and the tissue was processed in the same way as described above for the embryos. Mutant tadpoles and frogs were then moved to 0.1x MMR supplemented with 50 μg/ml gentamicin. The tail and hindlimbs of tadpoles and frogs were genotyped only after their final round of imaging.

The extracted DNA was then prepared for PCR, by adding 1x Phire Tissue Direct PCR master mix (Thermo Fisher), the forward and reverse primers necessary to amplify the gene of interest. The PCR primers and reaction conditions are as described in **Table S2** and **Table S3**, respectively. PCR products were Sanger sequenced using the reverse primer and mutation rate was estimated using TIDE analysis software^78^. For juvenile TIDE analysis estimation, the genomic DNA for left and right hindlimbs was processed separately and TIDE results were averaged.

### Behavioral setup and video recording

The recording chamber measures 94 cm (length) by 73 cm (width) by 130 cm (height). For recordings, a 60 frames-per-second high-resolution IDS Imaging uEye+ UI-3180CP-M-GL camera, positioned 35.5 cm perpendicularly above a pull-out transparent acrylic sheet where glass dishes are put for imaging, was used to simultaneously allow for up to 6 dishes (14 cm in diameter) to be imaged in parallel. Infrared light was generated from below by six 850nm infrared LED light sources with two light diffusers (semi-transparent acrylic sheets). Before recording, the lens aperture is adjusted so that all dishes were in focus, all setup specifications and camera calibration files can be found on our github repository.

Before imaging, larvae, tadpoles and frogs were staged according to Nieuwkoop and Faber anatomical criteria^6^. Metamorphic stages were split into seven bins according to their anatomical features (NF37-37, NF44-48, NF52-55, NF57-58, NF59-62, NF63-64 and juvenile, which included animals from NF65 to less than 2.5 cm long froglets). For NF37-38, 5-10 animals were put per dish containing 100 ml of 0.1x MMR, whereas for all other stages only one animal per dish was put in 180 ml of 0.1x MMR. The animals were left for 15 min in the dishes to adapt to the behavioral setup conditions, and were then recorded using OBS Studio software while freely moving for two hours (2 times 1 hour videos per animal). Raw videos containing multiple dishes were cut into videos containing only one dish using a custom-made Matlab script to generate an undistorted image to correct for camera distortion and Fiji to draw a 800×800 pixel region of interest around individual dishes.

Videos were then subject to pose-estimation by the deep learning algorithm SLEAP^8,9^. For each stage bin, two SLEAP models were trained: a centroid model to quantify general features of tadpole-to-frog movement and a centered model to score kinematic features of the limbs and tail (**Figure S1C**). As larval tadpoles were small and largely stationary, multi-animal SLEAP detection and tracking was used at NF37-38 to capture a sufficient number of movement episodes ^27^; single-animal SLEAP pose estimation was used for all other stages^28^. Centroid models for each stage tracked the center point of the body across frames, whereas centered models tracked a fixed number of points along the length of the tail and a single point for each limb joint. For the centered model, the following tail and/or limb points are tracked: eight tail points for NF44-48; 12 tail points for NF52-55; 12 tail, 10 hindlimb points for NF57-58; 12 tail, 10 hindlimb, 8 forelimb points for NF59-62 and NF63-64; 1 tail, 10 hindlimb, 8 forelimb points for juvenile stage.

#### Pose Estimation by SLEAP

To predict the pose of tadpoles and frogs the SLEAP framework was used (ver. 1.2.9). For all stage bins, we applied a multi-animal top-down approach. In the centroid model we applied an input scaling of 0.5 on the gray-value movies. For training, an UNet backbone with max-stride of 16 and output-stride of 2 was selected. The Gaussian sigma for the central body-part was set to 3 (px). The number of filters was 16 with a filter-rate of 2. We used a middle block and up-interpolation. For training data augmentation random rotation, horizontal flips, scale and brightness with their corresponding default parameters were chosen. We trained for 400 epochs with a batch size of 8 and plateau patience of 40.

The centered-instance model was trained with no input scaling using an UNet backbone with max-stride of 32 and output-stride of 1 was selected. The number of filters was 32 with a filter-rate of 2. The Gaussian sigma for the body-parts was set to 5 (px). We used a middle block and up-interpolation. For training data augmentation rotation, random horizontal flips, scale and brightness with their corresponding default parameters were chosen. We trained for 800 epochs with a batch size of 8 and plateau patience of 60. In addition, online mining with a minimum of 2 hard key-points was selected. We used 10% of the training frames for validation and only the best model regarding the validation error was kept after training.

We manually annotated 448 frames for NF37-38, 607 frames for NF44-48, 451 frames for NF52-55, 405-499 frames for NF57-58, 405-457 frames for NF59-62, 250 frames for NF63-64, 405-410 frames for juvenile; the training was done on an AMD Ryzen 5950X 16-Core workstation equipped with an Nvidia A4000 GPU having 16 GB of VRAM.

To track the detected poses of tadpoles and frogs, we used SLEAP’s default centroid tracker using the Hungarian method for instance matching in a frame window of size 5.

The prediction and tracking of the recorded videos has been done with the SLEAP framework (ver. 1.2.9/1.3.3) on the institute SLURM HPC cluster using a custom submission script. The SLEAP output was converted to HDF5 for down-stream processing.

Our models achieved high levels of pose estimation accuracy as measured by quality control metrics such as average distance, visual recall, visual precision, object keypoint similarity (OKS) and mean precision per keypoint (mPCK) (**Table S4**). The average distance metric indicates the localization accuracy of predicted and manually annotated key points across the validation set. The overall keypoint detection accuracy is reflected by the Visual Recall and Visual Precision metrics. OKS metrics indicated the accuracy of the model overall, with higher numbers indicating better accuracy. Finally, mPCK metric provides information about the accuracy of the model to predict individual keypoints and is calculated as a percentage of predicted key points that are within 5 pixel from the manually annotated ones.

#### Quantification of Behavior

##### Locomotion Analysis

Locomotion features extract information about the basic movement of an SLEAP-tracked body-part. We partitioned the time-course of a tracked animal into *moving* and *non-moving episodes*. First, we extracted the xy-locations of the central body part from the SLEAP analysis HDF5 file. If the body-part was not detected in all frames, we linearly interpolated the missing frame locations using `numpy.interp`. Next, we applied a Gaussian smoothing to the x and y body-part locations using `skimage.filters.gaussian`, with the sigma set to 1. The resulting smoothed locations are used to compute the instantaneous speed by central differences with `numpy.gradient`. We then smoothed the resulting speed with a Gaussian with sigma=30. Each frame is associated with a speed in pixels per frame, which we calibrated to cm per sec using the known diameter of the dish and the camera speed. Each dish was manually annotated with a circle ROI in ImageJ/Fiji. Finally, we thresholded the calibrated speed with a manually chosen threshold of 1.2 cm/sec to obtain a binary partitioning into moving vs. non-moving animal episodes.

To compute the instantaneous speed and acceleration the central difference was used. For acceleration only the forward (positive) acceleration values were considered. In addition to the time spent moving per hour and the total distance, we computed various statistics of the instantaneous speed, acceleration and directional change.

To measure directional change, the angle between two succeeding time-points was computed. The angle at time *t_i_* is defined by the locations *P_i_* of the selected body-part. The angle was computed from two segments 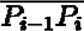 and 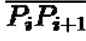. The angle is 0 if the segments are parallel and ranges from −180 to 180 degrees. To get a more robust signal to noise in the body part localizations, the raw xy-locations were smoothed with a Gaussian with sigma=1. In addition, the time was sub-sampled using a factor of 8. All angles were given in degrees.

To calculate the area explored by each animal, we measured the ratio of dish area explored by the frog by selecting a central SLEAP-tracked body-part and building the 2D histogram of its trajectory. The square containing the circular dish ROI was discretized into 128 x 128 2D bins of size 1.09 x 1.09 mm. The 2D location histogram of the selected body-part is built over these bins. Each bin contains the number of times the location of the body-part was overlapping with this bin over its trajectory. Hence, a bin with value zero was never *visited*. The ratio of area explored is built by the number of bins visited divided by the total number of bins. To make movies of different length comparable, we further normalize this ratio to per hour.

| Feature | Description |
| --- | --- |
| speed_moving_mean | mean of the instantaneous speed while moving (cm/sec) |
| time_spend_moving_percentage | ratio of moving and the total number of frames (au) |
| directional_change_mean | mean of the directional change (degree) |
| directional_change_pos_mean | mean directional change regarding positive angles only |
| directional_change_neg_mean | mean directional change for regarding negative angles |
| acceleration_mean | mean positive acceleration |
| total_distance | total distance traveled while moving |

#### Angle range and correlation analysis

Angle range features extract statistics of angle distributions at selected body-part segments. We defined several body-part segments where angles are computed in range (−180, 180) degrees, where an angle of 0 degree indicates parallel segments. Statistics of the angle ranges are partitioned from moving and non-moving episodes.

Angle Correlation features extract the correlation of angles measured at two pairs of body-part segments in a time resolved manner. The correlation was computed as windowed Pearson correlation. From the correlation coefficient distribution (during *active episodes*) several statistics were extracted.

First, angles at two pairs of body-part segments are computed. Both series of raw angles A and B were first smoothed with a Gaussian of sigma = 1 using `scipy.ndimage.gaussian_filter1d` function. Then, the z-score of the smoothed angles values were computed. The z-score subtracts the mean and divides by the standard deviation to center the angle values around zero with unit variance.

To define *active episodes*, where the angle series have enough variance to compute meaningful correlations, we computed the gradient magnitude of the smoothed angle z-scores for A and B using central differences with `numpy.gradient`. The gradient magnitude involves a second Gaussian smoothing of sigma=15.

The maximum over both gradient magnitudes were computed and thresholded with 5.73. Only correlations from time frames exceeding this threshold were used to compute the angle correlation distribution.

The correlation is computed by a centered, rolling Pearson cross-correlation using a `pandas.rolling(win, center=True).corr()` where inputs are the smoothed and z-scored input angle time series. From the resulting correlation distribution ranging ∈ [-1,1] several statistics were extracted. We used a window size of win=31 (frames) for the windowed cross-correlation.

| Feature | Description |
| --- | --- |
| angle_moving_std | std of angles during moving episodes (degree) |
| angle_moving_mean | mean of angles during moving episodes |
| corr_median | median of the correlation distribution |

##### Frequency Analysis

Frequency features extract information about frequency of angle changes at specified body-part segments. The frequency is estimated using continuous Wavelet transform using the python module PyWavelets (ver. 1.4.1). Frequency bins (N=24) were chosen in the range [0.937, 30] Hz with each bin being a factor of 1.16263 higher than the previous bin, and 30 Hz is the Nyquist limit for the recordings with a camera frame rate of 60 Hz.

We used the complex valued Morlet (Gabor) wavelet function for frequency estimation cmorl1.5-1.0, where cmorB-C with floating point values *B* = 1.5*, C* = 1.0 is given by:

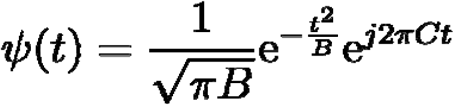

where *B* is the bandwidth and *C* is the center frequency.

We defined several body-part segments, from which angle frequencies are estimated and statistics of the power spectrum density (PSD) are extracted during *active episodes*.

As the tail mean power spectrum displayed a bimodal distribution, the frequency spectra were subdivided into low (0.9 −4.5 Hz) and high (4.5 −20 Hz) bins to estimate movement within each bin more accurately.

To define active episode, all angles are preprocessed by slightly smoothing with a Gaussian with sigma = 0.1, followed by computing the z-score.The preprocessed angle values are first smoothed with a Gaussian of sigma=15 using `scipy.ndimage.gaussian_filter1d` function. Then, the gradient magnitude is computed using central differences with `numpy.gradient`. The resulting magnitude is thresholded with 5.73. Only frequency estimates (mean power spectral density) from time frames exceeding this threshold are used to compute the mean PSD distribution.

To estimate angle change frequencies, we applied a continuous wavelet transform implemented in the PyWavelets function `pywt.cwt`. To obtain the power spectral density (PSD), we computed the magnitude by multiplying the wavelet coefficients with their complex conjugate. The time-resolved PSD is averaged in *active episodes* as described above.

Peak finding on the mean power spectral density is applied using the function `scipy.signal.find_peaks` to obtain the dominant frequency bin. The dominant frequency hence corresponds to a local maxima in the mean power spectral density (PSD). The prominence of a peak at the dominant frequency bin measures how much a peak *stands out* from the surrounding baseline of the signal and is defined as the vertical distance between the peak and its lowest contour line. The analysis is repeated for frequency ranges below and above a manual set frequency threshold of 4.5 Hz.

To remove spurious low frequency content, we optionally applied background subtraction to the smoothed angle z-scores. The background is estimated by smoothing the signal with a Gaussian of sigma=15.

| Feature | Description |
| --- | --- |
| `dominant_freq` | dominant frequency: local maximum frequency |
| `dominant_freq_prominence` | prominence of the dominant frequency peak |
| `dominant_freq_power` | mean PSD in active episodes at the `dominant_freq` |
| `dominant_freq_X-` | same as above, for frequencies < 4.5 Hz |
| dominant_freq | dominant frequency: local maximum frequency |
| dominant_freq_prominence_X- | same as above, for frequencies < 4.5 Hz |
| dominant_freq_power_X- | same as above, for frequencies < 4.5 Hz |
| dominant_freq_X+ | same as above, for frequencies > 4.5 Hz |
| dominant_freq_prominence_X+ | same as above, for frequencies > 4.5 Hz |
| dominant_freq_power_X+ | same as above, for frequencies > 4.5 Hz |
| sum_psd_X- | sum of mean PSD for frequencies < 4.5 Hz |
| sum_psd_X+ | sum of mean PSD for frequencies > 4.5 Hz |

##### Animal pose visualization using PCA

To visualize typical configuration of frog body-part poses and their variance, we employed unsupervised dimensionality reduction using Principal Component Analysis (PCA). First, we aligned the animal pose into ego-centric coordinates. We centered the body-part *Tail_1 or Tail_Stem* as the new origin and rotationally aligned body-part *Heart_Center* with the positive y-axis using the Euclidean transformation implemented in the Python library scikit-image (ver. 0.22). Then, from the aligned coordinates a minimum 8000 skeletons body-parts of interest are sampled randomly and normalized to zero mean and unit variance for the PCA analysis. We used the first principal component to color-code the line segments of randomly sampled the leg or tail poses.

### QUANTIFICATION AND STATISTICAL ANALYSIS

#### Behavioral tracking statistics

Statistical quantification of the behavioral data obtained from our Tadpose analysis was done only on videos with a minimum of 90% tracked frames and considering only moving episodes for overall animal movement and active episodes for tail and limb kinematics quantification.

ROUT outlier test was first applied to each dataset and outlier values removed. Then, each dataset was checked for normal distribution using D’Agostino & Pearson test, Anderson-Darling test, Shapiro-Wilk test, Kolmogorov-Smirnov test as appropriate. Data is assumed to not have the same variance. For comparison between two datasets, if the data was normally distributed unpaired t-test with Welch’s correction was applied, otherwise Mann-Whitney test. For comparison between three or more datasets, if the data was normally distributed Brown-Forsythe and Welch ANOVA test, otherwise Kruskal-Wallis test.

All plots for percentage of time moving, distance traveled, speed, acceleration and turning, as well as mean and std angle, dominant frequency and sum power are shown as median with min and max values as dots, and first quartile and third quartile as box limits. Dots represent a bout of animal movement for NF37-38, as everytime the animal touches the border of the dish it is considered as a new animal. For NF44 to juvenile stage animals, as each animal is recorded for two one-hour windows, two dots represent the same animal. For the limb kinematics analysis of wildtype and En1 ½ CRISPR movement, each animal during one-hour recording was sampled twice with left and right fore and/or hindlimb movement recorded separately and used as an independent data point. For FoxP1 ½ CRISPR analysis, left and right limbs were sampled separately. Mean power spectrum curves are plotted as the median between all data points for frequency bins sampled.

#### Cell-Type Quantification

Motor neurons and V1 interneuron (sub)populations were quantified in *Xenopus* using the spot detection function on full z-stacks within Imaris software. For marker co-expression analysis, a custom MATLAB script (https://github.com/sweeneylab/MN_V1_analysis) was used to identify colocalized nuclei within a 4.5 μm radius. These detection counts were then used to calculate the percentage of a given cell subset within the population. As a comparative reference, motor neuron counts for E13.5 mice^64^ and V1 interneurons counts from P0 mice^16^ were used.

#### Spatial Plotting

During Imaris-based cell detection, three anatomical reference points were annotated in the spinal cord: the center (central canal), the bottom (most ventral point of the white matter), and the side (most lateral point of the white matter). These reference points enabled normalization of the detected cell positions, which were then plotted using a MATLAB script (https://github.com/sweeneylab/MN_V1_analysis) onto a spatially normalized P0 mouse spinal cord at either thoracic or lumbar levels.

#### Entropy Analysis

To define and estimate the diversity and specialization of transcriptomes and gene specificity across developmental stages, we define transcriptome diversity using concepts of Shannon entropy of a gene frequency distribution at each developmental stage, based on the work of Martinez and Reyes-Valdez^71^. Gene specificity is defined as the mutual information between the tissues and the corresponding transcript. More specifically, we quantify transcriptome diversity using 9 TFs: FoxP1, FoxP2, FoxP4, Mafb, Nr3b3, Nr4a2, Otp, Pou6f2, Sp8. We created a measure Dj, of diversity in tissue j, which measures how much a given stage j departs from the transcriptomic distribution of all the stages under comparison. Dj = Hrj - Hj, where Hrj is the average log2 of the global transcript frequencies in a given tissue, and Hj represents an adapted entropy. They are calculated from the relative frequency of cells expressing each i TFs at stage j, pij, and the average frequency across stages pi, by the following expression: Hj = -\sum_i=1^G pij log(pij) and Hrj = -\sum_i=1^G pij log(pi).

#### Statistical Analysis

Cell-type data are presented as mean ± SEM in all graphs unless otherwise specified in Figure Legends. The normality of data distribution was assessed using the Shapiro-Wilk test. For normally distributed data, group differences were analyzed using a Student’s t-test or Analysis of Variance (ANOVA), with Tukey’s post hoc test applied for multiple comparisons when appropriate. In cases where data were non-normally distributed, the Kruskal-Wallis test was used, followed by Dunn’s multiple comparisons test as necessary. All statistical analyses were performed using GraphPad Prism software.

