## Supplemental Information for "Spinal neuron diversity scales exponentially with swim-to-limb transformation during frog metamorphosis"

**Figure S1. Extended analysis of tail and limb movement during *Xenopus* frog metamorphosis, related to Figure 1.**

**A-D. Workflow of SLEAP-based behavioral tracking** including imaging setup (**A**), video processing pipeline (**B**), and centroid (**C**) and centered (**D**) SLEAP models for quantification of animal or tail/limb movement, respectively.

**E-G. Low and high frequency tail movement across metamorphosis.** The dominant low frequency of the tail tip is largely constant with an increase from NF52-55 to NF63-64 (**E**; NF52-55 versus NF63-64, p = 0.001). The amount of tail tip movement in the high frequency bin, represented by the sum power, peaks at NF57-58 and then decreases until NF63-64 (**F**; for NF44-48 versus NF52-55 and NF44-48 versus NF57-58, p = <0.0001; NF52-55 versus NF57-58, p = 0.049; NF57-58 versus NF59-62, p = 0.013; NF57-58 versus NF63-64, p = 0.002). Loss of dominant high frequency at the tail tip from NF44-48 to NF57-58 with no animals displaying a high dominant frequency, and thus no data point, at NF63-64 (**G**; for NF44-48 versus NF52-55 and NF44-48 versus NF57-58, p = <0.0001; NF52-55 versus NF57-58, p = 0.002).

**H-J. Extended analysis of the gain of hindlimb movement across frog metamorphosis.** From NF57-58 to NF63-64, the knee displays an increased mean angle when moving (**H**; for NF57-58 versus NF59-62, NF57-58 versus NF63-64 and NF59-62 versus NF63-64, p = <0.0001). The amount of movement of the knee in the low frequency bin, represented by the sum power, increases from NF57-58 to NF63-64 (**I**; for NF57-58 versus NF59-62, NF57-58 versus NF63-64 and NF59-62 versus NF63-64, p = <0.0001). The dominant frequency of knee movement also increases from NF57-58 to NF59-62 to juvenile stage, reaching an average of 2.2 Hz (**J**; for NF57-58 versus NF59-62 and NF57-58 versus NF63-64, p = <0.0001; NF59-62 versus juvenile, p = 0.017).

**K-P. Gain of forelimb movement across frog metamorphosis.** PCA plots represent the position of the forelimb and its range of movement during 256 random frames and show an increase in the range of movement from NF59-62 to juvenile stage (**K**; shoulder, yellow; elbow, orange; wrist, red). Scale bar in **K** indicates the color-code of the first principal component of variation of the aligned forelimb positions. Quantification of the mean angle of the elbow shows an increase across metamorphosis (**L**; for NF59-62 versus NF63-64, NF59-62 versus juvenile, and NF63-64 versus juvenile, p = <0.0001). The range of elbow movement decreases from NF59-62 to juvenile (**M**; NF59-62 versus juvenile, p = 0.046; NF63-64 versus juvenile, p = 0.002). Mean power spectrum of the elbow oscillations for each stage of metamorphosis shows a single peak in the low frequency range (**N**; 0.9-4.5 Hz, dark gray). The amount of movement of the elbow in the low frequency bin, represented by the sum power, increases from NF57-58 to juvenile stage (**O**; for NF59-62 versus NF63-64 andNF59-62 versus juvenile, p = <0.0001). The dominant frequency of elbow movement also increases from NF57-58 to juvenile stage, reaching an average of 2 Hz (**P**; NF59-62 versus NF63-64, p = 0.031; NF59-62 versus juvenile, p = <0.0001).

n = 172 animals for NF37-38; n = 47 animals for NF44-48; n = 24 animals for NF52-55, n = 11 animals for NF57-58, n = 13 animals for NF59-62, n = 8 animals for NF63-64, n = 13 animals for juvenile stage.

**Figure S2. Rostrocaudal tail and proximodistal limb movement analysis across metamorphosis, related to Figure 1.**

**A-I. Range and frequency of movement along the rostrocaudal axis of the tail.** SLEAP skeleton (yellow) superimposed onto an image of a recorded animal at NF44-48, NF52-55 and

NF63-64 with all tracked points indicated (**A**; tail top, dark blue; tail mid, blue; tail tip, light blue). The range of movement is initially uniform for all three tail points at NF44-48; however, from NF52-55 to NF63-64, they diverge, with, for example, the tail tip having a greater range than the top (**B**. NF52-55: for top versus tip and mid versus tip, p = <0.0001. NF57-58: for top versus tip and mid versus tip, p = <0.0001. NF59-62: top versus tip, p = 0.006. NF63-64: top versus tip, p = <0.0001; top versus mid, p = 0.024). At NF57-58, when the knee starts participating in movement, it displays a similar low dominant frequency as the most rostral tail point, the top (**C**; tail tip versus knee, p = 0.029; tail mid versus knee, p = 0.118). Whereas, at NF63-64, time point of tail recession, the tail top most closely matches the low dominant frequency of the knee, followed by the tail tip (**D**; tail tip versus knee, p = 0.082; tail mid versus knee, p = 0.023). Mean power spectrum of oscillations at the tail top (circle, dark shade), mid (square, medium shade), and tip (triangle, light shade) for NF44-48 (**E**), NF52-55 (**F**), NF57-58 (**G**), NF59-62 (**H**), and NF63-64 (**I**) with low (0.9-4.5 Hz, dark gray) and high (4.5-20 Hz, light gray) frequency bins highlighted. From NF44-48 to NF59-62, all tail points display bimodal frequency spectra with a peak in the low and high frequency bins (**E-H**); while at NF63-64, they all show unimodal frequency spectra with only one low frequency peak (**I**). At NF44-48 (**E**), the top, mid and tip are similar in their frequency distribution; at all other stages (**F-I**), the power at the tail top is greater across the spectrum than the mid and tip.

**J-Q. Range, coordination, and frequency of movement along the proximodistal axis of the hindlimb.** When moving, the ankle and foot display an increased mean angle from NF57-58 to juvenile, while the mean angle of the hip first decreases until NF63-64 and then increases at juvenile stage (**J**. Hip: for NF57-58 to NF59-62 and NF63-64 versus juvenile, p = <0.0001; for NF57-58 versus juvenile, p = 0.0009. Ankle: NF57-58 to NF59-62, p = 0.0008; NF57-58 versus juvenile, p = <0.0001; NF59-62 versus NF63-64, p = 0.002; NF63-64 versus juvenile, p = 0.007. Foot: for NF57-58 to NF59-62, NF57-58 versus juvenile, NF59-62 versus NF63-64 and NF63-64 versus juvenile, p = <0.0001). The foot never reaches the same angle as the hip and ankle (**J**). From NF57-58 to NF59-62, only the range of ankle movement increases, while hip and foot remain unchanged. From NF59-62 to juvenile stage, all joints show a decrease in range of movement (**K**. Hip: for NF57-58 to juvenile and NF59-62 versus NF63-64, p = <0.0001; NF63-64 versus juvenile, p = 0.005. Ankle: for NF57-58 versus juvenile, NF59-62 versus NF63-64 and NF59-62 versus juvenile, p = <0.0001; for NF57-58 versus NF59-62 and NF63-64 versus juvenile, p = 0.0003. Foot: for NF57-58 versus juvenile, NF59-62 versus NF63-64 and NF59-62 versus juvenile, p = <0.0001; NF63-64 versus juvenile, p = 0.0005). Left-right coordination between knee, ankle and foot increases across metamorphosis, beginning with random bilateral movement at NF57-58 and gaining synchrony by NF63-64 (**L**; +1 = synchronous, 0 = random, -1 = alternating. Hip: NF57-58 versus NF59-62, p = 0.002; NF57-58 versus NF63-64, p = <0.0001; NF59-62 versus NF63-64, p = 0.0004. Ankle: NF57-58 versus NF59-62, p = 0.003; NF57-58 versus NF63-64, p = <0.0001; NF59-62 versus NF63-64, p = 0.004. Foot: NF57-58 versus NF63-64, p = <0.0001; NF59-62 versus NF63-64, p = 0.001). The foot never reaches the same level of bilateral synchronous movement as the hip and ankle (**L**). Mean power spectra of hip (circle, dark shade), knee (square, medium dark shade), ankle (triangle, medium shade) and foot (rhombus, light shade) oscillations from NF57-58 (**M**), NF59-62 (**N**), NF63-64 (**O**), to juvenile stage (**P**) show a similar unimodal distribution with only a low frequency peak. For all joints, the dominant low frequency (dotted black lines) increases across metamorphosis from ~1.5Hz to ~2.2Hz at NF63-64 (**M-P**). The amount of movement of the hip and foot in the low frequency bin, represented by the sum power, increases across metamorphosis reaching the peak at NF63-64 (**Q**. Hip: NF57-58 versus NF59-62, p = 0.001; for NF57-58 versus NF63-64 and NF59-62 versus NF63-64, p =

<0.0001. Foot: NF57-58 versus NF59-62, p = 0.002; for NF57-58 versus NF63-64 and NF59-62 versus NF63-64, p = <0.0001).
n = 172 animals for NF37-38; n = 47 animals for NF44-48; n = 24 animals for NF52-55, n = 11 animals for NF57-58, n = 13 animals for NF59-62, n = 8 animals for NF63-64, n = 13 animals for juvenile stage.

**Figure S3. Motor neuron subtypes in developing *Xenopus* tadpoles, related to Figure 2.**
**A.** Spinal cross sections of NF44-47 tadpoles showing motor neuron (Isl1/2 and Hb9), medial motor column (MMC; Lhx3) or lateral motor column (LMC; FoxP1) markers at the brachial level.
**B.** Spinal cross sections of NF44-47 tadpoles showing motor neuron (Isl1/2 and Hb9), medial motor column (MMC; Lhx3), lateral motor column (LMC; Raldh2) or preganglionic column (PGC; FoxP1, P-Smad) markers at the thoracic level.
**C.** A schematic showing separation of mouse limb motor neurons at the brachial level into a medial and lateral division, LMCm and LMCl, respectively, and pools innervating distinct muscle groups in the forelimb.
**D-E.** In NF54 tadpoles, the LMC is divided into LMCm and LMCl divisions (**D**) and motor pools (**E**) distinguishable by transcription factor expression.
**F**. In NF54 tadpoles, P-Smad and Isl1/2 co-staining marks a preganglionic column at the thoracic level.
**G.** LMC is not present in NF35 tadpoles as shown by Isl1/2 and Raldh2 co-staining. Shown is mean ± SEM (n = 4 animals).
**H-J.** Percentage of motor neurons belonging to each motor column in axial spinal cord at NF35-38 (**H**), limb (**I**) and thoracic (**J**) levels at NF44-47. Shown is mean ± SEM (n = 2–14 animals).
All images represent 15 µm cross sections.

**Figure S4. Timeline of limb and spinal cell type development in *Xenopus laevis,* related to Figures 2, 4, and 6.**
**A.** Schematic representation of the forelimb (FL) and hindlimb (HL) at NF48-57.
**B.** Immunostaining of motor neurons marked by Hb9 and motor columns marked by FoxP1 (lateral motor column, LMC) or Lhx3 (medial motor column, MMC) at the lumbar level.
**C-D.** Immunostaining of V1 interneurons marked by En1 and V1 subsets marked by FoxP1, FoxP2 or Pou6f2.
**E-K.** Quantification of the number of all motor neurons (**E**), MMC (**F**) or LMC (**G**) motor neurons, V1 interneurons (**H**), $V1^{FoxP2}$ (**I**), $V1^{FoxP2}$ (**J**) $V1^{Pou6f2}$ (**K**) at the lumbar level. Shown is mean ± SEM for n = 2 animals.
All images represent 15 µm cross sections. Scale bar, 50 µm. Drawings modified from Xenbase[118] and Xenopus illustrations © Natalya Zahn (2022)[117].

**Figure S5. Expression of En1 is maintained in lineage-traced V1 interneurons during mouse development, related to Figures 4 and 6.**
**A.** Lumbar cross section of *En1::Cre; RC.lsl.Sun1.sfGFP* e13.5 spinal cord showing immunodetection of En1 protein (red) in a subset of lineage-traced V1 interneurons (green)
**B-C.** Spatial distributions of the parental V1 population (green; **B**) and neurons actively expressing En1 (red; **C**).

**Figure S6. V1 subtype diversity emergence during metamorphosis, related to Figure 5.**
**A-C.** Spinal cross sections showing transcription factor expression at larval (**A**), free-swimming (**B**), and limb-circuit stages (**C**) of *Xenopus* development. Tadpole drawings adapted from Xenopus illustrations © Natalya Zahn (2022)[117].
**D.** Quantification of percentage of V1s expressing a given transcription factor in axial (NF35-38), thoracic (NF44-47 and NF54-55) or lumbar spinal cord (NF54). Shown is mean ± SEM for = 4–10 animals.
**E-H.** Quantification of percentage of V1s expressing a given transcription factor at the corresponding stages and levels shown in **A-C**. Shown is mean ± SEM for = 4–10 animals.
All images represent 15 µm cross sections.

**Figure S7. Molecular and spatial organization of V1 subsets along rostrocaudal axis of frog and mouse spinal cord, related to Figure 6.**
**A-C**. Immunoreactivity against V1 marker En1 (red) and subtype markers FoxP2 (green) and MafB (blue) at the brachial (**A**), thoracic (**B**), and lumbar (**C**) segment of the NF54-55 frog. The distribution of En1+ cells for each level is shown in **A'-C'**.
**D**. Number of V1s per spinal level (brachial, thoracic and lumbar) in the NF54-55 frog (black) and the P0 mouse (gray) per 15 µm ventral horn (mean ± SEM for n = 2–4 animals).
**E**. Percentage of V1s expressing a given subset transcription factor (TF) in the frog (black) and mouse (gray) thoracic spinal cord.
**F**. Fold change of the percentage of V1s co-expressing one TF (top, based on **E**) or two TFs (bottom) between the frog and mouse thoracic spinal cords. More than two enriched populations for frog (black) or mouse (gray) are indicated.
**G**. A ventral population (**G''**) of V1 interneurons (En1, green) co-expresses the Renshaw cell markers MafB (red) and Calbindin (blue) in the NF54-55 frog spinal cord, quantified in **G'**. Shown is mean ± SEM for n = 4 animals
**H-Q**. Spatial plots showing the distribution of V1 expressing a given TF at the thoracic (**H-Q**) and lumbar levels (**K'-Q'**) in the frog (top row) and the mouse (bottom row).

**Figure S8. FoxP1 knockout affects motor neuron subtype but not V1 specification or motor neuron limb projections, related to Figure 7.**
**A-B. Generation of bilateral FoxP1 CRISPR mutant frogs.** Injection of FoxP1 sgRNA and Cas9 protein at one cell stage (**A**). Resulting mutants (**B** right) largely lacked FoxP1 (red) and Raldh2 (green) immunoreactivity as compared to wiltype (**B** left). Isl1/2-positive (blue) motor neurons were present in both conditions (**B**).
**C-D**. Quantification of bilateral FoxP1 mutants showed that $FoxP1^+$ (**C**) and $Raldh2^+$ (LMC, **D**) motor neurons were decreased at all spinal levels at NF54 ($p < 0.05$ for all levels except for Raldh2 at thoracic levels). Shown are mean ± SEM (n = 6 animals) per 15 µm ventral horn.
**E-F. Genomic characterization of unilateral and bilateral FoxP1 CRISPR mutant animals.** TIDE analysis reveals high efficiency of FoxP1 sgRNA in generating bilateral mutant animals at NF44-48 (**E**; WT vs mutant, p = 0.024, n = 3 for WT and n = 6 for FoxP1 animals), as well as unilateral mutant animals at juvenile stage (**F**; n = 2 for WT and n = 4 for FoxP1 animals).
**G-J. Profiling of other spinal neuron types in FoxP1 mutant.** Quantification of MMC ($Hb9^+Lhx3^+$, **G**) and V1 ($En1^+$, **H**) neurons on the mutant and uninjected side of the spinal cord

at all spinal levels. Shown are mean ± SEM (n = 3–6 animals) per 15 µm ventral horn. V1$^{1TF}$ (**I**) and V1$^{2TF}$ (**J**) subtypes are largely unaffected in unilateral FoxP1 mutants at NF54-55. Shown are mean ± SEM (n = 3–6 animals) per 15 µm ventral horn.
**K-L. Retrograde labeling with rhodamine dextran** (RhD, red) labels LMC motor neurons (FoxP1+, blue; Raldh2+, green; Isl1/2+, white) that project to the hindlimb in both wildtype (**L**) and unilateral FoxP1 CRISPR (**M**) mutant animals. Scale bar, 40 µm.

**Figure S9. Extended analysis of the effect of FoxP1 loss-of-function on limb- and tail-based locomotion, related to Figure 7.**
**A-F. Unilateral FoxP1 CRISPR mutant frogs display reduced locomotion at juvenile stage.** Trajectories of the distance traveled by an exemplary WT (**A** left) and unilateral FoxP1 CRISPR animals (**A** right) show different patterns of movement. FoxP1 mutants edge track as WT (**A** left), but move with less consistent direction (**A** right). Scale bar in **A** indicates the number of times the animal was present in a specific area of the dish from no time ($10^0$ frames, yellow) to many times ($10^3$ frames, blue). Quantification of overall movement of FoxP1 mutant animals shows that they move for less time (**B**; WT versus FoxP1 ½ CRISPR, p = 0.036) with shorter trajectory length (**C**; WT versus FoxP1 ½ CRISPR, p = 0.024) and less acceleration (**E**; WT versus FoxP1 ½ CRISPR, p = 0.002). Mutants however employ similar speed to WT (**D**). Unilateral FoxP1 CRISPR frogs also turn more than WT, with no difference between turning towards or away from the mutant side (**F**; for WT + versus FoxP1 ½ CRISPR +, WT + versus FoxP1 ½ CRISPR -, WT - versus FoxP1 ½ CRISPR + and for WT - versus FoxP1 ½ CRISPR +, p = <0.0001). n = 13 for WT, n = 14 for unilateral FoxP1 CRISPR.
**G-I. Loss of range, coordination and amount of movement along the rostrocaudal axis of the FoxP1 mutant hindlimb.** Quantification of mean angle of the mutant hip, ankle and foot show a different position than WT when moving (**G**. Hip, ankle and foot: WT L versus FoxP1 ½ CRISPR, WT R versus FoxP1 ½ CRISPR and uninjected versus FoxP1 ½ CRISPR, p = <0.0001). Left-right coordination between hip, ankle and foot joints is lost in FoxP1 CRISPR animals (**H**; +1 = bilateral synchronous, 0 = random, -1 = alternate synchronous. Hip, ankle and foot: WT versus FoxP1 ½ CRISPR, p = <0.0001). At the hip, ankle and foot joints, the amount of movement in the low frequency bin (0.9-4.5 Hz), represented by the sum power, is reduced for the FoxP1 CRISPR mutant limb compared to WT hindlimbs (**I**. Hip: WT L versus FoxP1 ½ CRISPR, p = 0.005; WT R versus FoxP1 ½ CRISPR, p = 0.065; uninjected versus FoxP1 ½ CRISPR, p = 0.032. Ankle: WT L versus FoxP1 ½ CRISPR, p = 0.002; WT R versus FoxP1 ½ CRISPR, p = 0.002. Foot: WT L versus FoxP1 ½ CRISPR, p = 0.013; WT R versus FoxP1 ½ CRISPR, p = 0.002; uninjected versus FoxP1 ½ CRISPR, p = 0.005) n = 13 for WT, n = 14 for unilateral FoxP1 CRISPR.
**J-P. Bilateral FoxP1 loss-of-function in NF44-48 does not affect tadpole locomotion.** Trajectories of the distance traveled by an exemplary WT (left) and bilateral FoxP1 CRISPR animals (right) show similar patterns of movement (**J top**). Scale bar in **J top** indicates the number of times the animal was present in a specific area of the dish from no time ($10^0$ frames, yellow) to many times ($10^3$ frames, blue). SLEAP skeleton (yellow) superimposed on WT (left) and bilateral FoxP1 CRISPR (right) tadpoles at NF44-48 with PCA plots representing the position of the tail and its range of movement during 256 random frames (**J bottom**; tail top, dark blue; tail mid, blue; tail tip, light blue). Scale bar in **J bottom** indicates the color-code of the first principal component of variation of the aligned tail positions. Bilateral FoxP1 CRISPR animals move for comparable time (**K**) and distance (**L**) to WT, employing similar speed (**M**), acceleration (**N**), and turning (**O**). Quantification of the range of the tail tip (**J**, light blue)

movement shows similar displacement to WT (**P**). n = 47 for WT, n = 39 for bilateral FoxP1 CRISPR.

**Figure S10. Extended analysis of the effect of En1 loss-of-function on tail and limb movement in tadpoles and frogs, related to Figure 8.**

**A-F. En1 mutant tadpoles have less and slower locomotion at NF44-48 with increased range of tail movement.** Trajectories of the distance traveled by an exemplary WT (**A**, left) and bilateral En1 CRISPR animals (**A**, right) show that mutant animals swim in circles and explore less of the dish (**A**). Scale bar in **A top** indicates the number of times the animal was present in a specific area of the dish from no time ($10^0$ frames, yellow) to many times ($10^3$ frames, blue). SLEAP skeleton (yellow) superimposed on WT (left) and bilateral En1 CRISPR (right) tadpoles at NF44-48 with PCA plots representing the position of the tail and its range of movement during 256 random frames (**A bottom**; tail top, dark blue; tail mid, blue; tail tip, light blue). Scale bar in **A bottom** indicates the color-code of the first principal component of variation of the aligned tail positions. Bilateral En1 CRISPR animals move for less time (**B**; WT versus En1 CRISPR, p = 0.022) with shorter distance traveled (**C**; WT versus En1 CRISPR, p = 0.0006) and employ slower speed (**D**; WT versus En1 CRISPR, p = <0.0001) and acceleration (**E**; WT versus En1 CRISPR, p = 0.002), while turning more than WT (**F**; WT versus En1 CRISPR, p = <0.0001). Quantification of the range of the tail tip (**A**, light blue) movement in bilateral En1 CRISPR mutant animals shows increased range in bilateral En1 CRISPR mutants compared to WT (**G**). n = 47 for WT and n = 37 for bilateral En1 CRISPR.

**H-L. Loss of high frequency and gain of low frequency movement in En1 CRISPR tadpoles at NF44-48.** Mean power spectrum of the tail tip oscillation shows a bimodal distribution for WT, with two peaks in the low and high frequency bin, and a unimodal distribution for bilateral En1 CRISPR animals, with only one peak in the low frequency bin for (**H**; low frequency bin, 0.9-4.5 Hz, dark gray; high frequency bin, 4.5-20 Hz, light gray). Bilateral En1 CRISPR mutant animals increase low frequency movement, gaining sum power in the low frequency bin (**I**; WT vs En1 CRISPR, p = <0.0001) and losing power in the high frequency bin (**K**; WT vs En1 CRISPR, p = 0.003). This loss is also captured by the flattening of the curve in the high frequency bin for the mutants (**H**). En1 CRISPR bilateral mutant tadpoles also have a decreased dominant low frequency (**J**; WT vs En1 CRISPR, p = 0.002) and no change in dominant high frequency (**L**). Notably, only a third of the bilateral En1 CRISPR mutant tadpoles even generate a dominant high frequency. n = 47 for WT and n = 37 for bilateral En1 CRISPR.

**M-R. Overall locomotion is not affected in En1 CRISPR mutant frogs.** Trajectories of the distance traveled by an exemplary WT (**M** left) and unilateral En1 CRISPR animals (**M** right) show similar patterns of movement with dish edge tracking. Scale bar in **M** indicates the number of times the animal was present in a specific area of the dish from no time ($10^0$ frames, yellow) to many times ($10^3$ frames, blue). Unilateral En1 CRISPR animals move for comparable time (**N**), distances (**O**) and employ similar speed (**P**) and acceleration (**Q**), while turning more than WT (**R**; WT versus En1 ½ CRISPR, p = 0.015). n = 13 for WT, n = 8 for unilateral En1 CRISPR.

**S-U. Neither range nor coordination of movement are affected in En1 mutant frogs.** While moving, the displacement of the hip, knee and ankle of unilateral En1 CRISPR mutant animals is comparable to WT animals (**S**). The range of movement of the hip and ankle is similarly unaffected; only the foot shows a higher range of displacement in En1 mutants compared to WT animals (**T**. Foot: WT versus En1 ½ CRISPR, p = <0.0001). Left-right coordination between hip, ankle and foot joints is also unaffected in unilateral En1 CRISPR

(**U**; +1 = synchronous, 0 = random, -1 = alternating). n = 13 for WT, n = 8 for unilateral En1 CRISPR.

**V-W. Lower dominant frequency in En1 CRISPR mutant frogs at all hindlimb joints.** At the hip, ankle and foot joints, the amount of movement, represented by the sum power, in the low frequency bin (0.9-4.5 Hz) is similar between WT and unilateral En1 CRISPR animals (**V**). However, the dominant frequency of the hip, ankle and foot is lower in mutants compared to WT animals (**W**. Hip: WT versus En1 ½ CRISPR, p = 0.039. Ankle: WT versus En1 ½ CRISPR, p = 0.043. Foot: WT versus En1 ½ CRISPR, p = 0.029). n = 13 for WT, n = 8 for unilateral En1 CRISPR.

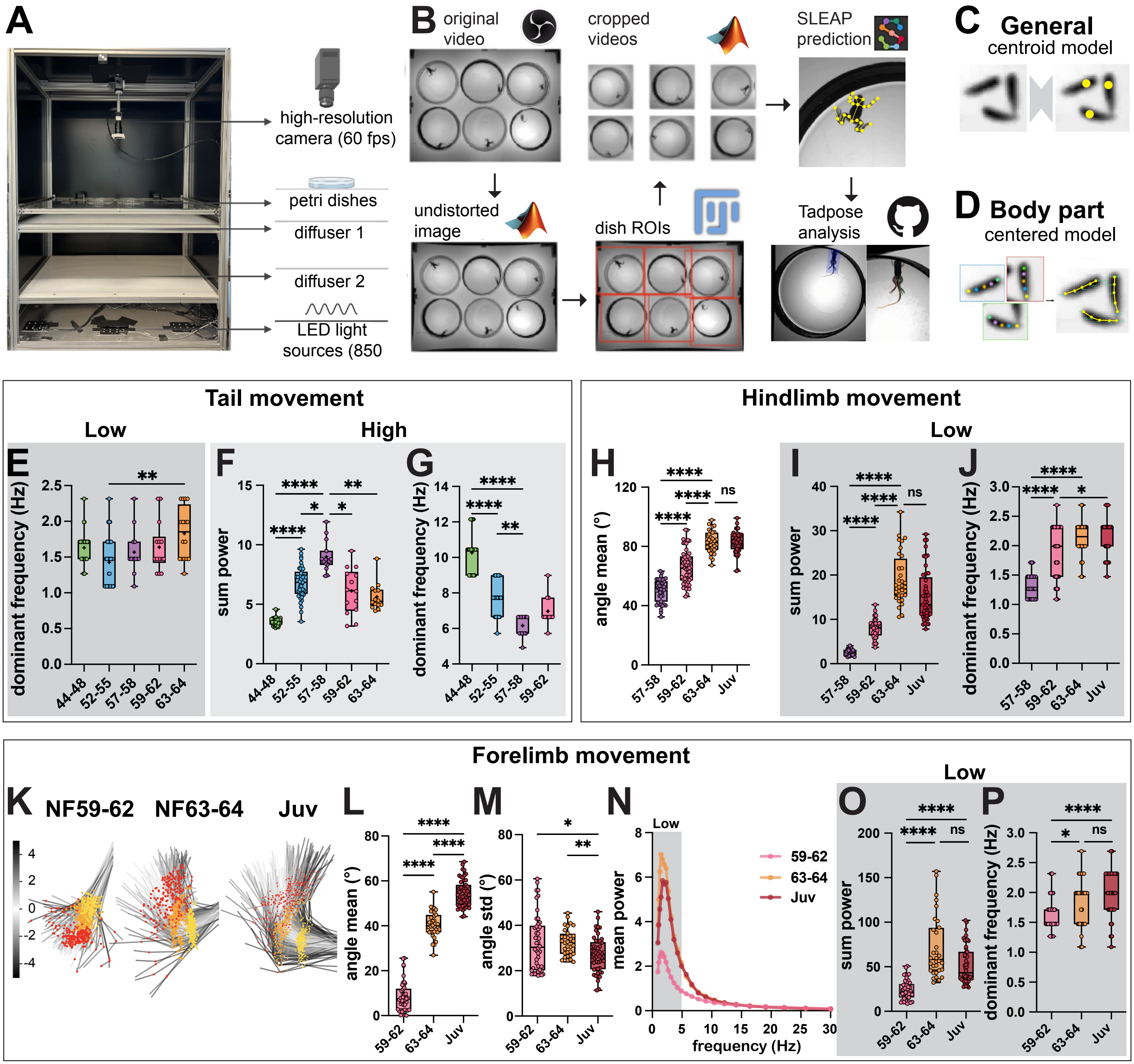
A
high-resolution camera (60 fps)
petri dishes
diffuser 1
diffuser 2
LED light sources (850
B
original video
cropped videos
SLEAP prediction
undistorted image
dish ROIs
Tadpose analysis
C
General
centroid model
D
Body part
centered model
Tail movement
Low
High
E
dominant frequency (Hz)
F
sum power
G
dominant frequency (Hz)
44-48
52-55
57-58
59-62
63-64
Hindlimb movement
Low
H
angle mean (°)
I
sum power
J
dominant frequency (Hz)
Juv
ns
Forelimb movement
K
NF59-62
NF63-64
Juv
L
angle mean (°)
M
angle std (°)
N
Low
mean power
frequency (Hz)
59-62
63-64
Juv
O
sum power
P
dominant frequency (Hz)

**Figure S1**

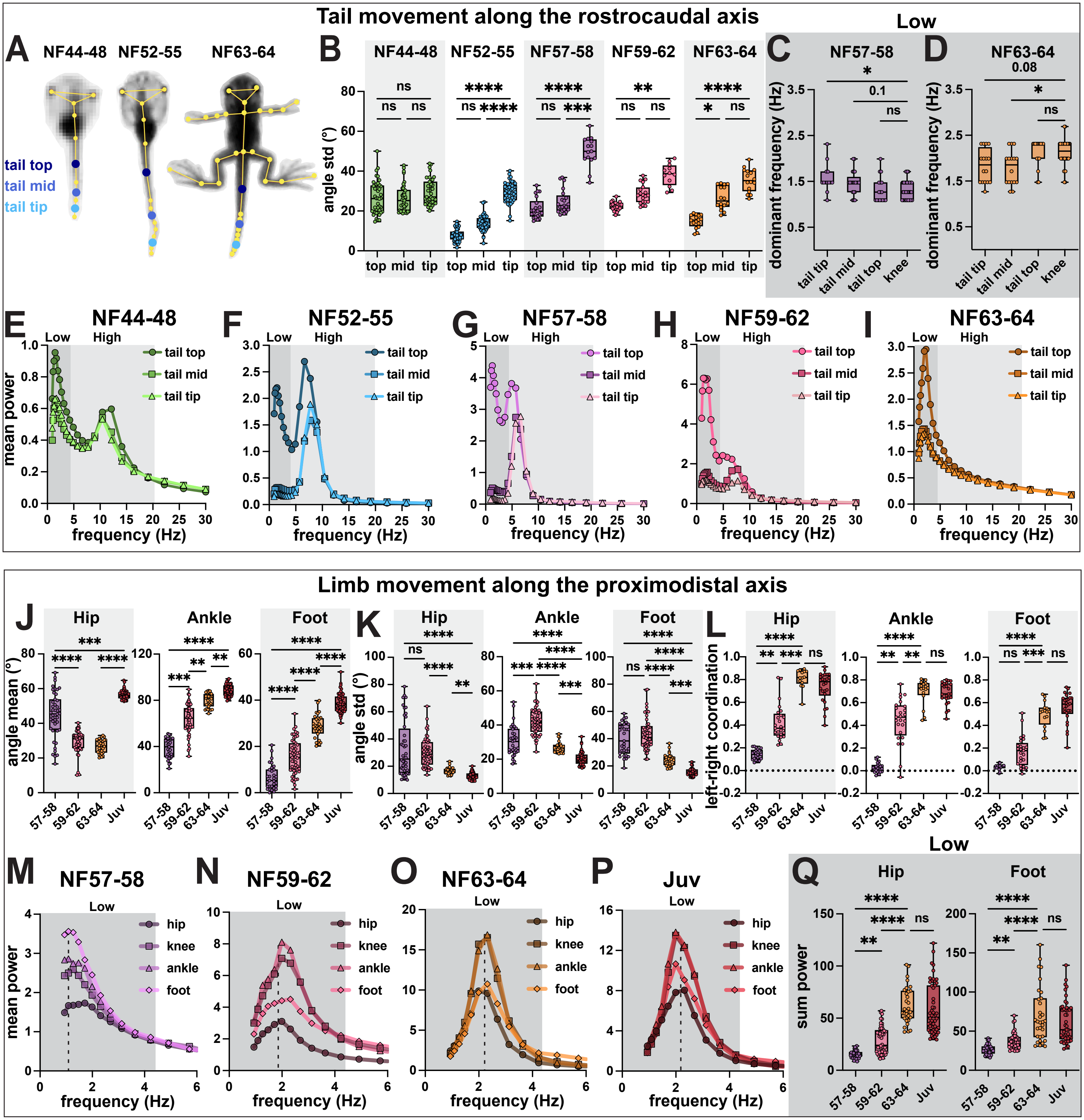
Tail movement along the rostrocaudal axis
A
NF44-48
NF52-55
NF63-64
tail top
tail mid
tail tip
B
angle std (°)
top mid tip
C
NF57-58
Low
dominant frequency (Hz)
tail tip
tail mid
tail top
knee
D
NF63-64
E
F
G
H
I
NF59-62
High
mean power
frequency (Hz)
Limb movement along the proximodistal axis
J
Hip
Ankle
Foot
angle mean (°)
57-58
59-62
63-64
Juv
K
L
left-right coordination
M
N
O
P
Juv
hip
knee
ankle
foot
Q
sum power

### Figure S2

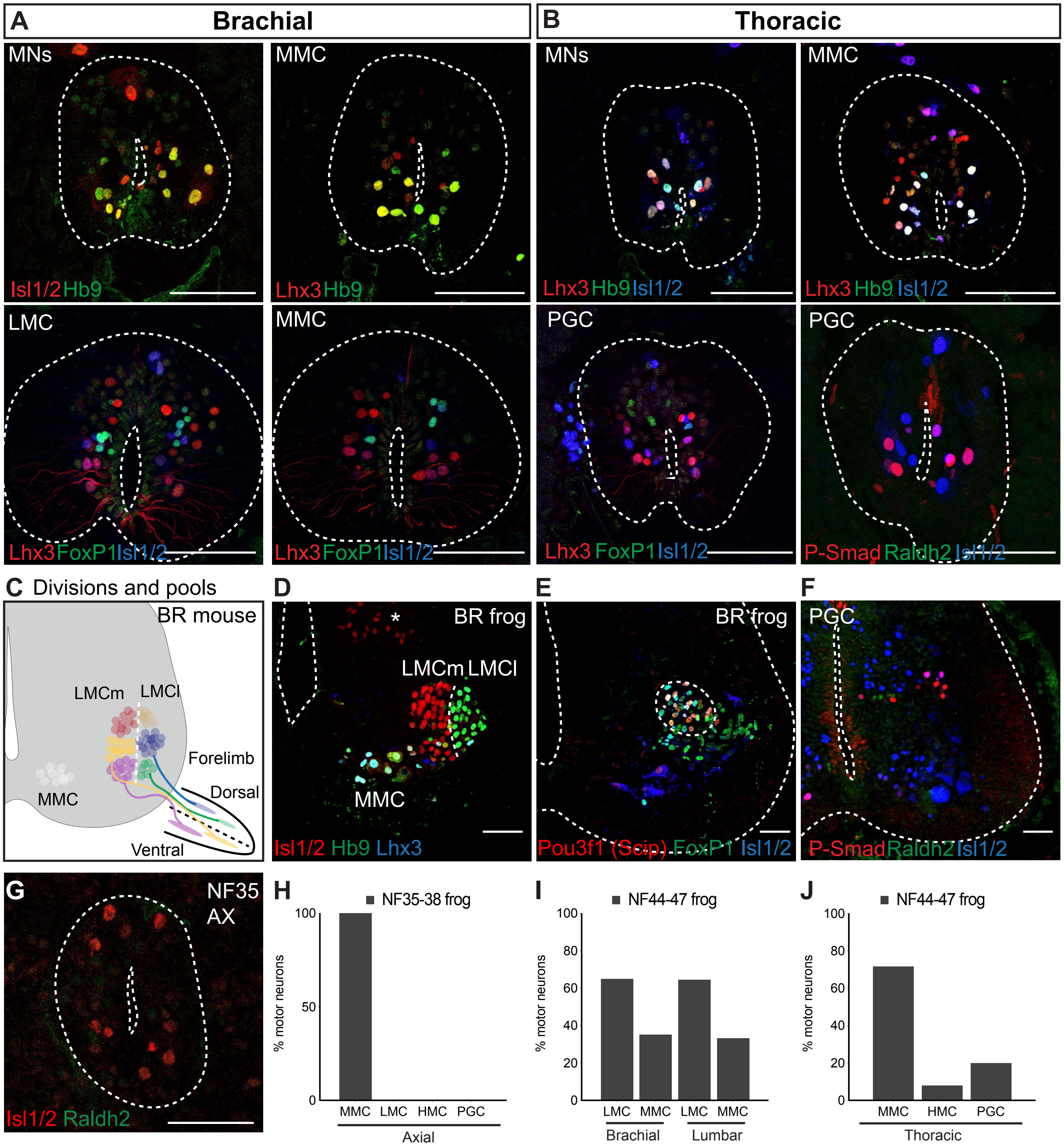
A
Brachial
B
Thoracic
MNs
Isl1/2 Hb9
MMC
Lhx3 Hb9
MNs
Lhx3 Hb9 Isl1/2
MMC
Lhx3 Hb9 Isl1/2
LMC
Lhx3 FoxP1 Isl1/2
MMC
Lhx3 FoxP1 Isl1/2
PGC
Lhx3 FoxP1 Isl1/2
PGC
P-Smad Raldh2 Isl1/2
C Divisions and pools
BR mouse
LMCm
LMCl
Forelimb
MMC
Dorsal
Ventral
D
BR frog
*
LMCm LMCl
MMC
Isl1/2 Hb9 Lhx3
E
BR frog
Pou3f1 (Scip) FoxP1 Isl1/2
F
PGC
P-Smad Raldh2 Isl1/2
G
NF35
AX
Isl1/2 Raldh2
H
NF35-38 frog
% motor neurons
100
50
0
MMC LMC HMC PGC
Axial
I
NF44-47 frog
% motor neurons
100
80
60
40
20
0
LMC MMC LMC MMC
Brachial
Lumbar
J
NF44-47 frog
% motor neurons
100
80
60
40
20
0
MMC HMC PGC
Thoracic

### Figure S3

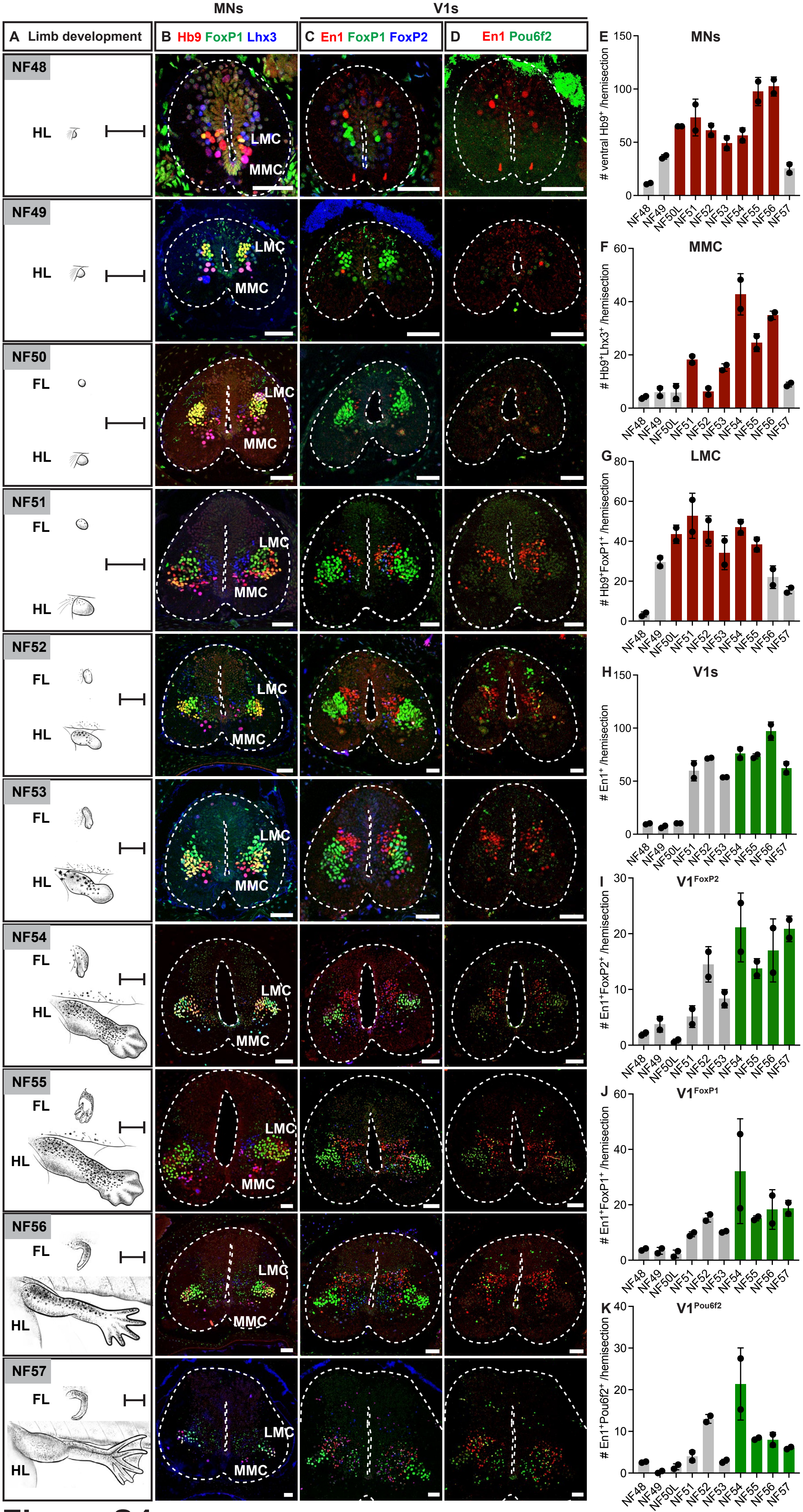

MNs
V1s
A Limb development
B Hb9 FoxP1 Lhx3
C En1 FoxP1 FoxP2
D En1 Pou6f2
NF48
NF49
NF50
NF51
NF52
NF53
NF54
NF55
NF56
NF57
FL
HL
LMC
MMC
E MNs
### ventral Hb9+ /hemisection
F MMC
### Hb9+Lhx3+ /hemisection
G LMC
### Hb9+FoxP1+ /hemisection
H V1s
### En1+ /hemisection
I V1FoxP2
### En1+FoxP2+ /hemisection
J V1FoxP1
### En1+FoxP1+ /hemisection
K V1Pou6f2
### En1+Pou6f2+ /hemisection
NF48 NF49 NF50L NF51 NF52 NF53 NF54 NF55 NF56 NF57


### Figure S4

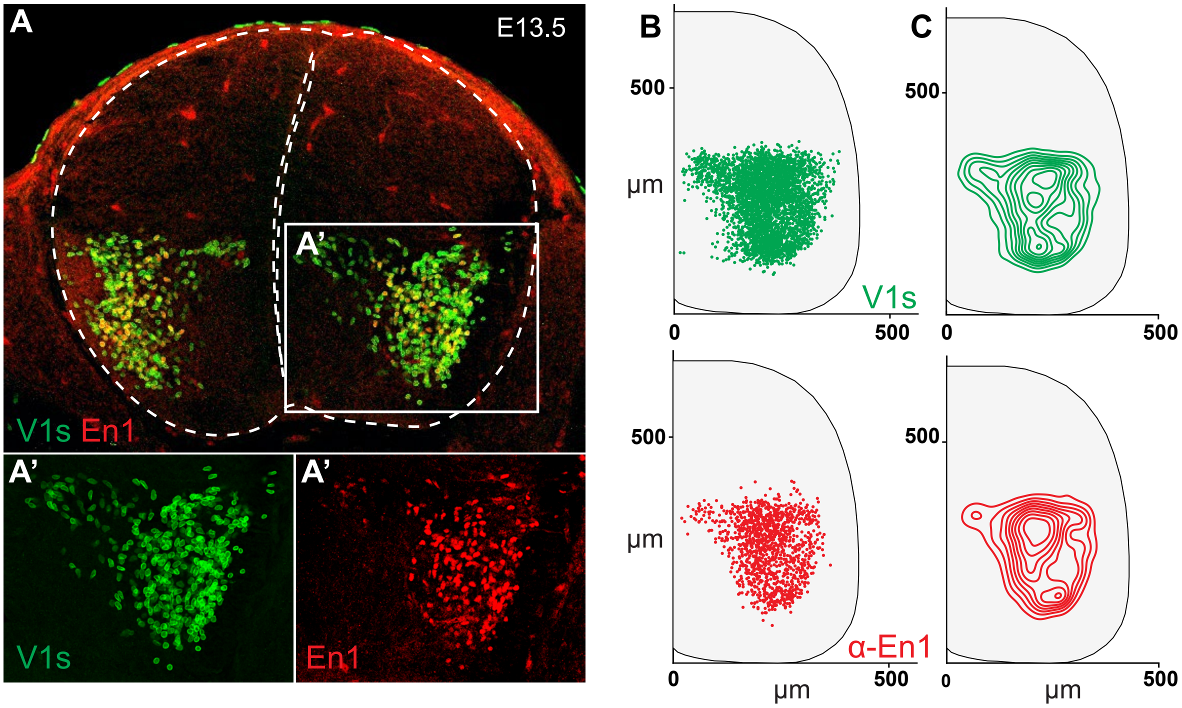
A
E13.5
A'
V1s En1
A'
A'
V1s
En1
B
500
µm
V1s
0
500
C
500
0
500
500
µm
α-En1
0
µm
500
500
0
µm
500

### Figure S5

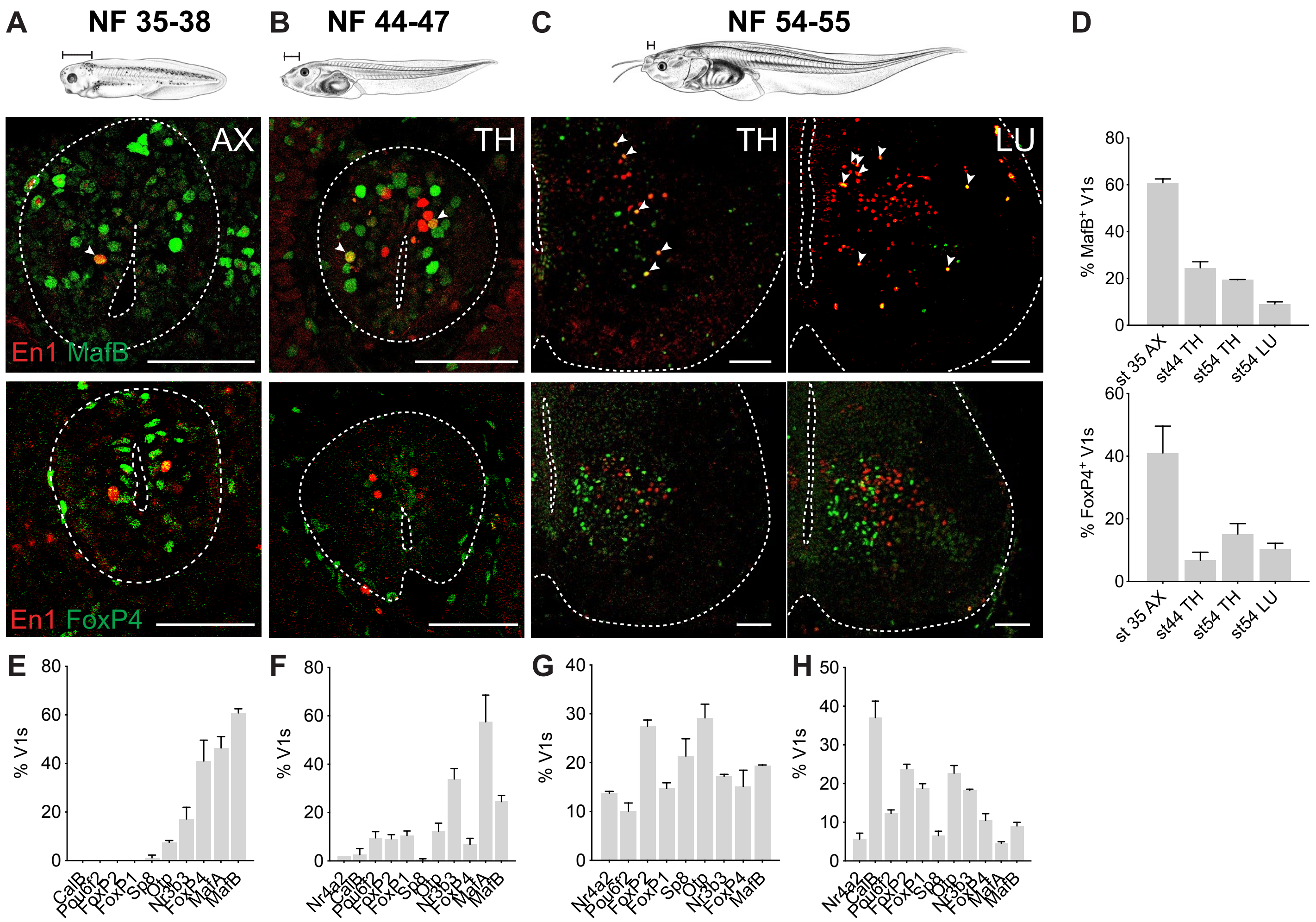
A
NF 35-38
B
NF 44-47
C
NF 54-55
D
AX
TH
TH
LU
En1 MafB
En1 FoxP4
% MafB+ V1s
st 35 AX
st44 TH
st54 TH
st54 LU
% FoxP4+ V1s
E
F
G
H
% V1s
CalB
Pou6f2
FoxP2
FoxP1
Sp8
Otp
Nr3b3
FoxP4
MafA
MafB
Nr4a2

### Figure S6

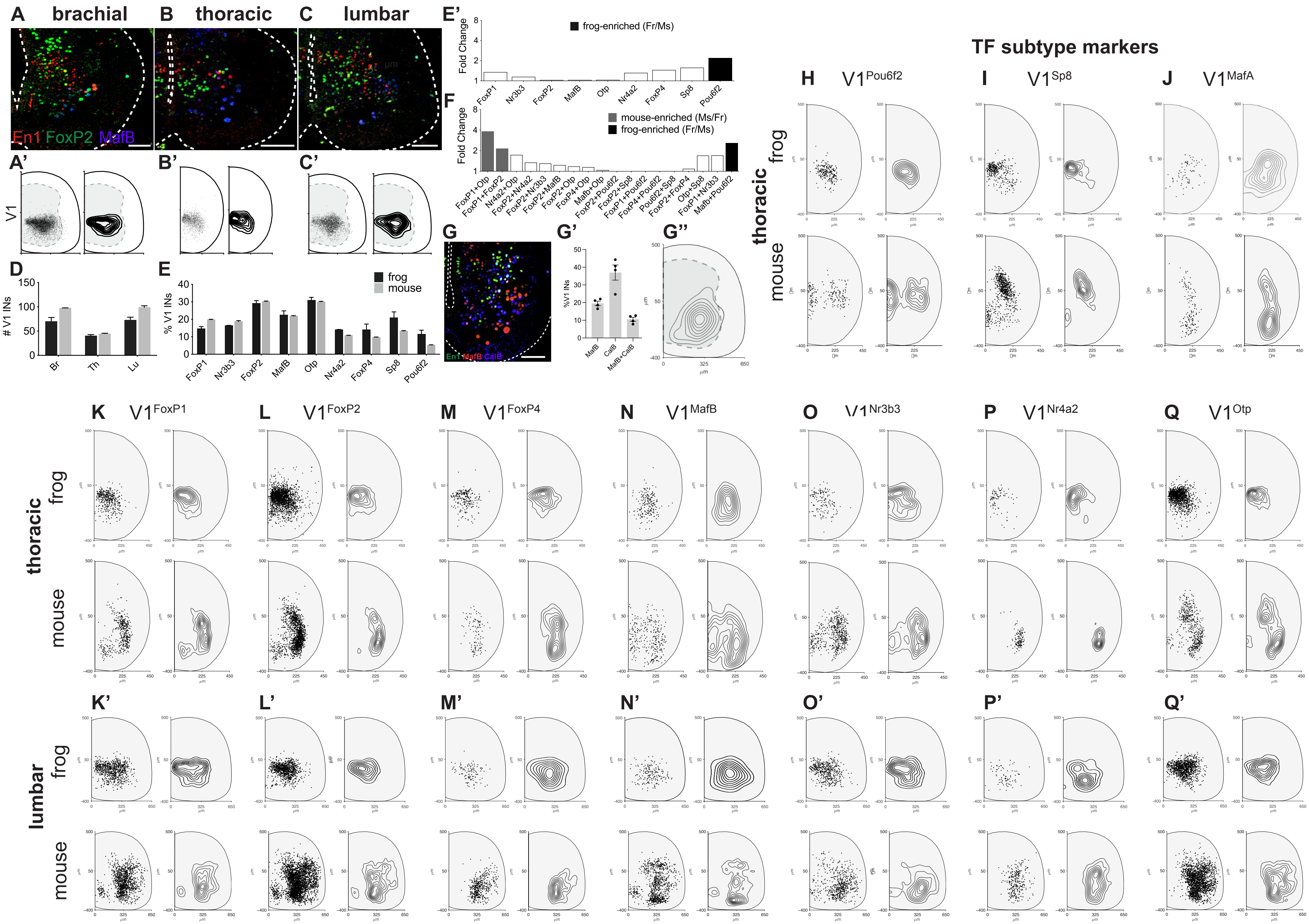

A brachial
B thoracic
C lumbar
En1 FoxP2 MafB
A'
B'
C'
V1
D
### V1 INs
Br
Th
Lu
E
% V1 INs
frog
mouse
FoxP1
Nr3b3
FoxP2
MafB
Otp
Nr4a2
FoxP4
Sp8
Pou6f2
E'
Fold Change
frog-enriched (Fr/Ms)
F
mouse-enriched (Ms/Fr)
frog-enriched (Fr/Ms)
FoxP1+Otp
FoxP1+FoxP2
Nr4a2+Otp
FoxP2+Nr4a2
FoxP2+Nr3b3
FoxP2+MafB
FoxP2+Otp
FoxP4+Otp
Mafb+Otp
FoxP2+Pou6f2
FoxP2+Sp8
FoxP1+Pou6f2
FoxP4+Pou6f2
Pou6f2+Sp8
FoxP2+FoxP4
Otp+Sp8
FoxP1+Nr3b3
Mafb+Pou6f2
G
En1 MafB CalB
G'
%V1 INs
MafB
CalB
MafB+CalB
G''
TF subtype markers
H V1Pou6f2
I V1Sp8
J V1MafA
thoracic
frog
mouse
K V1FoxP1
L V1FoxP2
M V1FoxP4
N V1MafB
O V1Nr3b3
P V1Nr4a2
Q V1Otp
K'
L'
M'
N'
O'
P'
Q'
lumbar


### Figure S7

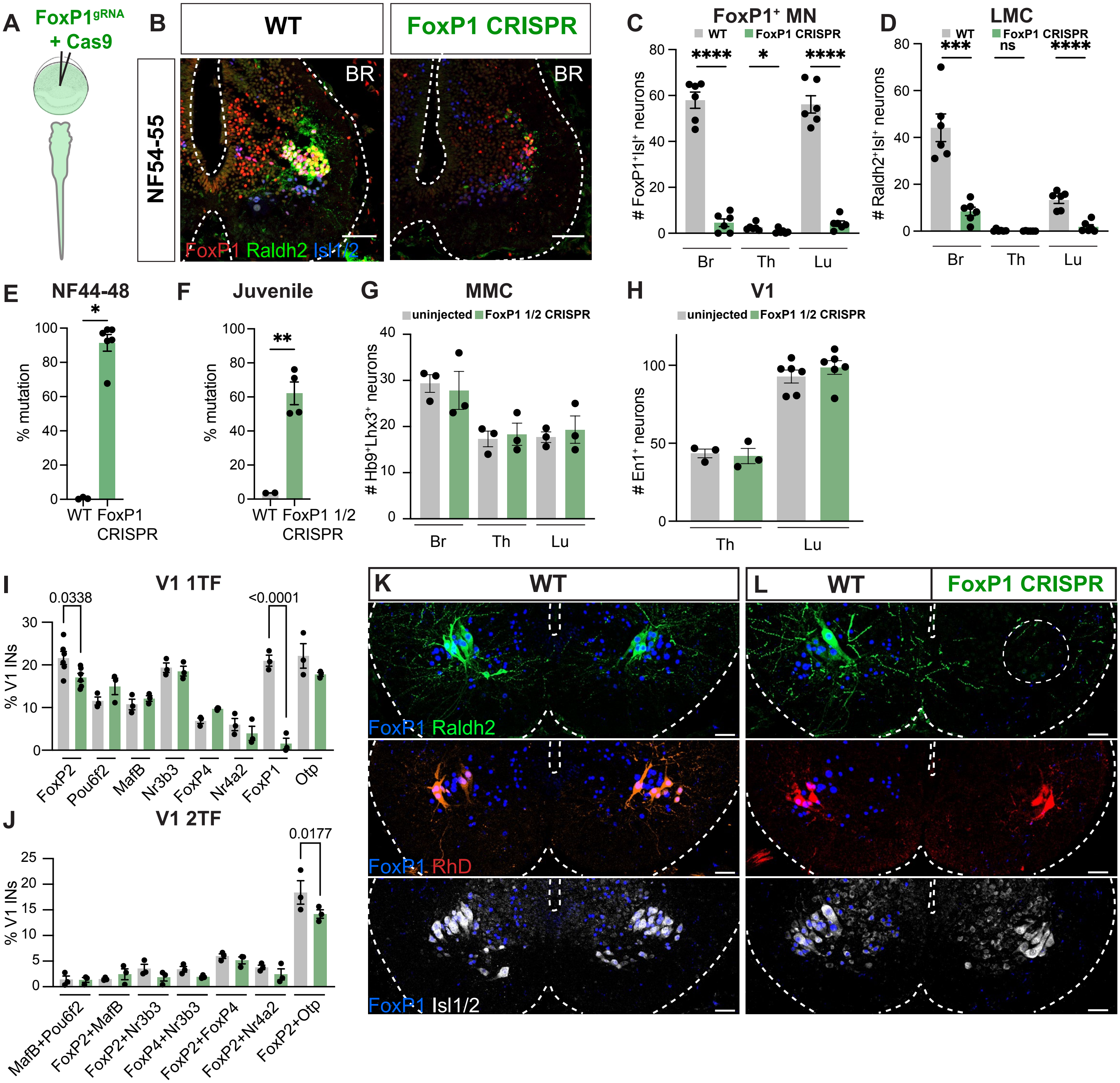
A
FoxP1gRNA
+ Cas9
B
WT
FoxP1 CRISPR
NF54-55
BR
FoxP1 Raldh2 Isl1/2
C
FoxP1+ MN
WT
FoxP1 CRISPR
### FoxP1+Isl+ neurons
Br
Th
Lu
D
LMC
### Raldh2+Isl+ neurons
E
NF44-48
% mutation
WT
FoxP1 CRISPR
F
Juvenile
FoxP1 1/2 CRISPR
G
MMC
uninjected
FoxP1 1/2 CRISPR
### Hb9+Lhx3+ neurons
H
V1
### En1+ neurons
I
V1 1TF
% V1 INs
0.0338
<0.0001
FoxP2
Pou6f2
MafB
Nr3b3
FoxP4
Nr4a2
FoxP1
Otp
J
V1 2TF
0.0177
MafB+Pou6f2
FoxP2+MafB
FoxP2+Nr3b3
FoxP4+Nr3b3
FoxP2+FoxP4
FoxP2+Nr4a2
FoxP2+Otp
K
WT
L
WT
FoxP1 CRISPR
FoxP1 Raldh2
FoxP1 RhD
FoxP1 Isl1/2

### Figure S8

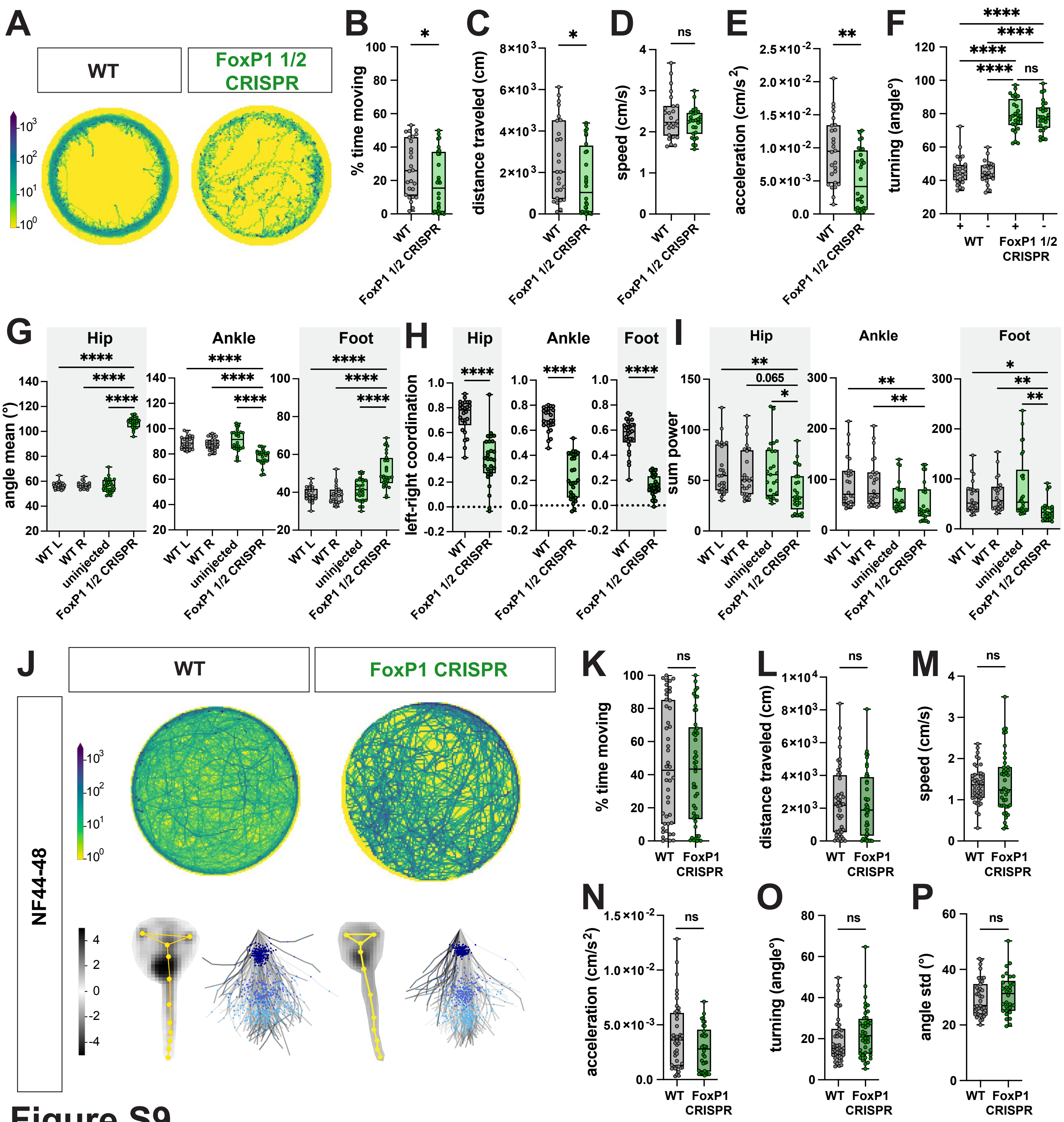

A
WT
FoxP1 1/2 CRISPR
B
% time moving
C
distance traveled (cm)
D
speed (cm/s)
E
acceleration (cm/s²)
F
turning (angle°)
WT
FoxP1 1/2 CRISPR
G
Hip
Ankle
Foot
angle mean (°)
WT L
WT R
uninjected
FoxP1 1/2 CRISPR
H
left-right coordination
I
sum power
J
WT
FoxP1 CRISPR
NF44-48
K
L
M
N
O
P
angle std (°)
FoxP1 CRISPR


Figure S9

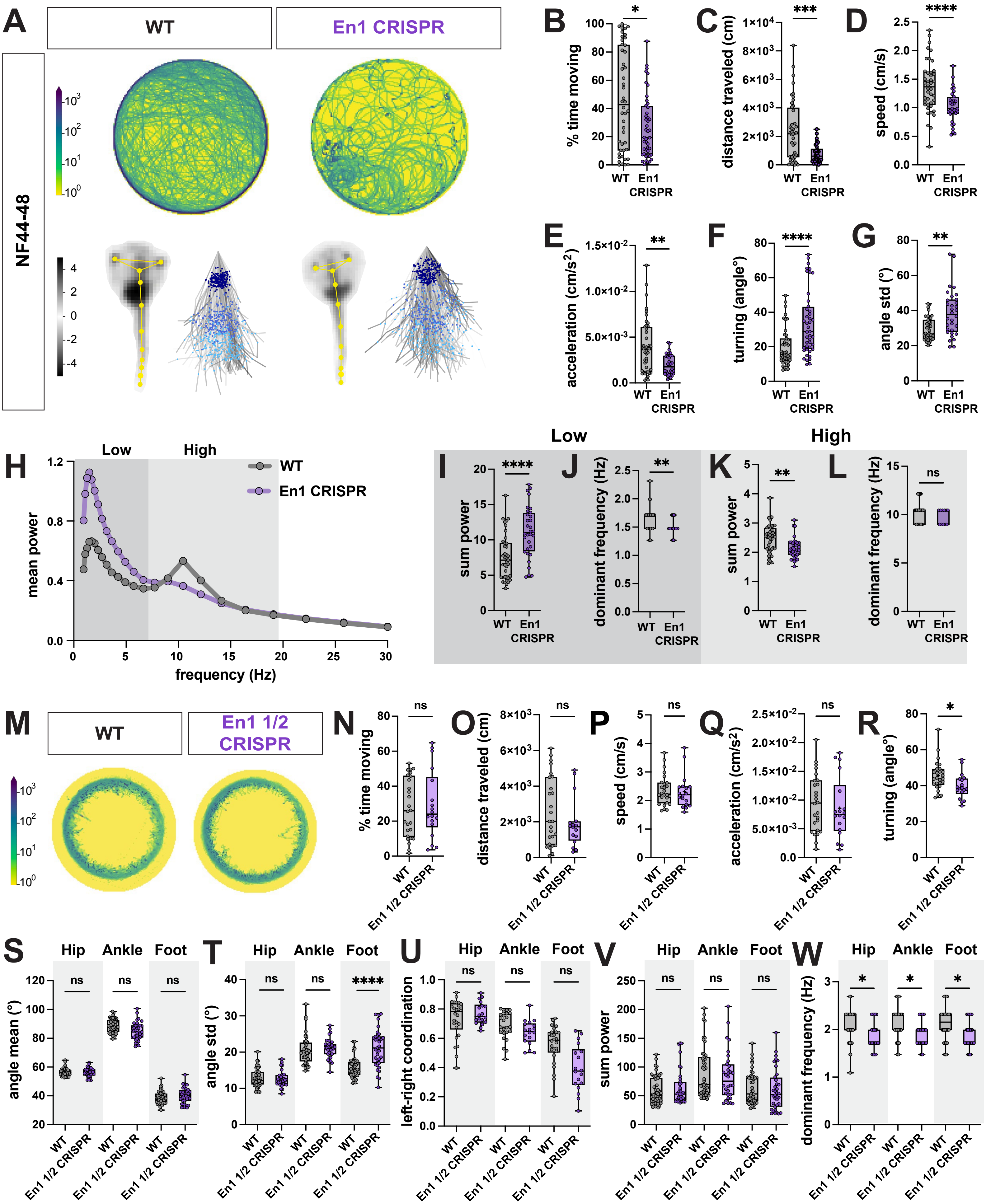

A
WT
En1 CRISPR
NF44-48
B
% time moving
C
distance traveled (cm)
D
speed (cm/s)
E
acceleration (cm/s2)
F
turning (angle°)
G
angle std (°)
H
Low
High
WT
En1 CRISPR
mean power
frequency (Hz)
I
sum power
J
dominant frequency (Hz)
K
sum power
L
dominant frequency (Hz)
M
WT
En1 1/2 CRISPR
N
% time moving
O
distance traveled (cm)
P
speed (cm/s)
Q
acceleration (cm/s2)
R
turning (angle°)
S
Hip
Ankle
Foot
angle mean (°)
T
angle std (°)
U
left-right coordination
V
sum power
W
dominant frequency (Hz)


Figure S10

| Number | Name | Sequence | Purpose | Source |
|---|---|---|---|---|
| SP533 | GFP_for | GCACGACTTCTTCAAGTCCGCCAT | Genotyping | IDT |
| SP534 | GFP_rev2 | GCGGATCTTGAAGTTCACCTTGAT | Genotyping | IDT |
| SP154 | FoxP1_gRNA1_for | GGAGTGAGGGCATCATTTGT | Genotyping | IDT |
| SP155 | FoxP1_gRNA1_rev | GGTAAGGGAGGGAGGTAACG | Genotyping/TIDE | IDT |
| SP01 | En1_gRNA4_for | GGAGAATTGTTGGAAGGGAGA | Genotyping | IDT |
| SP02 | En1_gRNA4_rev | AGTAGATTTCCCTGTTGCTGG | Genotyping/TIDE | IDT |

**Table S1**: Primer sequences used for PCR genotyping and TIDE analysis.

| Target gene | Name | Sequence | Purpose | Source |
|---|---|---|---|---|
| FoxP1 | FoxP1_gRNA1 | ACTTGTGTAGACTAAGATTATGG | generating bilateral and unilateral mutants | Synthego |
| En1 | En1_gRNA4 | GCTCAGCAGTGAAGGCTGTTCGG | generating bilateral and unilateral mutants | Synthego |

**Table S2**: Guide RNA sequences used to generate CRISPR mutants.

| Metrics | | | | | | |
|---|---|---|---|---|---|---|
| **Model** | **Stage bin** | **Average distance** | **Visual recall** | **Visual Precision** | **OKS** | **mPCK** |
| Centroid model | NF37-38 | 1.03709 | 1 | 1 | 0.94186 | 0.94395 |
| | NF44-48 | 6.95001 | 1 | 1 | 0.54758 | 0.8125 |
| | NF52-55 | 0.00001 | 1 | 1 | 0.92079 | 0.85714 |
| | NF57-58 | 0.00001 | 1 | 1 | 0.47525 | 1 |
| | NF59-62 | 0.00001 | 1 | 1 | 0.32673 | 1 |
| | NF63-64 | 0.00001 | 1 | 1 | 0.76238 | 1 |
| | Juvenile | 0.00001 | 1 | 1 | 0.62376 | 1 |
| Centered model | NF37-38 | 2.97621 | 0.98443 | 0.99751 | 0.34501 | 0.86163 |
| | NF44-48 | 6.1345 | 0.93077 | 0.97059 | 0.10264 | 0.71244 |
| | NF52-55 | 3.44924 | 0.93722 | 0.99524 | 0.33574 | 0.68918 |
| | NF57-58 | 3.17561 | 0.94041 | 0.95863 | 0.5896 | 0.66311 |
| | NF59-62 | 2.6415 | 0.99508 | 0.95997 | 0.77718 | 0.68521 |
| | NF63-64 | 1.71405 | 0.99754 | 0.98901 | 0.81273 | 0.87478 |
| | Juvenile | 2.14075 | 0.99657 | 0.97865 | 0.88478 | 0.81674 |

**Table S3**: Metrics of the behavioral tracking models.

| Gene to amplify | Step | PCR temperature | Time | Cycles |
|---|---|---|---|---|
| **FoxP1** | Initial denaturation | 98 °C | 30s | 1 |
| | Denaturation | 98 °C | 5s | 32 |
| | Annealing | 57.7 - 63.3°C | 5s | |
| | Extension | 72 °C | 10s | |
| | Final extension | 72 °C | 60s | 1 |
| | Storage | 12 °C | ∞ | Storage |
| **En1** | Initial denaturation | 98 °C | 30s | 1 |
| | Denaturation | 98 °C | 5s | 32 |
| | Annealing | 57.3 - 63.3°C | 5s | |
| | Extension | 72 °C | 10s | |
| | Final extension | 72 °C | 60s | 1 |
| | Storage | 12 °C | ∞ | Storage |

**Table S4**: PCR conditions used for genotyping FoxP1 and En1 genes.
